# Promoting long-term forest landscape resilience in the Lake Tahoe basin

**DOI:** 10.1101/2021.06.29.450434

**Authors:** Eric S. Abelson, Keith M. Reynolds, Angela M. White, Jonathan W. Long, Charles Maxwell, Patricia N. Manley

## Abstract

Rapid environmental changes expected in the 21^st^ century challenge the resilience of wildlands around the world. The western portion of the Lake Tahoe basin (LTW) in California is an important ecological and cultural hotspot that is at risk of degradation from current and future environmental pressures. Historical uses, fire suppression, and a changing climate have created forest landscape conditions at risk of drought stress, destructive fire, and loss of habitat diversity. We prospectively modeled forest landscape conditions for a period of 100 years to evaluate the efficacy of five unique management scenarios in achieving desired landscape conditions across the 23,600 hectares of LTW. Management scenarios ranged from no management other than fire suppression to applying treatments consistent with historical fire frequencies and extent (i.e., regular and broadscale biomass reduction). We developed a decision support tool to evaluate environmental and social outcomes within a single framework to provide a transparent set of costs and benefits; results illuminated underlying mechanisms of forest resilience and provided actionable guidance to decision makers. Sixteen attributes were assessed in the model after assigning weights to each, derived through a survey of stakeholder priorities, so that the contribution of each attribute to evaluations of scenario performance was influenced by the combined priorities of stakeholders. We found that removing forest biomass across the landscape, particularly when accomplished using extensive fire-based removal techniques, led to highly favorable conditions for environmental quality and promoting overall landscape resilience. Environmental conditions resulting from extensive fire-based biomass removal also had nominal variation over time, in contrast with strategies that had less extensive and/or used physical removal techniques, namely thinning. Our analysis provided a transparent approach to data assessment, considering the priorities of stakeholders, to provide insights into the complexities of maintaining optimal conditions and managing landscapes to promote ecosystem resilience in a changing world.

## INTRODUCTION

As society faces a changing climate and the uncertainty that it entails, nearly all sectors (government and private) are working feverishly to understand and mitigate threats to ecological integrity (*sensu* Cleland et al. 2017) and the persistence of ecosystem function and services (Daily and Matson 2008). The concept of resilience offers a tangible beacon for a future that inevitably will be different, but that also can be ecologically diverse, productive, aesthetic, and meet the needs and desires of society in a sustainable manner. Resilience is the “capacity of a system to absorb disturbance and reorganize while undergoing change so as to still retain essentially the same function, structure, identity, and feedbacks” (Walker et al. 2004). In the ecological literature, outcomes are typically understood as characterizing the *state* of a system and such characterizations are therefore also commonly understood as evaluating ecosystem integrity, as opposed to ecosystem resilience, for which the focus is on the response of systems to perturbations as a function of ecosystem *processes* (e.g., Holling 1973, Folke et al. 2002, Gunderson et al. 2010, Walker et al. 2010). As such, resilience only can be truly evaluated after perturbations have occurred, the system has responded, and the outcomes are evaluated.

Lake Tahoe and its basin are rich with biodiversity, aesthetic values (e.g., scenic beauty), and recreational opportunities, that are particularly vulnerable to impacts from climate change. A recent vulnerability analysis commissioned by the California Tahoe Conservancy (California Tahoe Conservancy 2020), and supported by empirical studies (e.g., Coats et al. 2006, Scheller et al. 2018), concluded that both the Lake and the upland watersheds are not only vulnerable to changing climates, but are already exhibiting signs of stress and impact. The future of ecological integrity in the Lake Tahoe basin is a core concern across a wide array of stakeholders, including scientists and managers operating in the basin (Chilton 1995, Imperial and Kauneckis 2003, Weible et al. 2005).

Attention paid to Lake Tahoe is largely directed toward the ecological and social value of the basin, but it is also a bellwether for much of the Sierra Nevada mountain range; investments in science and management in the basin are significant compared to most other landscapes in the Sierra, so science-based management solutions are more likely emerge there first (Murphy and Knopp 2000, Manley 2004, Hymanson et al. 2010). The Lake Tahoe West Restoration Partnership (LTWRP) is a prime example of a science-based management solution, formed to develop and implement a large-scale landscape restoration strategy across the west side of the Lake Tahoe basin, which ideally could serve as a model for restoration across the basin. The goal of the Lake Tahoe West (LTW) project is to restore the resilience of the forests, watersheds, and communities of the west side of the Lake Tahoe basin to disturbances including wildfire, drought, and climate change. The LTWRP is a multi-stakeholder collaborative initiative lead by the California Tahoe Conservancy, U.S. Forest Service Lake Tahoe Basin Management Unit, California State Parks, Tahoe Regional Planning Agency, and the National Forest Foundation.

Typically, the first step in developing a large-scale restoration strategy is to assess current and potential future conditions, and compare those to desired conditions (e.g., Stanturf et al. 2013). Given the focus of decision makers on sustainability and resilience across multiple social and ecological values, a focus of this study was to evaluate future trajectories under different management scenarios in the face of climate change (Hymanson et al. 2010). Large-scale ecological assessments have many moving parts that make defining the analytical problem, assembling large volumes of data, and evaluating and distilling results challenging. Decision support tools (DSTs) are increasingly used to help managers understand the spatial and temporal variability in conditions across multiple resources, and how best to balance management objectives where they may be in conflict or at least difficult to achieve in an equitable manner (e.g., Bagstad et al. 2013, Vacik et al. 2013, Gordon and Reynolds 2014). DSTs are valuable not only to managers, but they support the entire decision process, which often includes a broad array of engaged stakeholders. DSTs can provide transparency, credibility, trust, and confidence in the decision process, which are critical to successful implementation of management decisions (Reynolds et al. 2014a).

Many DSTs applied in forest landscape restoration projects are designed to identify potential project areas and quantify the relative risks and benefits associated with various treatment approaches (Hessburg et al. 2013, Povak et al. 2017). Prioritization of treatment areas is often the objective, with the near-term goal of reducing risk to high value resources. In this study, we instead focused on identifying the ecological, social and operational underpinnings of management activities for improving forest landscape resilience over a long time-horizon. To this end, we evaluated management regimes (in the form of scenarios) conducted over multiple decades across the entire landscape to determine the degree to which landscape resilience had been improved and by what measures.

We developed a DST that provided a unified modeling framework for assembling and evaluating multiple resource conditions over a 100-year timeframe to determine if and how management could promote greater forest landscape resilience across the LTWRP landscape into the future. Three essential tenets of effective decision support solutions (Reynolds and Hessburg 2014, Reynolds et al. 2014b) guided the development of the LTW DST tool and its application: 1) the tool needs to facilitate collaboration among multiple and diverse subject-matter experts and stakeholders; 2) the design of decision models for decision makers needs to formally incorporate the diversity of decision maker perspectives; and 3) the analytical models used need to yield results that are intuitive and accessible to broad stakeholder communities.

## METHODS

### Study area

The Lake Tahoe basin is an 88,000 hectare lake basin that rests between the Sierra Nevada range to the west and the Carson Range to the east along the California and Nevada border, respectively. Lake Tahoe itself sits at 1880 m in elevation, with the surrounding landscape consisting of over 60 forested watersheds that stretch from the lake shore up to the crests reaching over 3000 m. The LTW landscape consists of the 22 watersheds and approximately 23,600 hectares (59,000 acres) of federal, state, local, and private lands on the western side of the basin from the crest to the lake. The landscape consists of extensive forests and many creeks that course through glaciated valleys and wet meadows before draining into the lake. The forests on the west side of Lake Tahoe have experienced a high degree of disturbance over the past 100 years. Although they have not experienced the large and severe wildfires that have impacted much of the Sierra Nevada in recent decades (Lydersen et al. 2014, Jones et al. 2016), there have been two fires in the past 15 years: the 2007 Angora Fire that burned over 1200 hectares in the southern part of the basin causing enormous loss of property (Safford et al. 2009), and the 2016 Emerald Fire that burned just over 70 hectares near Emerald Bay, CA. In addition, the basin regularly experiences significant smoke events from fires outside the basin. The basin was extensively logged, beginning in the 1860s during the Comstock silver mining rush in Virginia City in Nevada, and continuing into the 20^th^ century (Lindström 2000), creating a predominantly single cohort of trees, the survivors of which are greater than 100 years old. In recent decades, management has focused on restoration rather than the production of timber, livestock grazing and other consumptive uses.

### Management scenarios

We developed criteria for a range of management approaches to evaluate the effectiveness of the extent, distribution, and type of management treatments for achieving desired conditions. Specifically, management scenarios were developed that addressed three key management options to restore forest landscape resilience: 1) protecting infrastructure, primarily within the wildland urban interface (WUI), versus managing across the entire landscape, including “back country” wilderness areas; 2) forest biomass reduction by thinning versus the use of prescribed fire and managed wildland fire; and 3) management and societal costs and benefits relative to environmental costs and benefits. The result was five management scenarios that ranged from no management, other than fire suppression, to landscape-wide management with various combinations of thinning and prescribed fire (Table 1). This approach reflected decisionmakers’ interests in understanding how increases in the pace and scale of treatment would enhance the resilience of Lake Tahoe’s social-ecological system (North et al. 2012, Stephens et al. 2014, Manley et al. 2021), and enhance their understanding of inherent tradeoffs associated with management approaches in meeting resource objectives. Specific evaluation criteria and thresholds were developed through collaborations between members of the LTWRP science team (the scientists that conducted the environmental modeling represented here), managers from multiple agencies operating within the basin, and stakeholders representing a range of interests.

**Table 1.** Management scenarios evaluated using Ecosystem Management Decision Support tool for the Lake Tahoe West restoration landscape.

| Scenario number | Description | Management specifications |
| --- | --- | --- |
| 1 | Fire suppression only | The only management activity was to suppress fires that start naturally. No other management activities were implemented in this scenario. |
| 2 | Wildland-urban interface focus | Management activities were focused predominantly on forest thinning in the wildland-urban interface (WUI, areas near human habitation). This management strategy was designed to provide a buffer of defensible space around human-built structures and property, with the goal of protecting those properties and their inhabitants. It treated approximately 1.8% of the vegetated area each year, all in the WUI (58% of the landscape). This scenario most closely resembled current management activities in the Lake Tahoe basin. |
| 3 | Thinning-based approach | This scenario builds upon Scenario 2 by expanding management activities into the remaining forested landscape (42% of the landscape) in proximity to the WUI and used predominantly mechanical and some hand removal methods to thin the forest and reduce woody biomass. It treats approximately 6.7% of the vegetated area each year. |
| 4 | Fire-based approach | This scenario builds upon Scenario 2 by expanding management activities into the remaining forested landscape. While Scenario 3 employs mechanical and hand methods to thin the forest, Scenario 4 uses primarily prescribed fire and managed wildfire. This scenario treats approximately 4% of the vegetated area each year. |
| 5 | Extensive fire-based approach | This scenario builds upon Scenario 2 by expanding management activities into the remaining forested landscape. Like Scenario 4, Scenario 5 predominately employs fire-driven techniques, but with a greatly expanded the use of prescribed fire. This scenario treats a approximately 7.2% of the vegetated area each year, slightly more than Scenario 3, but with the majority of treatments (75%) being fire. |

### Assessing landscape condition

Decision support tools not only support management decisions once data are compiled, but they also provide a framework for organizing information in the initial stages of model building. Thus, an important contribution of our DST to the LTWRP was to provide structure and a process to clarify what resources were most relevant to managers, what conditions were relevant and desired for those resources, and what metrics and values best represented those desired conditions. Resources essential to understanding forest resilience in response to management scenarios were determined by environmental experts, decision makers, and project stakeholders, and included not only resources pertaining to environmental quality (terrestrial and aquatic ecological conditions), but also community values and management operations (Figure 1), which can both drive and limit implementation.

**Figure 1.**
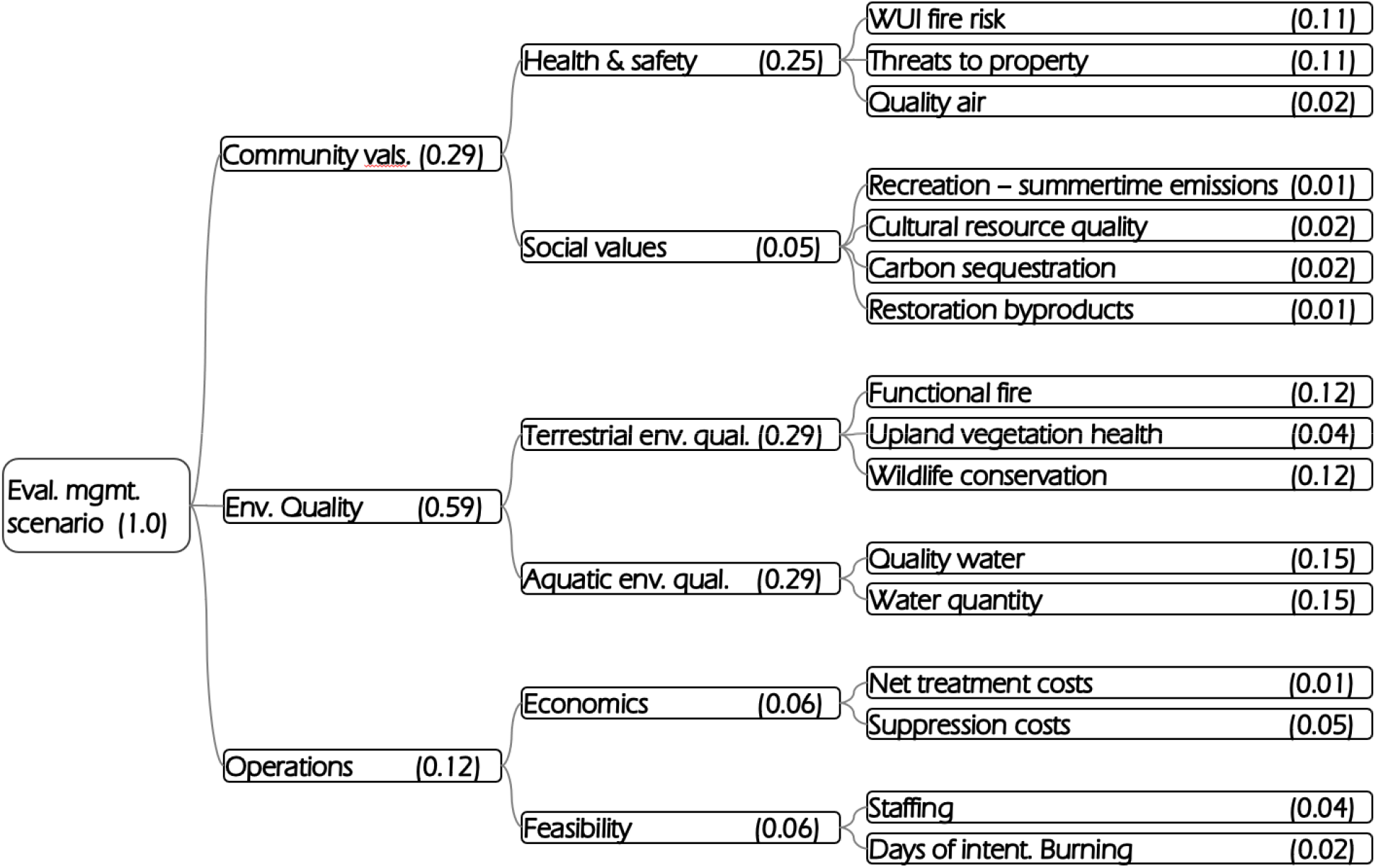
Decision model hierarchy to address the performance of management scenarios in achieving desired conditions across three focal areas: environmental quality, community values, and management operations. Each focal area is represented by two topic areas and each topic is represented by two to four attributes, for a total of 16 attributes. Parenthetical values indicate weighting derived by stakeholders (rounded to two decimals for display purposes), with assigned weights summing to one for each tier of the hierarchy.

For each of the three focal areas (environmental quality, community values, and management operations), quantifiable metrics were identified based on available, relevant, and peer-reviewed data sources. The type and number of attributes varied among the three focal areas based on scientific merit (environmental quality), stakeholder priorities (community values), and management considerations (operations). Each focal area is represented by two tiers: topic areas and attributes. A total of 16 attributes were established across the three focal areas (Figure 1).

The 16 individual attributes were in turn represented by one or more metrics of condition. The number of metrics selected to represent each attribute varied based on the complexity of the attribute (see Miller and Saunders 2002, Saunders and Miller 2014), with those associated with environmental quality having the largest suite of metrics, and those associated with community values and operations typically being represented by single metrics (Appendices 1, 2, and 3; see Abelson et al. *in review* for detailed methods). For example, the quality water topic was represented by two metrics (phosphorus load and fine sediment), whereas biodiversity conservation was represented by 13 metrics representing three focal species of interest, six functional species groups, and four measures of species diversity (Appendices 2 and 3). Subject-matter experts identified the data that was necessary to evaluate attributes and determined the metric values that corresponded with poor to optimal conditions (Appendices 1 and 4). These values were used in the DST to evaluate forest conditions, propagating from the bottom of the hierarchy (attributes) to the top (resilience), for each incoming piece of data (Appendix 5) following methods described in Abelson et al. (in review).

### Forest, fire, and climate dynamics

LANDIS-II modeling was used to simulate management activities and natural processes, and their interaction, over a 100-year period (2010 to 2110) across the entire Lake Tahoe Basin landscape (see Maxwell et al. 2021 this series). In short, above-ground forest dynamics along with nutrient flow, pest outbreaks and climate variability are integrated into a single process-based simulation (Scheller et al. 2019). Five unique LANDIS-II models were run with all model variables being held constant except management activity. The LANDIS-II modeling that provided inputs to the DST was based on a representative concentration pathway (RCP) of 4.5, and averaged outputs from four different global circulation models (GCMs) identified as the most representative of California’s hydrology in California’s Fourth Climate Change Assessment (Pierce et al. 2018) served as the basis for the future climate projections used in this study: Hadley Center Global Environment Model (HadGEM2), Canadian Earth System Model (CanESM), Centre National de Recherches Meteorologiques (CNRM5), and Model for Interdisciplinary Research on Climate (MIROC5). The outputs of the LANDIS-II modeling for the LTW landscape were then used by topic-area specialists to quantify the metrics used to represent each of the 16 attributes in our DST to evaluate scenario performance (Table 2). In an important sense, the evaluation conducted in this study reflects both integrity and resilience because, although we evaluated management performance in terms of outcomes, most of the model inputs were derived from the process-based LANDIS-II system, which dynamically models the evolution of system states over time based on the simulation of processes (Scheller et al. 2019).

**Table 2.** Attributes of the three focal areas evaluated using the Ecosystem Management Decision Support tool to determine the degree to which five different management scenarios met desired conditions for the Lake Tahoe West landscape.

| Attributes of focal areas | Modeling category description |
| --- | --- |
| *Environmental quality:* |  |
| Functional fire | Measures how close to the natural range of variability fires are predicted to burn at low, moderate, and high severity. |
| Upland vegetation health | Considers to what extent early, mid, and late seral forests are represented across the landscape compared to modern reference conditions. |
| Wildlife conservation | Represents species richness, biodiversity across multiple functional groups, and the quality and connectivity of old-growth associated species habitat. |
| Quality water | Represents fine sediment and nutrient loading to streams and lakes compared to baseline conditions. |
| Water quantity and timing | A qualitative measure of increased water yield and delayed runoff to down-stream water bodies and meadows. |
| *Community values:* |  |
| Fire risk to property | Measured by the value of properties threatened by predicted wildfires. |
| Wildland-urban interface fire risk | The percent of forest in areas, near communities, at risk of burning at high severity. |
| Quality air | Represented using a proxy based upon the number of predicted high, very high, and extreme emission days. |
| Carbon sequestration | Represents emissions with global warming implications including carbon stored in the entire system (in-forest and harvested wood products). |
| Restoration byproduct | Indicates the predicted amount of biomass and wood product utilization resulting from a management scenario. |
| Cultural resource quality | Evaluated through a synthesis of indicators important to the Washoe Tribe, including predicted amounts of low-intensity fire, habitat for culturally important terrestrial species (e.g., deer, flicker, mountain quail, and aspen), and beneficial water flows to meadows and water bodies. |
| Recreation quality | Measured by summer-time smoke impact and winter-time snowpack duration. |
| *Management operations:* |  |
| Net treatment cost | Consists of cost of treatments (thinning and prescribed burning) less value of products removed. |
| Suppression cost | All cost related to suppressing wildland fires. |
| Staffing | Represents the number of agency personnel required to implement forest management projects. |
| Days of intentional burning | The number of days of prescribed fire or pile burning. |

### Decision support tool development

We focus here on how values derived for attributes were prioritized by stakeholders and how the interaction of priorities, current conditions, future climate, and management resulted in the overall performance of each of the five management scenarios (Table 1). Conditions across such a wide array of resources within a landscape could be considered equally important, but that is rarely the case. Most often, individuals and institutions have specific priorities at any specific point in time, motivated, for example, by perceived need, concern, risk, or institutional missions.

We used multi-criteria decision analysis (MCDA; Kamenetsky 1982, Saaty 1994, Mendoza et al. 2006, Murphy 2014) to compare the performance of the five scenarios in terms of the desired condition outcomes established for the 16 attributes associated with the three focal areas (Figure 1, Appendix 1). The performance scores of the multi-criteria decision model (MCDM) in EMDS are derived by the decision engine of Criterium DecisionPlus (Murphy 2014). Criterion weights in the decision hierarchy were derived using Saaty’s (1994) pairwise comparison methods in the Analytic Hierarchy Process (AHP). Using moderated discussion groups, we convened a group of 24 stakeholders (National Forest Foundation 2019) to derive the relative weighting of the 16 attributes in our DST (Abelson et al. *in review*). Stakeholders also rated the utility of each of the 16 attributes using the Simple Multi-Attribute Rating Technique (SMART; Kamenetsky 1982), where the utility of an observed value for each attribute indicates how well the observed value contributes to the model’s (Figure 1) overall performance score.

Weights are calculated on a scale of 0 to 1, and sum to 1 at each level of the hierarchy, starting with the 16 attributes. The weighting assigned to the attributes ranged from a low of 1% to a high of 15% (Figure 1), with six attributes carrying 78% of the weight: water quantity (15%), quality water (15%), wildlife conservation (13%), functional fire (13%), fire threat to property (11%), and wildland-urban interface fire (11%). The remaining 10 attributes carried a total of 22% of the weight and individually were assigned weights equivalent to ≤ 5%. Attribute weights propagate up through each level of the hierarchy (Figure 1). The primary objective of the LTW project was to improve environmental quality, and this is reflected in the environmental quality focal area being weighted most heavily at 59% of the evaluation. In contrast, community values carried 29% and operations carried 12% of the weight in the evaluation.

### Scenario performance scores

MCDM performance scores were derived by evaluating conditions for the 16 attributes for each the five alternative scenarios at each of 10 time steps, and these results were plotted and summarized. The overall performance score of each scenario was calculated in CDP as the sum of products of the attribute weights (range = 0 to 1) and their utility scores (range = 0 to 1), which then carried through each level of hierarchy such that performance can be evaluated at any level.

Performance scores range from 0 to 1; scores close to 1 indicate that conditions are optimal because they are approaching the target desired conditions, whereas scores close to 0 indicate suboptimal conditions because they are deviating from target desired conditions. We divided the range of values from 0 to 1 into five intervals of 0.2 to aid in the interpretation of differences in performance among scenarios (Table 3). We used both quantitative and qualitative approaches to compare the performance of scenarios with respect to maintaining or achieving desired conditions. Quantitative metrics included the mean, standard deviation, range (i.e., minimum and maximum values) in performance scores for any given level of the hierarchy, as well as the number of decades above a specified threshold (i.e., 0.8 or 0.6). Standard deviation and range can be informative to decision-makers because predictability may be more important in some situations (e.g., it may be preferable to have a scenario that slightly underperforms over the full temporal trajectory but does not have large year-to-year swings in forest condition). Qualitative metrics rank and compare scenarios regarding their relative performance compared to other management scenarios, both at any given time point and across temporal trajectories.

**Table 3.** Decision model outputs (range from 0 to 1) interpreted in terms of performance scores and associated condition classes in the evaluation of management scenarios in the Lake Tahoe West landscape restoration project.

| Performance score | Condition class |
| --- | --- |
| 0.8 – 1.0 | Optimal |
| 0.6 – 0.8 | Good |
| 0.4 – 0.6 | Marginal |
| 0.2 – 0.4 | Suboptimal |
| 0.0 – 0.2 | Poor |

### Sensitivity analysis

CDP models output sensitivity statistics (originally described for AHP models by Saaty 1994). In the case of our CDP modeling, the sensitivity analysis was repeated at each 10-year time step to assess model sensitivity over the 100-year period for each management scenario. The key metric produced by the sensitivity analysis is *criticality,* which is the absolute percent change in a criterion weight that would cause the top-rated scenario in our analysis results to be replaced by another scenario. CDP output, like most commercial MCDMs that implement the AHP, provides a *criticality score* for all attributes. The criticality score is a measure of model sensitivity in the sense that the identification of the top-rated scenario may be sensitive to how the attributes have been weighted. Low values of criticality indicate high model sensitivity to the associated attribute, and, conversely, high criticality values indicate relative model robustness (e.g., insensitivity). A simple way to assess model robustness is to examine the criticality value of the most sensitive attribute. The long-standing and well-accepted heuristic for judging robustness was proposed by Saaty (1994) as a criticality value of at least 10 percent for the most sensitive criterion. We follow this convention in assessing the robustness of our CDP models in the results.

## RESULTS

### Overall scenario performance

Relatively speaking, Scenario 5 (extensive fire-based approach, Table 1) outperformed all other scenarios, while Scenarios 1 and 2 consistently underperformed (Figure 2, Appendix 6). Overall, the five scenarios formed three clusters: Scenario 1 (fire suppression only) and Scenario 2 (wildland-urban interface focus) had similar and the lowest performance scores that trended together over time. Scenario 3 (thinning-based approach) and Scenario 4 (fire-based approach) trended together with moderate performance scores, while Scenario 5 consistently stood alone with the highest performance values. Specific aspects of overall scenario performance are explored below.

**Figure 2.**
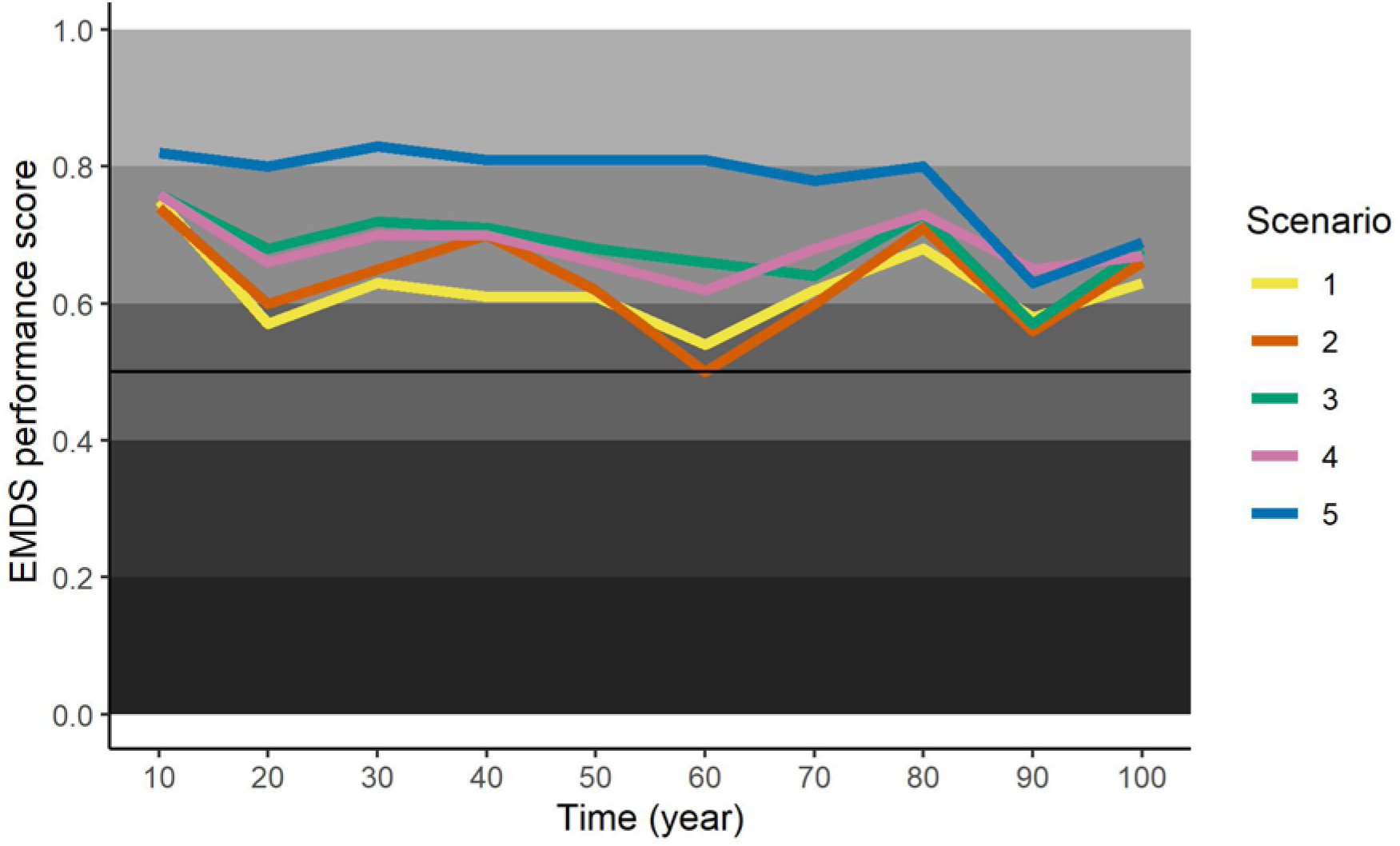
Performance of five management scenarios (S1-S5) in terms of meeting overall desired conditions over a 100-year time period (2010-2110) on the west side of the Lake Tahoe basin. Scenarios are arrayed from minimal management investment (S1) to landscape-wide management using thinning (S3) or fire (S4 and S5). Forest conditions resulting from each of management scenarios were modeled using LANDIS-II and evaluated each decade across an array of metrics representing environmental quality, community values, and management operations using the Ecosystem Management Decision Support tool (EMDS). Scenarios with performance scores closer to one indicate that optimal conditions resulted from management, while performance scores near zero that indicate poor conditions resulted.

#### Mean performance of scenarios

Scenario 5 was the best performing scenario based on the mean performance score (mean = 0.78, SD = 0.07), indicating nearly optimal performance across the 100-year period (Figure 2; Appendix 6). All other scenarios had good performance scores, but the lowest scoring scenario (Scenario 1) came close to marginal performance. The remaining scenarios, in descending order of performance, were Scenario 3 and Scenario 4, both with mean performance scores of 0.68 (SD = 0.05, SD = 0.04, respectively), followed by Scenario 2 with a mean value of 0.63 (SD = 0.07) and Scenario 1 with a mean value of 0.62 (SD = 0.06). The variance of these estimates indicates that Scenarios 1-4 were not well differentiated in their performance.

#### Year-to-year variation in scenario performance

Variation over time reflects uncertainty in management outcomes, so lower year-to-year variation is most often preferable. In general, year-to-year variation within scenarios was low, with standard deviations being ≤ 10% of mean values. In the first 80 years, Scenarios 3, 4, and 5 were relatively stable with regard to inter-decadal variability; in contrast, Scenarios 1 and 2 were considerably more variable (Figure 2).

Scenario 4 resulted in forest conditions that had the least year-to-year variation (SD = 0.039). Over the first 80 years, Scenario 5 had the lowest variability (SD = 0.016), however conditions deteriorated in year 90 with only a slight recovery in year 100. As a result, Scenario 5 had a nearly the highest standard deviation (SD = 0.065) across full the 100-year time frame. Scenarios 3 and 1 had intermediate standard deviation values of 0.053 and 0.059, respectively. Scenario 2 had the largest standard deviation value of 0.074.

#### Residency time in condition classes

Another facet of performance is the amount of time that conditions meet or approach desired target conditions, particularly when desired conditions are expected to be most resilient to disturbance. In general, residency times in good and optimal conditions were greater with increased management and increased use of fire (Figure 2). Scenario 5 (extensive use of fire) had the greatest number of decades in optimal (n = 7) and good (n = 3) conditions over the 100year period evaluated. Scenario 4 (use of fire) had the next highest residency times associated with desired conditions, with no decades in the optimal condition but all 10 decades in the good condition. Scenario 3 (primarily thinning) performed almost as well as Scenario 4, with all but one decade resulting in good conditions. Scenarios 1 and 2 performed the worst by this measure: Scenario 2 had eight decades in the good range and two in the marginal range, while Scenario 1 had seven decades in the good range and three in the marginal range.

### Focal area outcomes and contributions to scenario performance

Scenario performance was a function of the conditions of each of the three focal areas (environmental quality, community values, and management operations) and their relative weights. In an evaluation of management effectiveness, it is important to understand how the component parts of the system are faring, in addition to overall performance. For example, the same mean performance score of “good” could result from two very different landscape conditions: 1) all three focal areas are in good condition; or 2) one focal area is in poor condition and the other two are in optimal condition. Here we delve into how individual focal area conditions contributed to the observed overall performance of each scenario.

#### Environmental quality

Environmental quality was the primary objective of the Lake Tahoe West project, and consequently it was weighted (and contributed) the most (59%) to overall scenario performance (Figure 1). Scenario 5 (extensive fire) outperformed all other scenarios with little variation between decades (Figure 3, Appendix 6). Environmental quality remained within an optimal range under management Scenario 5 over the entire century with nominal variation (mean = 0.87, SD = 0.028). In contrast, the four remaining scenarios achieved nearly consistently good, but not optimal, environmental quality. Scenario 3 produced performance scores with a mean value of 0.72 (SD = 0.014), and Scenario 4 performed slightly worse with a mean value of 0.68 (SD = 0.030), reflecting their associated intermediate management inputs. Scenarios 1 and 2 (limited management) performed the worst, but were still in the good range (mean = 0.65, SD = 0.042 and mean = 0.62, SD = 0.044, respectively), with higher between-decade variability (periodic drops into the marginal condition range).

**Figure 3.**
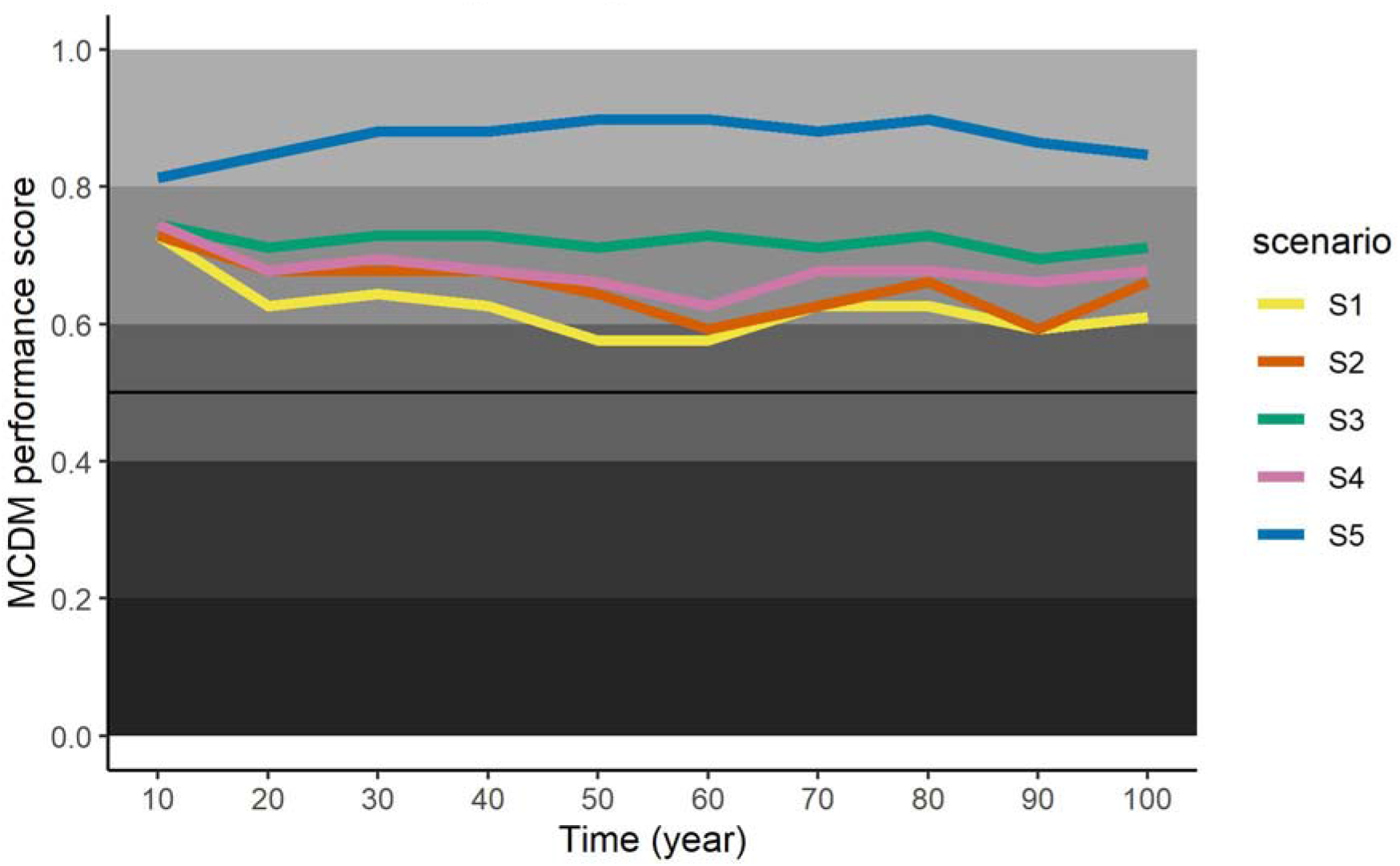
Performance of five management scenarios (S1-S5) in terms of meeting desired environmental quality outcomes over a 100-year time period (2010-2110) on the west side of the Lake Tahoe basin. Scenarios are arrayed from minimal management investment (S1) to landscape-wide management using thinning (S3) or fire (S4 and S5). Forest conditions resulting from each of management scenarios were modeled using LANDIS-II and evaluated each decade across an array of metrics representing environmental quality, community values, and management operations using the Ecosystem Management Decision Support tool (EMDS). Scenarios with performance scores closer to one indicate that optimal conditions resulted from management, while performance scores near zero that indicate poor conditions resulted.

In general, scenario performance for environmental quality had low variability over the 100-year period, regardless of the management scenario, as reflected in the small standard deviation values for mean performance scores. This indicates that management is, or at least can be, the primary driver of environmental quality, as opposed to stochastic or dynamic factors associated with forest ecosystems in the Lake Tahoe West landscape. Another salient aspect of performance was the consistent relative ranking of scenario performance in environmental quality over time. Specifically, for nearly every decade, Scenario 5 was the highest ranked scenario followed by Scenarios 3, 4, 2, and then 1.

Environmental quality was represented by two topics: terrestrial and aquatic environmental quality, both of which carried equal weight in representing environmental quality (Figure 1). For terrestrial environmental quality, two of the three attributes carried most (86%) of the weight – functional fire and wildlife conservation. For aquatic environmental quality, the two attributes (water quality and quantity) carried equal weight. Because of the relative equity of the topics and all but one of the attributes (44-50% of the weight other than upland vegetation health with just 14% of the weight), environmental quality condition scores truly reflect a composite of conditions, as opposed to some version of stakeholder priorities. In terms of scenario performance, terrestrial conditions started and remained good to optimal over the course of the century (min = 0.69, max = 0.93), regardless of management scenario (min mean = 0.74, max mean = 0.89). Differences in terrestrial conditions were largely driven by functional fire, which was the only one of the three attributes that varied in condition class over time (min = 0.52, max = 0.93), and varied by management scenario (min mean = 0.66 for Scenario1, max mean = 0.88 for Scenario 5). Aquatic conditions varied slightly more over the course of the century from mediocre to optimal (min = 0.45, max = 0.93), and similarly by management scenario (min mean = 0.53 for Scenario 1, max mean = 0.88 for Scenario 5). Within aquatic conditions, water quality was uniformly optimal, whereas water quantity was much more variable, ranging from poor to optimal (min = < 0.01, max = 1), and driven strongly by management scenario (mean = 0.13 for Scenario 1, max mean = 0.88 for Scenario 5).

#### Community values

The community values were an important but secondary focal area in the Lake Tahoe West landscape restoration project, and as such it carried 29% of the weight in overall scenario performance (Figure 1). Overall, the more management intensive scenarios (Scenarios 3, 4, and 5) performed well, resulting in generally good conditions over most of the entire century.

Scenario 3 performed the best (mean = 0.69, SD = 0.10), followed closely by Scenario 5 (mean = 0.68, SD = 0.18) and Scenario 4 (mean = 0.67, SD = 0.07). The variation within scenarios was high, making the performance of these top three scenarios indistinguishable. Scenarios 1 and 2 were more variable, which resulted in mediocre overall performance. (mean = 0.57, SD = 0.15, and mean = 0.53, SD = 0.11, respectively). Again, variation in performance within these two scenarios was high, obscuring differences in their performance.

In terms of residency time in desired conditions, Scenario 5 outperformed the other scenarios for community values over the first eight decades, although conditions steadily dropped from optimal in year 10 to good in year 80. Performance scores for all but Scenario 5 increased in the eighth decade. In year 90, while all scenarios have declining conditions, Scenario 5 dropped precipitously from good conditions to poor conditions.

Another interesting aspect of performance in community values was that, unlike environmental quality, the year-to-year variation in the community values focal area was not synchronous among scenarios (Figure 4). The best performing scenario in any given decade shifted frequently over time. For example, Scenarios 1 and 2 showed a large drop in performance in year 60, while Scenario 5 remained stable. In year 90, all scenarios experienced a drop in performance but the drop in Scenario 5 (0.41) was markedly more substantial than the other scenarios (ranging from 0.14 to 0.31; Appendix 6). This temporal variability in performance suggests that community values (as defined in this project) are more vulnerable to conditions created by dynamic processes and stochastic events as modeled in our landscape.

**Figure 4.**
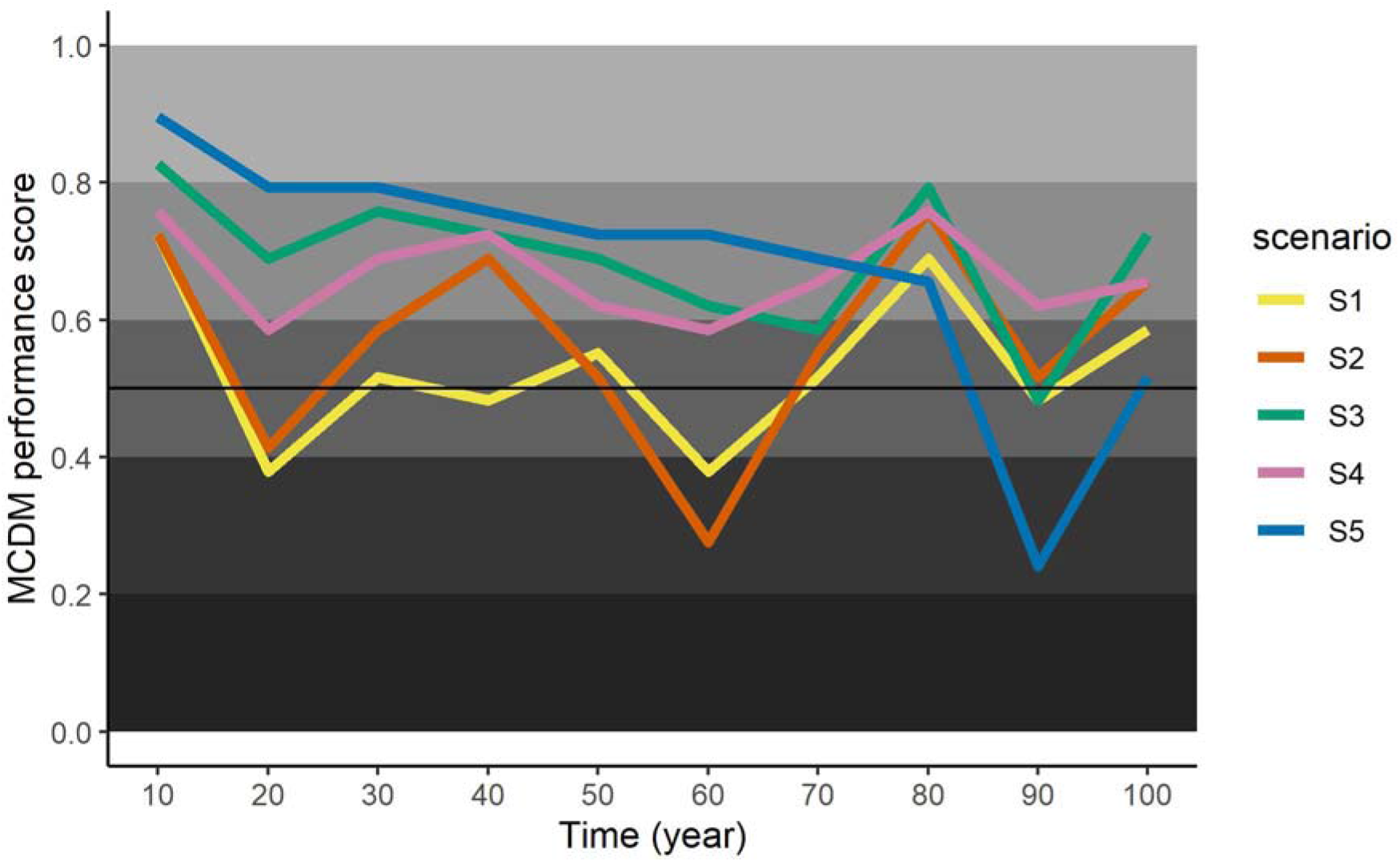
Performance of five management scenarios (S1-S5) in terms of meeting desired community values outcomes over a 100-year time period (2010-2110) on the west side of the Lake Tahoe basin. Scenarios are arrayed from minimal management investment (S1) to landscape-wide management using thinning (S3) or fire (S4 and S5). Forest conditions resulting from each of management scenarios were modeled using LANDIS-II and evaluated each decade across an array of metrics representing environmental quality, community values, and management operations using the Ecosystem Management Decision Support tool (EMDS). Scenarios with performance scores closer to one indicate that optimal conditions resulted from management, while performance scores near zero that indicate poor conditions resulted.

Community values were represented by two topic areas, with health and safety carrying 83% of the weight, and social values (carbon sequestration, recreation impacts from emissions, cultural resource quality, and wood byproducts from restoration) carrying only 17% of the weight (Figure 1). Thus, social values in this analysis, despite the widely held importance put on carbon sequestration (Forest Climate Action Team 2018) and cultural resources (Long 2019, Long et al. 2020) in particular, had limited influence on overall performance scores. For cultural resource quality, Scenario 5 was the only stand out, with good performance (mean = 0.64) compared to poor to suboptimal performance for all the other scenarios. None of the scenarios performed well for carbon sequestration, with Scenarios 3 and 5 having suboptimal performance (mean = 0.34 and 0.23, respectively), and the remaining scenarios having better but only marginal performance (range = 0.49 to 0.55).

Health and safety was represented by three attributes (Figure 1), two of which equally carried a total of 88% of the weight (wildland urban interface [WUI] fire risk and fire threats to property). WUI fire risk was the primary attribute driving observed performance scores for community values. It varied widely from poor (min = 0) to optimal (max = 0.97) over time, and varied among scenarios although generally declined over time across all scenarios (min mean = 0.26 for Scenario 1, max mean = 0.55 for Scenario 5) indicating a strong influence of climate on fire risk in the WUI. Risks to property varied over time from suboptimal to optimal (min = 0.33, max = 1.0), but retained mean performance values of optimal across all scenarios (min mean = 0.87, max mean = 0.97), indicating that realized risk to property is episodic in the form of infrequent but high impact events. Quality air, the third attribute, varied widely in conditions over time (poor to optimal) and among scenarios (marginal to optimal), it had little influence on performance.

#### Management operations

The management operations focal area was assigned the lowest importance, carrying just 12% of the total model weight (Figure 1). It exhibited unique and somewhat opposing scenario performance relative to the other two focal areas (Figure 5, Appendix 6). Generally, scenario performance declined with greater management input, primarily reflecting the cost of management. Mean scenario performance in the operations focal area was led by Scenario 1 with an optimal performance score of 0.83 (SD = 0.07), in contrast to community values and environmental quality focal areas, whereas Scenario 1 performed the worst or second worst of all the scenarios. The remaining scenarios generally followed degree of management input: Scenarios 4 and 2 with good performance (mean = 0.74, SD = 0.06, mean = 0.68, SD = 0.12), followed by Scenario 5 with marginal performance and high variability (and mean = 0.56, SD = 0.11, respectively), and finally Scenario 3 with suboptimal performance and high variability (mean = 0.48, SD = 0.14). Like the community values focal area, there was variability in the relative ranks of the scenario’s performance over time, with frequent shifts in relative performance from decade to decade. Notably, Scenario 5 was most often ranked nearly the lowest for operations, but was most often ranked the highest for the other two focal areas.

**Figure 5.**
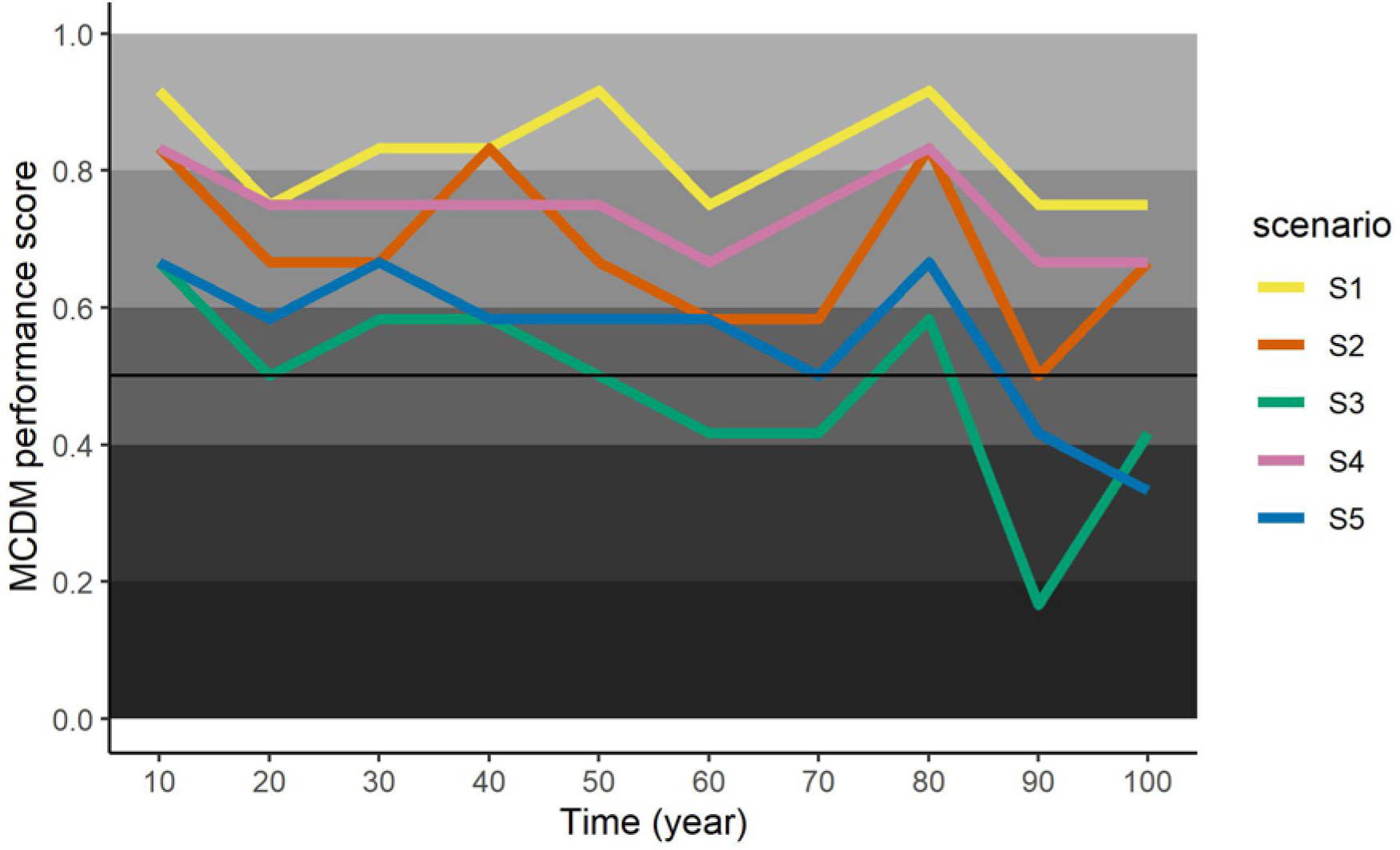
Performance of five management scenarios (S1-S5) in terms of meeting desired management operations outcomes over a 100-year time period (2010-2110) on the west side of the Lake Tahoe basin. Scenarios are arrayed from minimal management investment (S1) to landscape-wide management using thinning (S3) or fire (S4 and S5). Forest conditions resulting from each of management scenarios were modeled using LANDIS-II and evaluated each decade across an array of metrics representing environmental quality, community values, and management operations using the Ecosystem Management Decision Support tool (EMDS). Scenarios with performance scores closer to one indicate that optimal conditions resulted from management, while performance scores near zero that indicate poor conditions resulted.

Management operations were represented by two topic areas that carried equal weight: economics and feasibility (Figure 1), although recall that management operations itself only carried 12% of the weight, so the contribution of individual topic and attribute conditions was minimal. Economics varied widely from poor (min = 0) to optimal (max = 0.83), but varied less among scenarios from marginal to barely optimal (min mean = 0.53 for Scenario 3, max mean = 0.82 for Scenario 4). Within economics, the two attributes were treatment costs and suppression costs, with suppression costs carrying nearly all (83%) of the weight. However, suppression costs did vary widely over time (min = 0, max = 0.98) and by scenario from marginal to barely optimal (min mean = 0.62 for Scenario 2, max mean = 0.82 for Scenario 4). Interestingly, if net treatment costs (cost of removing material minus its market value) had been weighted more heavily, it would have been one of the few instances where Scenario 3 performed poorly, and the lowest ranked among all the scenarios (other than no management) with a mean score of 0.19. The remaining three scenarios were rated as good and were nearly identical in their mean performance scores (Scenario 2 = 0.71, Scenario 4 = 0.76, Scenario 5 = 0.60). Finally, feasibility varied widely from suboptimal (min = 0.33) to optimal (max = 1) and among scenarios (min mean = 0.35 for Scenario 5, max mean = 1.0 for Scenario 1). Within feasibility, the two attributes were staffing and days of intentional burning, with staffing carrying the majority (67%) of the weight. Staffing varied to a degree over time, ranging from suboptimal (min = 0.29) to marginal (max = 0.69), and it ranged more widely among scenarios from suboptimal (min mean = 0.36 for Scenario 3) to good (max mean = 0.72 for Scenario 4), which is not surprising given that management investments were held fairly constant over time within scenarios.

### Robustness of the decision model

We summarize the sensitivity analyses of the MCDM model results by the ten time steps (Table 4). Consistent with earlier results (Figure 2), Scenario 5 was the top-ranked scenario of the five management scenarios across all time steps except year 90, at which point Scenario 4 was the highest ranked scenario. Criticality values exceeded Saaty’s (1994) 10 % threshold heuristic for years 10 through 80, indicating a high degree of confidence in identifying Scenario 5 as the best performing scenario up through year 80. At years 90 and 100, the criticality statistic dropped to nearly 1%, indicating that a change in weight of the most sensitive focal area could readily shift away from Scenario 5 being the best performing scenario later in the century, when conditions become more variable. Management operations was the most sensitive focal area for most of the century (Table 4), meaning a sufficiently large change in its weight would cause Scenario 5 to be replaced as the top ranked alternative in years 10 through 80 by another scenario for a given decade (Table 4); however, as already noted, the criticality results for years 10 through 80 indicate a robust model with Scenario 5 ranked highest in performance in this period. In contrast, in years 90 and 100, the criticality scores indicate that we could not practically distinguish between Scenarios 4 and 5 as the top ranked alternative.

**Table 4.**
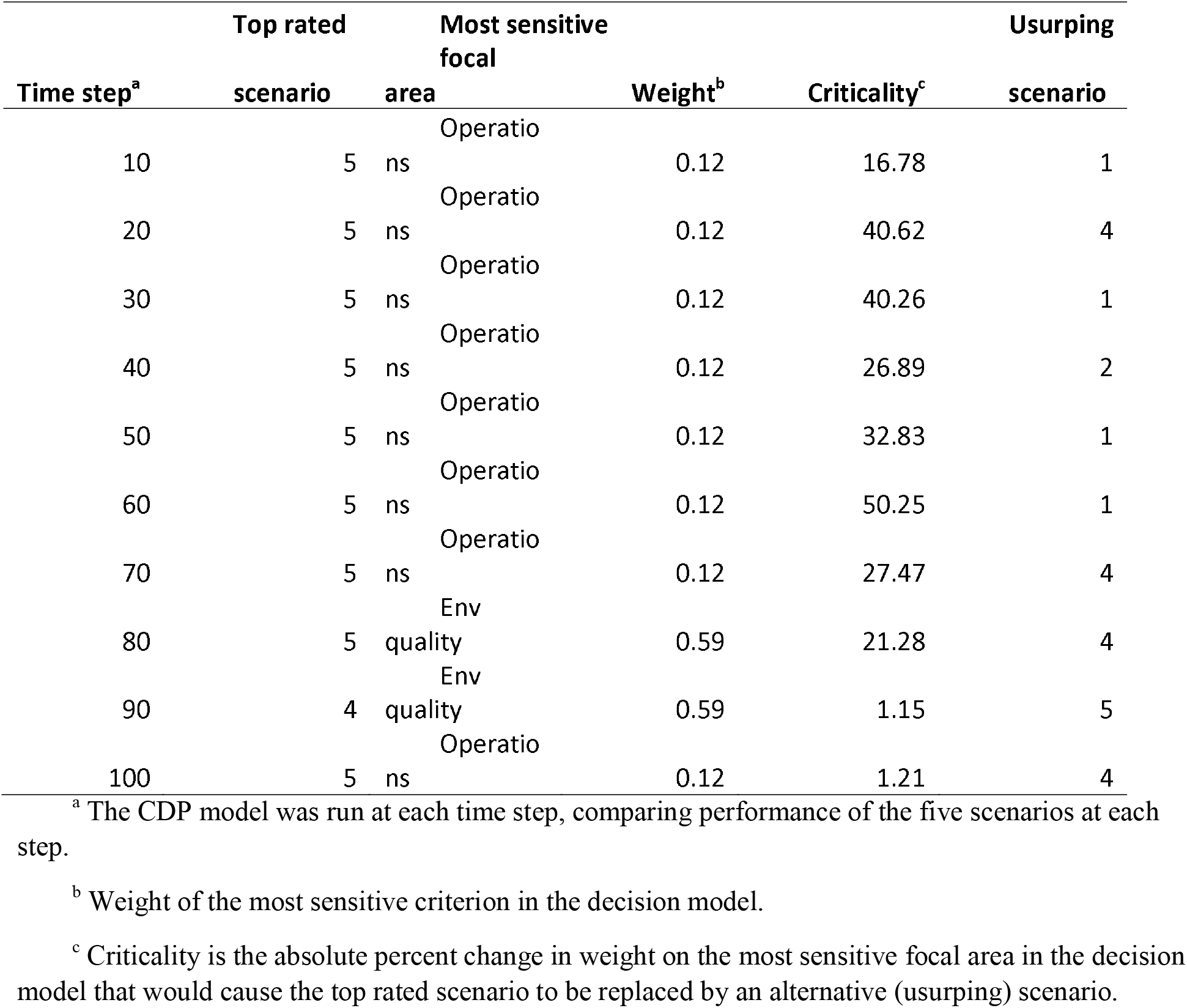
Summary of sensitivity analyses calculated by Criterium DecisionPlus (CDP) for each of the 10 decadal time steps of the 100-year period modeled.

## DISCUSSION

Management activities influence forest dynamics over long time-horizons and are increasingly essential for cultivating resilient forest landscape conditions. The decision support tool (DST) that we developed using EMDS demonstrated that management investments can make a substantial difference in creating and maintaining desired forest landscape conditions over time across the westside of the Lake Tahoe basin, despite a changing climate. Specifically, forest landscape conditions benefitted substantially by management activities that reduced woody biomass through thinning and intentional fire, resulting in a more productive role of fire as a natural process (i.e., functional fire), wildlife habitat conservation, and improved water quantity and availability (Figure 6). The DST enabled the transparent assessment of many complex, and often compensatory, attributes that were informed by considering forest and climate dynamics 100 years into the future. Model transparency also provided many insights into how stakeholder priorities can greatly affect the outcomes and perceived effectiveness of various management approaches.

**Figure 6.**
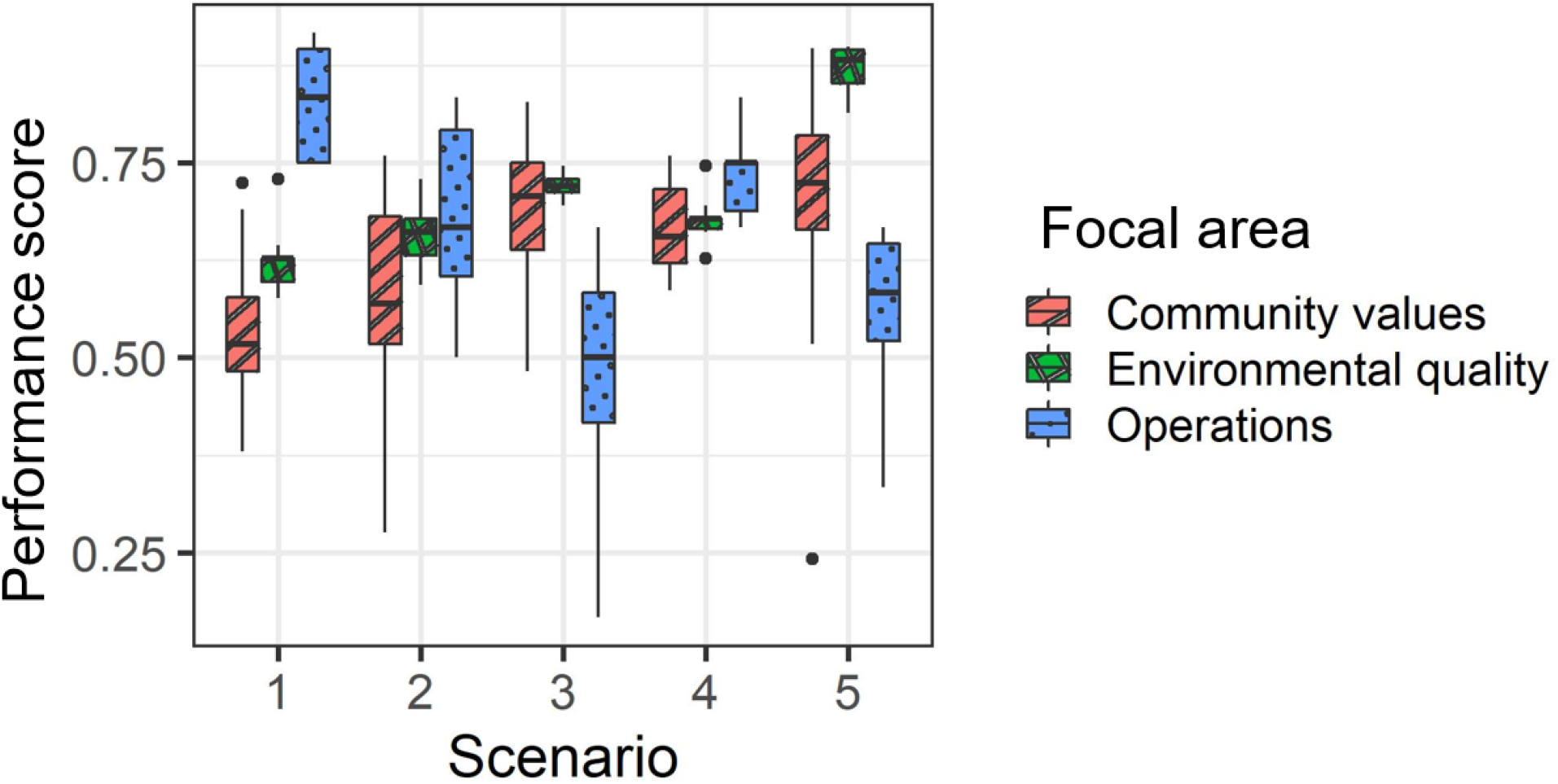
Comparison of the performance of five management scenarios in meeting objectives for three focal areas (environmental quality, community values, and management operations) based on a multi-criterion decision model with 16 attributes for conditions across the west side of the Lake Tahoe basin.

### Landscape-wide management is important

Our analysis results resoundingly showed that increasing the extent of management across the landscape (as represented by Scenarios 1 to 5) improved the achievement of desired outcomes (Figure 6). We found substantial differences among the management strategies in terms of their influence on landscape conditions, and environmental quality, in particular, over the course of the 21^st^ century in markedly different ways. Performance closely followed the extent of management across the landscape – the greater the number of acres treated, the more positive the outcome for environmental quality based on the performance score. This is consistent with other studies that found that treating large landscape areas would mitigate impacts of severe wildfires (Moghaddas et al. 2010). Stevens et al. (2016) conducted a similar study in the northern portion of the LTW landscape, and although they used a different fire and vegetation modeling framework and did not incorporate climate change, they also found that more management treatment had beneficial effects by reducing the prevalence of high severity wildfire.

In Lake Tahoe, as throughout California and the western U.S., the threat of wildfire to forests, property, and people has become a primary driver in forest management, with the primary objective being to reduce the probability that large, high severity fires will occur (Hessburg et al. 2016). Our results showed that forest management treatments consisting of thinning and/or prescribed fire were effective at reducing the risk and probability of significant and destructive wildfires. Scenarios 1 and 2 were more variable; this in part reflects that extreme wildfires in one period may inhibit the incidence of fire in the same area for many years, while more dispersed treatments and fires tend to moderate fire effects over time (e.g., Scholl and Taylor 2010, Perry et al. 2011). This finding is consistent with previous synthesis work (e.g., Long et al. 2014) and landscape modeling in the region (e.g., Krofcheck et al. 2017). Previous research (Ager et al. 2010, 2013) had found that concentrating treatments in smaller areas, such as the WUI, could reduce the risk of damage in those areas (including buildings), but at the expense of greater loss of old trees in the untreated areas.

The two most aggressive treatment scenarios (Scenarios 3 and 5) treated approximately 7% of the landscape each year, or roughly 50-70% of the forested landscape per decade (locations can be treated more than once per decade), which equates to a 20-year disturbance frequency. This is consistent with historic fire return intervals. Previous research has suggested that gains in performance in some metrics, such as fire and carbon outcomes, can be achieved by targeting treatments to the areas at greatest risk of high-severity fire (Stevens et al. 2017, Krofcheck et al. 2017).

### Superior performance of landscape fire as a management tool

One of the most important results of our analysis was that the expanded use of fire as a management tool to reduce forest biomass, as represented by Scenario 5, was consistently superior to other management approaches in achieving and maintaining optimal desired conditions, particularly regarding environmental quality and overall condition (Figure 6). In addition to reflecting the ecological benefits of fire, as reported by others (e.g., Agee and Skinner), the strong ecological performance of Scenario 5 was bolstered by the emphasis (i.e., weights) that stakeholders put on the importance of increasing functional fire and reducing the risk of large, high severity fires (particularly in the WUI) while placing relatively low emphasis on the cost and logistics of treatments (e.g., restrictions on when burning is permitted) (e.g., Kolden 2019). The results from these analyses provided a scientific foundation for managers to propose a gradual increase in the use of fire as part of their restoration strategy for Lake Tahoe West (Lake Tahoe West Restoration Partnership 2019).

Stability did not explicitly alter performance scores of management strategies directly, but some ecological processes (e.g., persistence of species at risk of local extirpation), and many aspects social well-being (e.g., consistent conditions to which people can adapted versus periodic significant impacts) favor stability (Angeler and Allen 2016, *sensu* Johnstone et al. 2016). We found that scenarios that treated smaller proportions of the landscape per year and treatments that were more concentrated around the WUI showed greater variability in landscape conditions over time. Consistent with other studies (e.g., Maxwell et al. 2020), this outcome appeared to be a function of the greater probability of large and high severity wildfires when smaller proportions of the landscape were receiving management inputs in the form of thinning or fire. Frequent low to moderate severity fire is particularly effective in reducing the risk of future high severity fires in any given area (Omi and Martinson 2004, Arkle et al. 2012). Scenarios that extensively used prescribed burning (Scenario 5 in this study) performed well in terms of achieving desired conditions, as well as producing stable and reliable condition outcomes (i.e., low variability over time and low sensitivity to weighting) over the course of at least the first 80 years.

Fire as a management tool has many advantages, which were reflected in the performance of Scenario 5, but also has some disadvantages, which although they were given lower weight by stakeholders, are still recognized as important. Prescribed fire as a tool results in smoke emissions, which are less impactful than wildfire, so shifting toward intentional fire can result in a net benefit in terms of reduced public health impacts (Long et al. 2018, Schweizer and Cisneros 2016). Our modeling also indicated that use of fire was less expensive than thinning, although this carried minimal weight in the evaluation. Challenges with using prescribed fire include having a sufficient number of allowable burn days (based on suitable weather conditions and air quality) and adequate staffing to accomplish the work – both of which are commonly limiting factors in implementation (Ryan et al. 2013). In our model, stakeholders considered feasibility a low priority so, although feasibility is the primary reason more burning is not accomplished (Kolden 2019), it did not substantially downgrade the performance of the extensive burning scenario. The decision model helped highlight two important outcomes: 1) that given sufficient social license and commitment of resources, optimal environmental outcomes can be achieved but only through the use of extensive fire; and 2) knowing that desired conditions can be achieved may, in turn, help agencies and communities rally to overcome the social barriers to its implementation (McCaffrey et al. 2013).

### End of century variability and uncertainty

Dynamic modeling requires parameterizing many processes and interactions, based primarily on estimates derived from empirical studies (Seidl 2017). Future climate conditions are also estimates based on modeling and this introduces even greater uncertainty as it is not certain which of the multiple plausible future conditions are most likely to occur (Scholes et al. 2014). The value in longer-term modeling is to look for trends, thresholds, tradeoffs (Kim et al. 2017). We modeled a 100-year period to explore trends and variability within and among management scenarios, but the uncertainty in how climates will change over this time frame makes observed dynamics in later decades less reliable. In our study, we found instabilities in later decades for the community values and management operations focal areas. Specifically, in year 80 and 90 we observed declines in the more aggressive management scenarios. Scenario 5’s decline in year 90 was due to the interaction of the prescribed fire parameterization and the repetition of 2100 climate conditions out to 2110 (due to lack of climate modeling data after 2100), and, as such, may reflect model anomalies.

### Influence of stakeholder priorities on perceived management performance

In our DST, community values and operations (together comprising 41% of the total model) did not carry as much weight as ecological considerations (which alone comprised 59% of the model) though they still played an important role (Figure 1). While environmental conditions over the course of the century showed a clear and consistent positive response to expanded treatment and expanded use of fire, variable (and sometimes compensatory) responses over time were observed in community values and operations (Figures 4 and 5). A main finding was that management scenarios that did not reduce densities of small trees and associated fuels resulted in landscapes that were prone to more severe fire events, which then set in motion larger oscillations between poor and good community values and operational conditions. While wildfires can be an effective fuel reduction tool, managers are increasingly concerned that uncharacteristically severe wildfires will result in long-lasting losses of social and ecological values (e.g., Spies et al. 2019).

The aquatic topic area was weighted heavily by stakeholders, accounting for 50% of environmental quality. This is likely to reflect the exceptional importance of restoring and maintaining the clarity of Lake Tahoe, although most efforts to date have focused on urban sources of impact rather than forest lands (Riverson et al. 2008). Our DST used LANDIS-II projections (Scheller et al. 2018) for leaf-area-index (LAI) as the proxy for water quantity, and scenarios with increased thinning and burning treatments (both of which reduce LAI through removal of live biomass) performed well. Although increased management activity modestly increased loading of fine sediment particles and phosphorus, those increases generally offset loads from wildfires; as a result the differences across scenarios were not substantial. For example, previous studies have found that increased treatments can realize benefits to aquatic systems by mitigating wildfire impacts to water quality (Buckley et al. 2014) and increasing water yield (Roche et al. 2020). Further, results of hydrologic modeling conducted in the study area indicated that thinning small trees (up to 20 m tall) would reduce LAI and also increase snow accumulation and melt volume (Krogh et al. 2020), further enhancing water quantity. While our results suggest that managers in the basin ought not to expect substantial improvements in water quality from forest management, it importantly indicates that water quality concerns do not limit larger restoration efforts. Our results indicated that both water quality and quantity were likely to degrade in the future without broad-scale use of fire (Appendix 6), which provides compelling evidence to overcome near-term concerns about the impacts of forest treatments, fire in particular, on Lake Tahoe’s water clarity. Our findings add to the growing body of literature (e.g., Nair and Howlett 2016) that indicate that effective approaches to promoting long-term ecosystem resilience focus on longer time-frames, adapting to change, and consider a wide range of interacting values, as opposed to a focus on near-term risk reduction and maintaining the status quo.

Both the attributes of community values and operations had high year-to-year variability, and to some extent gains in community values are compensated by losses in operations. That is, while Scenario 1 generally performed poorly (and Scenario 5 generally performed very well) regarding the community values because of the high risk of property damage, the opposite was true for the performance of management operations because the suppression-only scenario was less expensive in terms of implementation costs. Various studies have suggested that suppression-only strategies fail to account for the full social costs of future disturbances (Buckley et al. 2014, Bagdon and Huang 2016), and that restoration-based strategies can achieve social and ecological win-win outcomes (Bagdon et al. 2016, Spies et al. 2019).

In the DST, environmental outcomes were weighted nearly 2:1 over social and economic outcomes. As a result, management activity was favored over management costs, and environmental outcomes over feasibility. For example, increasing treatments were needed to mitigate impacts from future wildfires, but the type and extent of management were not substantially tempered by factors other than environmental outcomes. These choices were made intentionally to help ensure that strategies would promote ecological resilience without being unduly constrained by policies, regulations, and other institutional limitations (e.g., Stephens et al. 2013, Schoennagel et al. 2017) However, if community values and management operations had been given more weight, it is likely that Scenario 5 may not have been as much of a standout in terms of performance, perhaps Scenario 4 would have fared as well or better, as reflected in the sensitivity analysis (Table 4), and Scenario 3 probably would perform more poorly, given the high cost of thinning. In short, there is no gain (progress) without pain (cost by some measure). Another study of forest landscape collaboratives found that social values often received less attention than ecological ones, and that participants were sometimes discouraged when projects were planned that were proved economically infeasible (e.g., Urgenson et al. 2017); they suggested that clear articulation of tradeoffs is important.

### Decision support for landscape resilience

EMDS has been used since the late 1990s to provide decision support for numerous management issues related to environmental analysis and planning (Reynolds et al. 2014 and work cited therein), but its application to the LTW resilience project is novel. Whereas previous applications have been point-in-time analyses based on the current observed state of ecosystems (e.g., Cleland et al. 2017, Hessburg et al. 2013), the LTW DST considers the evolution of ecosystem states over a 100-yr timeframe based on the LANDIS-II process model. Examining a long time-horizon provided a framework for an analysis of ecosystem resilience (Holling 1973, Walker and Salt 2010) in the sense discussed in the Methods section. The LTW resilience analysis was facilitated by a new EMDS workflow (Chappell 2009) component that automates model processing by invoking sequences of models as activities and by using iterative loops. To avoid belaboring the details, we simply note that workflow automation reduced the task of performing many model runs to pressing a button to execute all the required models. For an in-depth discussion of the workflow as implemented in the LTW DST, see Abelson et al. (in review).

In other respects, however, the LTW DST is similar to other EMDS applications in its use of knowledge-based reasoning (Walters and Neilson 1988) to assess ecosystem states and evaluate alternative management scenarios. Especially in the past 20+ years, analyses and assessments in decision support have increasingly turned to knowledge-based methods to address contemporary management issues related to the sustainability, integrity, and resilience of ecosystems because these are large, complex, and abstract topics that are not easily modeled using traditional scientific analytical tools (Swanson and Greene 1999, Gunderson 1999, Reynolds 2001). Both of the analytical tools applied to the LTW resilience analysis (NetWeaver and CDP) are knowledge-based systems, which are useful in complex analytical contexts because, essentially, if one can reason about the problem at hand, then it can be modeled.

Here, we focused on CDP in particular because our primary objective here was to evaluate the performance of the five management scenarios (while NetWeaver was principally used to preprocess complex data before inclusion in the CDP model). A virtue of MCDMs in general, and CDP in particular, is that they are rationale, transparent and repeatable (Murphy 2014). That said, MCDM models typically operate at the interface between science and policy and have an inherent, unavoidable element of subjectivity because they require judgments from decision makers as to the relative importance of the criteria (i.e., the focal areas and topics in our analysis). Indeed, there is no such thing as a value-free decision in environmental management. Deciding what topics matter, and how much they matter, are critical in the design of an MCDM for decision making in general, and decision making in environmental management in particular. The sensitivity analysis for the LTW model of scenario performance (Table 4) is helpful in balancing subjectivity on the one hand and a rational, transparent, and repeatable model on the other, because it highlights those model weights that are most sensitive to determining the relative ordering of performance scores, and this is useful to focus discussion about choices that have been made in MCDM weights.

## CONCLUSION

Interdisciplinary modeling is becoming an essential feature of bringing the best available information to bear on landscape restoration planning and implementation. This is largely driven by observed and projected future climate change, and the threat it poses to long term sustainability and resilience of ecological and social systems. Future dynamics and possible characteristics and impacts of extreme events, as well as chronic stress from directional change in temperature or precipitation, are essential to attempt to understand (at least in terms of bounding what is probable). However, the ways in which climate change will alter landscapes and ecological perturbations are not fully understood, and projections of future conditions and dynamics become less reliable the further out in time they go. Decision makers rarely make project-level decisions that commit to implementation spanning more than 5 to 10 years, and they are likely to change their management approaches in response to environmental outcomes over time. Consequently, decision-makers may be not be overly concerned with projections of future declines, so long as the strategy improves conditions in the near-term and maintain options for avoiding damaging losses in the long-term. Such an approach can be successful, provided that the management system is also designed to monitor and adapt to changing conditions in a timely manner.

Our modeling effort helped managers to consider a wide range of social and ecological values over multiple decades. Ultimately, the Lake Tahoe West restoration strategy proposed by LTWRP (Lake Tahoe West Restoration Partnership 2019) focused on increasing thinning treatments across the landscape while gradually ramping up the use of prescribed fire. This choice represented a mix of Scenarios 3, 4, and 5, which our model all suggested would promote greater ecological and social values over the status quo management approach (Scenario 2). Our decision framework identified the effects of climate change and different management strategies on resilience over multiple decades in addition to the consideration of tradeoffs among stakeholder values; taken together this helped decision makers avoid the trap of management informed only by short-term gains. This work also provided scientific understanding of the potential benefits and downsides of increasing the use of intentional fire. The use of such an integrated modeling system helps managers and stakeholders to envision the challenges they will face in addressing dramatically changing future conditions and garnering support for implementing new management approaches. It also provides a ready-made system for evaluating management effectiveness, evaluating system conditions, and monitoring progress toward greater resilience.

## ACKNOWLEDGEMENTS

Amelia Wolf, Sarah Di Vittorio, Dorian Fougeres, Keith M. Slauson, Stacy A. Drury, Brandon M. Collins, William Elliot, Rob Scheller, Alec Kretchun, Mariana Dobre, Erin Brooks, Sam Evans, Tim Holland, Matthew Potts, Adrian Harpold, Sebastian Krogh Navarro. We also thank all the members of the LTW Science Team, LTW Interagency Design Team, LTW Stakeholder Science Committee, and the LTW Stakeholder Committee.

## Supplemental information

Appendix 1: CriterionDecision Plus (CDP) model in CDPX format.

Appendix 2: NetWeaver (NW) model in NW2 format.

Appendix 3: NetWeaver (NW) model in user readable format (HTML).

Appendix 4: NetWeaver (NW) model input specifying values used in NW fuzzy logic.

Appendix 5: Raw data inputs for the NetWeaver model (i.e. appendix 2) and CriterionDecison Plus (i.e. appendix 1)

Appendix 6: Multi-criteria decision model values and summary statistics for focal areas, topic areas, and attributes.

