## Appendix 3 for "Promoting long-term forest landscape resilience in the Lake Tahoe basin": _index.html

untitled0


### untitled0

- dependency outline
- groups
- dependency networks
- data

#### dependency outline

- LTW resilience final
  - Wildlife conservation
    - Species richness
      - SpecRich1
      - SpecRichI1
      - SpecRich2
      - SpecRichI2
      - SpecRich
    - Ecological function
      - Insectivore funct redund
        - Insectivore1
        - InsectivoreI1
        - Insectivore2
        - InsectivoreI2
        - Insectivore
      - Herbivore funct redund
        - Herbivore1
        - HerbivoreI1
        - Herbivore2
        - HerbivoreI2
        - Herbivore
      - Seed disp funct redund
        - Disperser1
        - DisperserI1
        - Disperser2
        - DisperserI2
        - Disperser
      - Scavenger funct redund
        - Predator1
        - PredatorI1
        - Predator2
        - PredatorI2
        - Predator
      - Decomp funct redund
        - Decomposer1
        - DecomposerI1
        - Decomposer2
        - DecomposerI2
        - Decomposer
      - Soil aerator funct redund
        - SoilAerator1
        - SoilAeratorI1
        - SoilAerator2
        - SoilAeratorI2
        - SoilAerator
    - Species diversity
      - Early seral beta div
        - EarlyBeta1
        - EarlyBetaI1
        - EarlyBeta2
        - EarlyBetaI2
        - EarlyBeta
      - Mid seral beta div
        - MidBeta1
        - MidBetaI1
        - MidBeta2
        - MidBetaI2
        - MidBeta
      - Late seral beta div
        - LateBeta1
        - LateBetaI1
        - LateBeta2
        - LateBetaI2
        - LateBeta
    - Apex predators
      - CASPO territories
        - CASPO1
        - CASPOI1
        - CASPO2
        - CASPOI2
        - CASPO
      - NOGO territories
        - NOGO1
        - NOGOI1
        - NOGO2
        - NOGOI2
        - NOGO
      - Marten territories
        - Marten1
        - MartenI1
        - Marten2
        - MartenI2
        - Marten
  - Quality water
    - Phosphorus load
      - Phosphorus1
      - PhosphorusI1
      - Phosphorus2
      - PhosphorusI2
      - Phosphorus
    - Fine sediment load
      - Sediment1
      - SedimentI1
      - Sediment2
      - SedimentI2
      - Sediment
  - Upland vegetation health
    - Big trees
      - BigTrees1
      - BigTreesI1
      - BigTrees2
      - BigTreesI2
      - BigTrees3
      - BigTreesI3
      - BigTrees
    - Forest cover
      - Conifer cover
        - Forest1
        - ForestI1
        - Forest2
        - ForestI2
        - Forest3
        - ForestI3
        - Forest4
        - ForestI4
        - Forest5
        - ForestI5
        - Forest
      - Hardwood cover
        - Hardwood1
        - HardwoodI1
        - Hardwood2
        - HardwoodI2
        - Hardwood3
        - HardwoodI3
        - Hardwood4
        - HardwoodI4
        - Hardwood
      - Shrub cover
        - Shrub1
        - ShrubI1
        - Shrub2
        - ShrubI2
        - Shrub3
        - ShrubI3
        - Shrub4
        - ShrubI4
        - Shrub5
        - ShrubI5
        - Shrub
    - Seral stage
      - High elevation
        - High elevation early
          - HighEarly1
          - HighEarlyI1
          - HighEarly2
          - HighEarlyI2
          - HighEarly3
          - HighEarlyI3
          - HighEarly4
          - HighEarlyI4
          - HighEarly5
          - HighEarlyI5
          - HighEarly6
          - HighEarlyI6
          - HighEarly
        - High elevation mid
          - HighMid1
          - HighMidI1
          - HighMid2
          - HighMidI2
          - HighMid3
          - HighMidI3
          - HighMid4
          - HighMidI4
          - HighMid5
          - HighMidI5
          - HighMid6
          - HighMidI6
          - HighMid
        - High elevation late
          - HighLate1
          - HighLateI1
          - HighLate2
          - HighLateI2
          - HighLate3
          - HighLateI3
          - HighLate4
          - HighLateI4
          - HighLate5
          - HighLateI5
          - HighLate6
          - HighLateI6
          - HighLate
      - Mid elevation
        - Mid elevation early
          - MidEarly1
          - MidEarlyI1
          - MidEarly2
          - MidEarlyI2
          - MidEarly3
          - MidEarlyI3
          - MidEarly4
          - MidEarlyI4
          - MidEarly5
          - MidEarlyI5
          - MidEarly6
          - MidEarlyI6
          - MidEarly
        - Mid elevation mid
          - MidMid1
          - MidMidI1
          - MidMid2
          - MidMidI2
          - MidMid3
          - MidMidI3
          - MidMid4
          - MidMidI4
          - MidMid5
          - MidMidI5
          - MidMid6
          - MidMidI6
          - MidMid
        - Mid elevation late
          - MidLate1
          - MidLateI1
          - MidLate2
          - MidLateI2
          - MidLate3
          - MidLateI3
          - MidLate4
          - MidLateI4
          - MidLate5
          - MidLateI5
          - MidLate6
          - MidLateI6
          - MidLate
    - Composition
      - Yellow pine forest
        - YellowPine1
        - YellowPineI1
        - YellowPine2
        - YellowPineI2
        - YellowPine3
        - YellowPineI3
        - YellowPine4
        - YellowPineI4
        - YellowPine
      - White pine forest
        - WhitePine1
        - WhitePineI1
        - WhitePine2
        - WhitePineI2
        - WhitePine3
        - WhitePineI3
        - WhitePine4
        - WhitePineI4
        - WhitePine
      - Aspen forest
        - Aspen1
        - AspenI1
        - Aspen2
        - AspenI2
        - Aspen3
        - AspenI3
        - Aspen4
        - AspenI4
        - Aspen
  - Functional fire
    - High sev patch
      - PerHSpatch1
      - PerHSpatchI1
      - PerHSpatch2
      - PerHSpatchI2
      - PerHSpatch
    - Percent land burned
      - Burned low sev
        - PercentLandscapeLow1
        - PercentLandscapeLowI1
        - PercentLandscapeLow2
        - PercentLandscapeLowI2
        - PercentLandscapeLow3
        - PercentLandscapeLowI3
        - PercentLandscapeLow4
        - PercentLandscapeLowI4
        - PercentLandscapeLow5
        - PercentLandscapeLowI5
        - PercentLandscapeLow6
        - PercentLandscapeLowI6
        - PercentLandscapeLow
      - Burned mod sev
        - PercentLandscapeMod1
        - PercentLandscapeModI1
        - PercentLandscapeMod2
        - PercentLandscapeModI2
        - PercentLandscapeMod3
        - PercentLandscapeModI3
        - PercentLandscapeMod4
        - PercentLandscapeModI4
        - PercentLandscapeMod5
        - PercentLandscapeModI5
        - PercentLandscapeMod
      - Burned high sev
        - PercentLandscapeHigh1
        - PercentLandscapeHighI1
        - PercentLandscapeHigh2
        - PercentLandscapeHighI2
        - PercentLandscapeHigh3
        - PercentLandscapeHighI3
        - PercentLandscapeHigh4
        - PercentLandscapeHighI4
        - PercentLandscapeHigh
  - WUI fire
    - Threat zone fire severity
      - WUI threat zone mod severity
        - WUITmod1
        - WUITmodI1
        - WUITmod2
        - WUITmodI2
        - WUITmod3
        - WUITmodI3
        - WUITmod4
        - WUITmodI4
        - WUITmod
      - WUI threat zone high severity
        - WUIThigh1
        - WUIThighI1
        - WUIThigh2
        - WUIThighI2
        - WUIThigh3
        - WUIThighI3
        - WUIThigh4
        - WUIThighI4
        - WUIThigh
    - Defense zone fire severity
      - WUI defense zone mod severity
        - WUIDmod1
        - WUIDmodI1
        - WUIDmod2
        - WUIDmodI2
        - WUIDmod3
        - WUIDmodI3
        - WUIDmod
      - WUI defense zone high severity
        - WUIDhigh1
        - WUIDhighI1
        - WUIDhigh2
        - WUIDhighI2
        - WUIDhigh3
        - WUIDhighI3
        - WUIDhigh
  - Quality air
    - Qual air extreme emission days
      - ExtremeEmissionDays1
      - ExtremeEmissionDaysI1
      - ExtremeEmissionDays2
      - ExtremeEmissionDaysI2
      - ExtremeEmissionDays3
      - ExtremeEmissionDaysI3
      - ExtremeEmissionDays
    - Qual air high emission days
      - HighEmissionDays1
      - HighEmissionDaysI1
      - HighEmissionDays2
      - HighEmissionDaysI2
      - HighEmissionDays3
      - HighEmissionDaysI3
      - HighEmissionDays
    - Qual air very high emission days
      - VHighEmissionDays1
      - VHighEmissionDaysI1
      - VHighEmissionDays2
      - VHighEmissionDaysI2
      - VHighEmissionDays3
      - VHighEmissionDaysI3
      - VHighEmissionDays
    - Qual air mod emission days
      - ModEmissionDays1
      - ModEmissionDaysI1
      - ModEmissionDays2
      - ModEmissionDaysI2
      - ModEmissionDays3
      - ModEmissionDaysI3
      - ModEmissionDays
  - Recreation
    - Sum extm emssn days
      - SumExtrmEmssnDays1
      - SumExtrmEmssnDaysI1
      - SumExtrmEmssnDays2
      - SumExtrmEmssnDaysI2
      - SumExtrmEmssnDays3
      - SumExtrmEmssnDaysI3
      - SumExtrmEmssnDays
    - Sum v high emssn day
      - SumVHighEmssnDays1
      - SumVHighEmssnDaysI1
      - SumVHighEmssnDays2
      - SumVHighEmssnDaysI2
      - SumVHighEmssnDays3
      - SumVHighEmssnDaysI3
      - SumVHighEmssnDays
    - Sum high emssn day
      - SumHighEmssnDays1
      - SumHighEmssnDaysI1
      - SumHighEmssnDays2
      - SumHighEmssnDaysI2
      - SumHighEmssnDays3
      - SumHighEmssnDaysI3
      - SumHighEmssnDays
    - Sum mod emmsn day
      - SumModEmssnDays1
      - SumModEmssnDaysI1
      - SumModEmssnDays2
      - SumModEmssnDaysI2
      - SumModEmssnDays3
      - SumModEmssnDaysI3
      - SumModEmssnDays
  - Cultural resource quality
    - Restoration of low intensity fire
      - CultFire1
      - CultFireI1
      - CultFire2
      - CultFireI2
      - CultFire3
      - CultFireI3
      - CultFire
    - Habitat for cultural keystone species
      - HCKS mule deer habitat
        - MuleDeer1
        - MuleDeerI1
        - MuleDeer2
        - MuleDeerI2
        - MuleDeer3
        - MuleDeerI3
        - MuleDeer
      - HCKS flicker habitat
        - Flicker1
        - FlickerI1
        - Flicker2
        - FlickerI2
        - Flicker3
        - FlickerI3
        - Flicker
      - HCKS mountain quail habitat
        - MtnQuail1
        - MtnQuailI1
        - MtnQuail2
        - MtnQuailI2
        - MtnQuail3
        - MtnQuailI3
        - MtnQuail
      - HCKS aspen habitat
        - CultASP1
        - CultASPI1
        - CultASP2
        - CultASPI2
        - CultASP3
        - CultASPI3
        - CultASP4
        - CultASPI4
        - CultASP
    - Cult water quant and timing
      - LMWtrQuantTime1
      - LMWtrQuantTimeI1
      - LMWtrQuantTime2
      - LMWtrQuantTimeI2
      - LMWtrQuantTime

#### groups

- LTW resilience final

#### dependency networks

- Apex predators
- Aspen forest
- Aspen habitat
- Big trees
- Burned high sev
- Burned low sev
- Burned mod sev
- CASPO territories
- Composition
- Conifer cover
- Cult water quant and timing
- Cultural resource quality
- Decomp funct redund
- Defense zone fire severity
- Early seral beta div
- Ecological function
- Extreme emission days
- Fine sediment load
- Flicker habitat
- Forest cover
- Functional fire
- Habitat for cultural keystone species
- Hardwood cover
- HCKS aspen habitat
- HCKS flicker habitat
- HCKS mountain quail habitat
- HCKS mule deer habitat
- Herbivore funct redund
- High elevation
- High elevation early
- High elevation late
- High elevation mid
- High emission days
- High sev patch
- Insectivore funct redund
- Late seral beta div
- Life and property
- Marten territories
- Mid elevation
- Mid elevation early
- Mid elevation late
- Mid elevation mid
- Mid seral beta div
- Moderate emission days
- Mtn quail habitat
- Mule deer habitat
- NOGO territories
- Percent land burned
- Phosphorus load
- Qual air extreme emission days
- Qual air high emission days
- Qual air mod emission days
- Qual air very high emission days
- Quality air
- Quality water
- Recreation
- Restoration of low intensity fire
- Scavenger funct redund
- Seed disp funct redund
- Seral stage
- Shrub cover
- Soil aerator funct redund
- Species diversity
- Species richness
- Sum extm emssn days
- Sum high emssn day
- Sum mod emmsn day
- Sum v high emssn day
- Summer extreme emission days
- Summer high emission days
- Summer moderate emission days
- Summer very high emission days
- Threat zone fire severity
- Upland vegetation health
- Very high emission days
- White pine forest
- Wildlife conservation
- WUI defense zone high severity
- WUI defense zone mod severity
- WUI fire
- WUI fire risk
- WUI threat zone high severity
- WUI threat zone mod severity
- Yellow pine forest
- \_Untitled1

#### data

- Aspen
- Aspen1
- Aspen2
- Aspen3
- Aspen4
- AspenI1
- AspenI2
- AspenI3
- AspenI4
- BigTrees
- BigTrees1
- BigTrees2
- BigTrees3
- BigTreesI1
- BigTreesI2
- BigTreesI3
- CASPO
- CASPO1
- CASPO2
- CASPOI1
- CASPOI2
- CultASP
- CultASP1
- CultASP2
- CultASP3
- CultASP4
- CultASPI1
- CultASPI2
- CultASPI3
- CultASPI4
- CultFire
- CultFire1
- CultFire2
- CultFire3
- CultFireI1
- CultFireI2
- CultFireI3
- Decomposer
- Decomposer1
- Decomposer2
- DecomposerI1
- DecomposerI2
- Disperser
- Disperser1
- Disperser2
- DisperserI1
- DisperserI2
- EarlyBeta
- EarlyBeta1
- EarlyBeta2
- EarlyBetaI1
- EarlyBetaI2
- ExtremeEmissionDays
- ExtremeEmissionDays1
- ExtremeEmissionDays2
- ExtremeEmissionDays3
- ExtremeEmissionDaysI1
- ExtremeEmissionDaysI2
- ExtremeEmissionDaysI3
- Flicker
- Flicker1
- Flicker2
- Flicker3
- FlickerI1
- FlickerI2
- FlickerI3
- Forest
- Forest1
- Forest2
- Forest3
- Forest4
- Forest5
- ForestI1
- ForestI2
- ForestI3
- ForestI4
- ForestI5
- Hardwood
- Hardwood1
- Hardwood2
- Hardwood3
- Hardwood4
- HardwoodI1
- HardwoodI2
- HardwoodI3
- HardwoodI4
- Herbivore
- Herbivore1
- Herbivore2
- HerbivoreI1
- HerbivoreI2
- HighEarly
- HighEarly1
- HighEarly2
- HighEarly3
- HighEarly4
- HighEarly5
- HighEarly6
- HighEarlyI1
- HighEarlyI2
- HighEarlyI3
- HighEarlyI4
- HighEarlyI5
- HighEarlyI6
- HighEmissionDays
- HighEmissionDays1
- HighEmissionDays2
- HighEmissionDays3
- HighEmissionDaysI1
- HighEmissionDaysI2
- HighEmissionDaysI3
- HighLate
- HighLate1
- HighLate2
- HighLate3
- HighLate4
- HighLate5
- HighLate6
- HighLateI1
- HighLateI2
- HighLateI3
- HighLateI4
- HighLateI5
- HighLateI6
- HighMid
- HighMid1
- HighMid2
- HighMid3
- HighMid4
- HighMid5
- HighMid6
- HighMidI1
- HighMidI2
- HighMidI3
- HighMidI4
- HighMidI5
- HighMidI6
- Insectivore
- Insectivore1
- Insectivore2
- InsectivoreI1
- InsectivoreI2
- LateBeta
- LateBeta1
- LateBeta2
- LateBetaI1
- LateBetaI2
- LMWtrQuantTime
- LMWtrQuantTime1
- LMWtrQuantTime2
- LMWtrQuantTimeI1
- LMWtrQuantTimeI2
- Marten
- Marten1
- Marten2
- MartenI1
- MartenI2
- MidBeta
- MidBeta1
- MidBeta2
- MidBetaI1
- MidBetaI2
- MidEarly
- MidEarly1
- MidEarly2
- MidEarly3
- MidEarly4
- MidEarly5
- MidEarly6
- MidEarlyI1
- MidEarlyI2
- MidEarlyI3
- MidEarlyI4
- MidEarlyI5
- MidEarlyI6
- MidLate
- MidLate1
- MidLate2
- MidLate3
- MidLate4
- MidLate5
- MidLate6
- MidLateI1
- MidLateI2
- MidLateI3
- MidLateI4
- MidLateI5
- MidLateI6
- MidMid
- MidMid1
- MidMid2
- MidMid3
- MidMid4
- MidMid5
- MidMid6
- MidMidI1
- MidMidI2
- MidMidI3
- MidMidI4
- MidMidI5
- MidMidI6
- ModEmissionDays
- ModEmissionDays1
- ModEmissionDays2
- ModEmissionDays3
- ModEmissionDaysI1
- ModEmissionDaysI2
- ModEmissionDaysI3
- MtnQuail
- MtnQuail1
- MtnQuail2
- MtnQuail3
- MtnQuailI1
- MtnQuailI2
- MtnQuailI3
- MuleDeer
- MuleDeer1
- MuleDeer2
- MuleDeer3
- MuleDeerI1
- MuleDeerI2
- MuleDeerI3
- NOGO
- NOGO1
- NOGO2
- NOGOI1
- NOGOI2
- PercentLandscapeHigh
- PercentLandscapeHigh1
- PercentLandscapeHigh2
- PercentLandscapeHigh3
- PercentLandscapeHigh4
- PercentLandscapeHighI1
- PercentLandscapeHighI2
- PercentLandscapeHighI3
- PercentLandscapeHighI4
- PercentLandscapeLow
- PercentLandscapeLow1
- PercentLandscapeLow2
- PercentLandscapeLow3
- PercentLandscapeLow4
- PercentLandscapeLow5
- PercentLandscapeLow6
- PercentLandscapeLowI1
- PercentLandscapeLowI2
- PercentLandscapeLowI3
- PercentLandscapeLowI4
- PercentLandscapeLowI5
- PercentLandscapeLowI6
- PercentLandscapeMod
- PercentLandscapeMod1
- PercentLandscapeMod2
- PercentLandscapeMod3
- PercentLandscapeMod4
- PercentLandscapeMod5
- PercentLandscapeModI1
- PercentLandscapeModI2
- PercentLandscapeModI3
- PercentLandscapeModI4
- PercentLandscapeModI5
- PerHSpatch
- PerHSpatch1
- PerHSpatch2
- PerHSpatchI1
- PerHSpatchI2
- Phosphorus
- Phosphorus1
- Phosphorus2
- PhosphorusI1
- PhosphorusI2
- Predator
- Predator1
- Predator2
- PredatorI1
- PredatorI2
- Sediment
- Sediment1
- Sediment2
- SedimentI1
- SedimentI2
- Shrub
- Shrub1
- Shrub2
- Shrub3
- Shrub4
- Shrub5
- ShrubI1
- ShrubI2
- ShrubI3
- ShrubI4
- ShrubI5
- SoilAerator
- SoilAerator1
- SoilAerator2
- SoilAeratorI1
- SoilAeratorI2
- SpecRich
- SpecRich1
- SpecRich2
- SpecRichI1
- SpecRichI2
- SumExtrmEmssnDays
- SumExtrmEmssnDays1
- SumExtrmEmssnDays2
- SumExtrmEmssnDays3
- SumExtrmEmssnDaysI1
- SumExtrmEmssnDaysI2
- SumExtrmEmssnDaysI3
- SumHighEmssnDays
- SumHighEmssnDays1
- SumHighEmssnDays2
- SumHighEmssnDays3
- SumHighEmssnDaysI1
- SumHighEmssnDaysI2
- SumHighEmssnDaysI3
- SumModEmssnDays
- SumModEmssnDays1
- SumModEmssnDays2
- SumModEmssnDays3
- SumModEmssnDaysI1
- SumModEmssnDaysI2
- SumModEmssnDaysI3
- SumVHighEmssnDays
- SumVHighEmssnDays1
- SumVHighEmssnDays2
- SumVHighEmssnDays3
- SumVHighEmssnDaysI1
- SumVHighEmssnDaysI2
- SumVHighEmssnDaysI3
- VHighEmissionDays
- VHighEmissionDays1
- VHighEmissionDays2
- VHighEmissionDays3
- VHighEmissionDaysI1
- VHighEmissionDaysI2
- VHighEmissionDaysI3
- WhitePine
- WhitePine1
- WhitePine2
- WhitePine3
- WhitePine4
- WhitePineI1
- WhitePineI2
- WhitePineI3
- WhitePineI4
- WUIDhigh
- WUIDhigh1
- WUIDhigh2
- WUIDhigh3
- WUIDhighI1
- WUIDhighI2
- WUIDhighI3
- WUIDmod
- WUIDmod1
- WUIDmod2
- WUIDmod3
- WUIDmodI1
- WUIDmodI2
- WUIDmodI3
- WUIThigh
- WUIThigh1
- WUIThigh2
- WUIThigh3
- WUIThigh4
- WUIThighI1
- WUIThighI2
- WUIThighI3
- WUIThighI4
- WUITmod
- WUITmod1
- WUITmod2
- WUITmod3
- WUITmod4
- WUITmodI1
- WUITmodI2
- WUITmodI3
- WUITmodI4
- YellowPine
- YellowPine1
- YellowPine2
- YellowPine3
- YellowPine4
- YellowPineI1
- YellowPineI2
- YellowPineI3
- YellowPineI4

#### data not in use

- HighEarlySeral
- HighLateSeral
- HighMidSeral
- MidEarlySeral
- MidLateSeral
- MidMidSeral
