## Appendix 3 for "Promoting long-term forest landscape resilience in the Lake Tahoe basin": jquery.cookie.js

```
/**
 * Cookie plugin
 *
 * Copyright (c) 2006 Klaus Hartl (stilbuero.de)
 * Dual licensed under the MIT and GPL licenses:
 * http://www.opensource.org/licenses/mit-license.php
 * http://www.gnu.org/licenses/gpl.html
 *
 */

/**
 * Create a cookie with the given name and value and other optional parameters.
 *
 * @example $.cookie('the_cookie', 'the_value');
 * @desc Set the value of a cookie.
 * @example $.cookie('the_cookie', 'the_value', {expires: 7, path: '/', domain: 'jquery.com', secure: true});
 * @desc Create a cookie with all available options.
 * @example $.cookie('the_cookie', 'the_value');
 * @desc Create a session cookie.
 * @example $.cookie('the_cookie', null);
 * @desc Delete a cookie by passing null as value.
 *
 * @param String name The name of the cookie.
 * @param String value The value of the cookie.
 * @param Object options An object literal containing key/value pairs to provide optional cookie attributes.
 * @option Number|Date expires Either an integer specifying the expiration date from now on in days or a Date object.
 *                             If a negative value is specified (e.g. a date in the past), the cookie will be deleted.
 *                             If set to null or omitted, the cookie will be a session cookie and will not be retained
 *                             when the the browser exits.
 * @option String path The value of the path atribute of the cookie (default: path of page that created the cookie).
 * @option String domain The value of the domain attribute of the cookie (default: domain of page that created the cookie).
 * @option Boolean secure If true, the secure attribute of the cookie will be set and the cookie transmission will
 *                        require a secure protocol (like HTTPS).
 * @type undefined
 *
 * @name $.cookie
 * @cat Plugins/Cookie
 * @author Klaus Hartl/
 */

/**
 * Get the value of a cookie with the given name.
 *
 * @example $.cookie('the_cookie');
 * @desc Get the value of a cookie.
 *
 * @param String name The name of the cookie.
 * @return The value of the cookie.
 * @type String
 *
 * @name $.cookie
 * @cat Plugins/Cookie
 * @author Klaus Hartl/
 */
jQuery.cookie = function(name, value, options) {
    if (typeof value != 'undefined') { // name and value given, set cookie
        options = options || {};
        if (value === null) {
            value = '';
            options.expires = -1;
        }
        var expires = '';
        if (options.expires && (typeof options.expires == 'number' || options.expires.toUTCString)) {
            var date;
            if (typeof options.expires == 'number') {
                date = new Date();
                date.setTime(date.getTime() + (options.expires * 24 * 60 * 60 * 1000));
            } else {
                date = options.expires;
            }
            expires = '; expires=' + date.toUTCString(); // use expires attribute, max-age is not supported by IE
        }
        var path = options.path ? '; path=' + options.path : '';
        var domain = options.domain ? '; domain=' + options.domain : '';
        var secure = options.secure ? '; secure' : '';
        document.cookie = [name, '=', encodeURIComponent(value), expires, path, domain, secure].join('');
    } else { // only name given, get cookie
        var cookieValue = null;
        if (document.cookie && document.cookie != '') {
            var cookies = document.cookie.split(';');
            for (var i = 0; i < cookies.length; i++) {
                var cookie = jQuery.trim(cookies[i]);
                // Does this cookie string begin with the name we want?
                if (cookie.substring(0, name.length + 1) == (name + '=')) {
                    cookieValue = decodeURIComponent(cookie.substring(name.length + 1));
                    break;
                }
            }
        }
        return cookieValue;
    }
};
```
