## Appendix 3 for "Promoting long-term forest landscape resilience in the Lake Tahoe basin": jquery.jHelperTip.1.0.min.js

```
/* Copyright (c) 2008 Kean Loong Tan http://www.gimiti.com/kltan
 * Licensed under the MIT (http://www.opensource.org/licenses/mit-license.php)
 * Copyright notice and license must remain intact for legal use
 * jHelpertip
 * Version: 1.0 (Jun 2, 2008)
 * Requires: jQuery 1.2+
 */
(function(A){A.fn.jHelperTip=function(C){var E=A.extend({},A.fn.jHelperTip.defaults,C);if(A(E.ttC).length==0){A('<div id="'+E.ttC.slice(1)+'"></div>').appendTo("body")}if(A(E.dC).length==0){A('<div id="'+E.dC.slice(1)+'"></div>').appendTo("body")}if(A(E.aC).length==0){A('<div id="'+E.aC.slice(1)+'"></div>').appendTo("body")}A(E.ttC).add(E.aC).css({position:"absolute",display:"inline"}).hide();A(E.dC).hide();var F=function(){if(E.source=="attribute"){A(E.aC).hide().empty()}else{A(E.ttC).hide().empty()}};A(".jHelperTipClose").bind("click",F);A(E.ttC).bind("mouseover",function(){A(E.ttC).show();return false});var B=function(H,G){if(E.source=="ajax"){D(G);A(E.ttC).html('<div><img src="'+E.loadingImg+'"/> '+E.loadingText+"</div>").show();A.ajax({type:E.type,url:E.url,data:E.data,success:function(I){A(E.ttC).html(I);A(".jHelperTipClose").unbind("click",F);A(".jHelperTipClose").bind("click",F)}})}else{if(E.source=="container"){A(E.ttC).show().empty();A(E.dC).clone(true).show().appendTo(E.ttC)}}if(E.source=="attribute"){A(E.aC).html(A(H).attr(E.attrName))}};var D=function(I){var H=I.pageY+E.topOff;var G=I.pageX+E.leftOff;if(E.source=="attribute"){A(E.aC).css({top:H,left:G,opacity:E.opacity}).show()}else{A(E.ttC).css({top:H,left:G,opacity:E.opacity}).show()}};if(E.trigger=="hover"){A(this).bind("mouseover",function(G){G.preventDefault();B(this,G);return false});A(this).bind("mousemove",function(G){D(G);return false});A(this).bind("mouseout",function(G){if(E.source=="attribute"){A(E.aC).hide().empty()}else{A(E.ttC).hide().empty()}return false})}else{if(E.trigger=="click"){A(this).bind("click",function(G){B(this,G);D(G);A(document).bind("click",function(H){if(E.autoClose){if(E.source=="attribute"){A(E.aC).hide().empty()}else{A(E.ttC).hide().empty()}}});return false})}}};A.fn.jHelperTip.defaults={trigger:"click",topOff:3,leftOff:10,source:"container",attrName:"",ttC:"#jHelperTipContainer",dC:"#jHelperTipDataContainer",aC:"#jHelperTipAttrContainer",opacity:1,loadingImg:"ajax-loader.gif",loadingText:"Loading...",type:"GET",autoClose:true}})(jQuery);
```
