## Appendix 3 for "Promoting long-term forest landscape resilience in the Lake Tahoe basin": jquery.pack.js

```
/*
 * jQuery 1.2.6 - New Wave Javascript
 *
 * Copyright (c) 2008 John Resig (jquery.com)
 * Dual licensed under the MIT (MIT-LICENSE.txt)
 * and GPL (GPL-LICENSE.txt) licenses.
 *
 * $Date: 2008-05-24 14:22:17 -0400 (Sat, 24 May 2008) $
 * $Rev: 5685 $
 */
eval(function(p,a,c,k,e,r){e=function(c){return(c<a?'':e(parseInt(c/a)))+((c=c%a)>35?String.fromCharCode(c+29):c.toString(36))};if(!''.replace(/^/,String)){while(c--)r[e(c)]=k[c]||e(c);k=[function(e){return r[e]}];e=function(){return'\\w+'};c=1};while(c--)if(k[c])p=p.replace(new RegExp('\\b'+e(c)+'\\b','g'),k[c]);return p}('(H(){J w=1b.4M,3m$=1b.$;J D=1b.4M=1b.$=H(a,b){I 2B D.17.5j(a,b)};J u=/^[^<]*(<(.|\\s)+>)[^>]*$|^#(\\w+)$/,62=/^.[^:#\\[\\.]*$/,12;D.17=D.44={5j:H(d,b){d=d||S;G(d.16){7[0]=d;7.K=1;I 7}G(1j d=="23"){J c=u.2D(d);G(c&&(c[1]||!b)){G(c[1])d=D.4h([c[1]],b);N{J a=S.61(c[3]);G(a){G(a.2v!=c[3])I D().2q(d);I D(a)}d=[]}}N I D(b).2q(d)}N G(D.1D(d))I D(S)[D.17.27?"27":"43"](d);I 7.6Y(D.2d(d))},5w:"1.2.6",8G:H(){I 7.K},K:0,3p:H(a){I a==12?D.2d(7):7[a]},2I:H(b){J a=D(b);a.5n=7;I a},6Y:H(a){7.K=0;2p.44.1p.1w(7,a);I 7},P:H(a,b){I D.P(7,a,b)},5i:H(b){J a=-1;I D.2L(b&&b.5w?b[0]:b,7)},1K:H(c,a,b){J d=c;G(c.1q==56)G(a===12)I 7[0]&&D[b||"1K"](7[0],c);N{d={};d[c]=a}I 7.P(H(i){R(c 1n d)D.1K(b?7.V:7,c,D.1i(7,d[c],b,i,c))})},1g:H(b,a){G((b==\'2h\'||b==\'1Z\')&&3d(a)<0)a=12;I 7.1K(b,a,"2a")},1r:H(b){G(1j b!="49"&&b!=U)I 7.4E().3v((7[0]&&7[0].2z||S).5F(b));J a="";D.P(b||7,H(){D.P(7.3t,H(){G(7.16!=8)a+=7.16!=1?7.76:D.17.1r([7])})});I a},5z:H(b){G(7[0])D(b,7[0].2z).5y().39(7[0]).2l(H(){J a=7;1B(a.1x)a=a.1x;I a}).3v(7);I 7},8Y:H(a){I 7.P(H(){D(7).6Q().5z(a)})},8R:H(a){I 7.P(H(){D(7).5z(a)})},3v:H(){I 7.3W(19,M,Q,H(a){G(7.16==1)7.3U(a)})},6F:H(){I 7.3W(19,M,M,H(a){G(7.16==1)7.39(a,7.1x)})},6E:H(){I 7.3W(19,Q,Q,H(a){7.1d.39(a,7)})},5q:H(){I 7.3W(19,Q,M,H(a){7.1d.39(a,7.2H)})},3l:H(){I 7.5n||D([])},2q:H(b){J c=D.2l(7,H(a){I D.2q(b,a)});I 7.2I(/[^+>] [^+>]/.11(b)||b.1h("..")>-1?D.4r(c):c)},5y:H(e){J f=7.2l(H(){G(D.14.1f&&!D.4n(7)){J a=7.6o(M),5h=S.3h("1v");5h.3U(a);I D.4h([5h.4H])[0]}N I 7.6o(M)});J d=f.2q("*").5c().P(H(){G(7[E]!=12)7[E]=U});G(e===M)7.2q("*").5c().P(H(i){G(7.16==3)I;J c=D.L(7,"3w");R(J a 1n c)R(J b 1n c[a])D.W.1e(d[i],a,c[a][b],c[a][b].L)});I f},1E:H(b){I 7.2I(D.1D(b)&&D.3C(7,H(a,i){I b.1k(a,i)})||D.3g(b,7))},4Y:H(b){G(b.1q==56)G(62.11(b))I 7.2I(D.3g(b,7,M));N b=D.3g(b,7);J a=b.K&&b[b.K-1]!==12&&!b.16;I 7.1E(H(){I a?D.2L(7,b)<0:7!=b})},1e:H(a){I 7.2I(D.4r(D.2R(7.3p(),1j a==\'23\'?D(a):D.2d(a))))},3F:H(a){I!!a&&D.3g(a,7).K>0},7T:H(a){I 7.3F("."+a)},6e:H(b){G(b==12){G(7.K){J c=7[0];G(D.Y(c,"2A")){J e=c.64,63=[],15=c.15,2V=c.O=="2A-2V";G(e<0)I U;R(J i=2V?e:0,2f=2V?e+1:15.K;i<2f;i++){J d=15[i];G(d.2W){b=D.14.1f&&!d.at.2x.an?d.1r:d.2x;G(2V)I b;63.1p(b)}}I 63}N I(7[0].2x||"").1o(/\\r/g,"")}I 12}G(b.1q==4L)b+=\'\';I 7.P(H(){G(7.16!=1)I;G(b.1q==2p&&/5O|5L/.11(7.O))7.4J=(D.2L(7.2x,b)>=0||D.2L(7.34,b)>=0);N G(D.Y(7,"2A")){J a=D.2d(b);D("9R",7).P(H(){7.2W=(D.2L(7.2x,a)>=0||D.2L(7.1r,a)>=0)});G(!a.K)7.64=-1}N 7.2x=b})},2K:H(a){I a==12?(7[0]?7[0].4H:U):7.4E().3v(a)},7b:H(a){I 7.5q(a).21()},79:H(i){I 7.3s(i,i+1)},3s:H(){I 7.2I(2p.44.3s.1w(7,19))},2l:H(b){I 7.2I(D.2l(7,H(a,i){I b.1k(a,i,a)}))},5c:H(){I 7.1e(7.5n)},L:H(d,b){J a=d.1R(".");a[1]=a[1]?"."+a[1]:"";G(b===12){J c=7.5C("9z"+a[1]+"!",[a[0]]);G(c===12&&7.K)c=D.L(7[0],d);I c===12&&a[1]?7.L(a[0]):c}N I 7.1P("9u"+a[1]+"!",[a[0],b]).P(H(){D.L(7,d,b)})},3b:H(a){I 7.P(H(){D.3b(7,a)})},3W:H(g,f,h,d){J e=7.K>1,3x;I 7.P(H(){G(!3x){3x=D.4h(g,7.2z);G(h)3x.9o()}J b=7;G(f&&D.Y(7,"1T")&&D.Y(3x[0],"4F"))b=7.3H("22")[0]||7.3U(7.2z.3h("22"));J c=D([]);D.P(3x,H(){J a=e?D(7).5y(M)[0]:7;G(D.Y(a,"1m"))c=c.1e(a);N{G(a.16==1)c=c.1e(D("1m",a).21());d.1k(b,a)}});c.P(6T)})}};D.17.5j.44=D.17;H 6T(i,a){G(a.4d)D.3Y({1a:a.4d,31:Q,1O:"1m"});N D.5u(a.1r||a.6O||a.4H||"");G(a.1d)a.1d.37(a)}H 1z(){I+2B 8J}D.1l=D.17.1l=H(){J b=19[0]||{},i=1,K=19.K,4x=Q,15;G(b.1q==8I){4x=b;b=19[1]||{};i=2}G(1j b!="49"&&1j b!="H")b={};G(K==i){b=7;--i}R(;i<K;i++)G((15=19[i])!=U)R(J c 1n 15){J a=b[c],2w=15[c];G(b===2w)6M;G(4x&&2w&&1j 2w=="49"&&!2w.16)b[c]=D.1l(4x,a||(2w.K!=U?[]:{}),2w);N G(2w!==12)b[c]=2w}I b};J E="4M"+1z(),6K=0,5r={},6G=/z-?5i|8B-?8A|1y|6B|8v-?1Z/i,3P=S.3P||{};D.1l({8u:H(a){1b.$=3m$;G(a)1b.4M=w;I D},1D:H(a){I!!a&&1j a!="23"&&!a.Y&&a.1q!=2p&&/^[\\s[]?H/.11(a+"")},4n:H(a){I a.1C&&!a.1c||a.2j&&a.2z&&!a.2z.1c},5u:H(a){a=D.3k(a);G(a){J b=S.3H("6w")[0]||S.1C,1m=S.3h("1m");1m.O="1r/4t";G(D.14.1f)1m.1r=a;N 1m.3U(S.5F(a));b.39(1m,b.1x);b.37(1m)}},Y:H(b,a){I b.Y&&b.Y.2r()==a.2r()},1Y:{},L:H(c,d,b){c=c==1b?5r:c;J a=c[E];G(!a)a=c[E]=++6K;G(d&&!D.1Y[a])D.1Y[a]={};G(b!==12)D.1Y[a][d]=b;I d?D.1Y[a][d]:a},3b:H(c,b){c=c==1b?5r:c;J a=c[E];G(b){G(D.1Y[a]){2U D.1Y[a][b];b="";R(b 1n D.1Y[a])1X;G(!b)D.3b(c)}}N{1U{2U c[E]}1V(e){G(c.5l)c.5l(E)}2U D.1Y[a]}},P:H(d,a,c){J e,i=0,K=d.K;G(c){G(K==12){R(e 1n d)G(a.1w(d[e],c)===Q)1X}N R(;i<K;)G(a.1w(d[i++],c)===Q)1X}N{G(K==12){R(e 1n d)G(a.1k(d[e],e,d[e])===Q)1X}N R(J b=d[0];i<K&&a.1k(b,i,b)!==Q;b=d[++i]){}}I d},1i:H(b,a,c,i,d){G(D.1D(a))a=a.1k(b,i);I a&&a.1q==4L&&c=="2a"&&!6G.11(d)?a+"2X":a},1F:{1e:H(c,b){D.P((b||"").1R(/\\s+/),H(i,a){G(c.16==1&&!D.1F.3T(c.1F,a))c.1F+=(c.1F?" ":"")+a})},21:H(c,b){G(c.16==1)c.1F=b!=12?D.3C(c.1F.1R(/\\s+/),H(a){I!D.1F.3T(b,a)}).6s(" "):""},3T:H(b,a){I D.2L(a,(b.1F||b).6r().1R(/\\s+/))>-1}},6q:H(b,c,a){J e={};R(J d 1n c){e[d]=b.V[d];b.V[d]=c[d]}a.1k(b);R(J d 1n c)b.V[d]=e[d]},1g:H(d,e,c){G(e=="2h"||e=="1Z"){J b,3X={30:"5x",5g:"1G",18:"3I"},35=e=="2h"?["5e","6k"]:["5G","6i"];H 5b(){b=e=="2h"?d.8f:d.8c;J a=0,2C=0;D.P(35,H(){a+=3d(D.2a(d,"57"+7,M))||0;2C+=3d(D.2a(d,"2C"+7+"4b",M))||0});b-=29.83(a+2C)}G(D(d).3F(":4j"))5b();N D.6q(d,3X,5b);I 29.2f(0,b)}I D.2a(d,e,c)},2a:H(f,l,k){J e,V=f.V;H 3E(b){G(!D.14.2k)I Q;J a=3P.54(b,U);I!a||a.52("3E")==""}G(l=="1y"&&D.14.1f){e=D.1K(V,"1y");I e==""?"1":e}G(D.14.2G&&l=="18"){J d=V.50;V.50="0 7Y 7W";V.50=d}G(l.1I(/4i/i))l=y;G(!k&&V&&V[l])e=V[l];N G(3P.54){G(l.1I(/4i/i))l="4i";l=l.1o(/([A-Z])/g,"-$1").3y();J c=3P.54(f,U);G(c&&!3E(f))e=c.52(l);N{J g=[],2E=[],a=f,i=0;R(;a&&3E(a);a=a.1d)2E.6h(a);R(;i<2E.K;i++)G(3E(2E[i])){g[i]=2E[i].V.18;2E[i].V.18="3I"}e=l=="18"&&g[2E.K-1]!=U?"2F":(c&&c.52(l))||"";R(i=0;i<g.K;i++)G(g[i]!=U)2E[i].V.18=g[i]}G(l=="1y"&&e=="")e="1"}N G(f.4g){J h=l.1o(/\\-(\\w)/g,H(a,b){I b.2r()});e=f.4g[l]||f.4g[h];G(!/^\\d+(2X)?$/i.11(e)&&/^\\d/.11(e)){J j=V.1A,66=f.65.1A;f.65.1A=f.4g.1A;V.1A=e||0;e=V.aM+"2X";V.1A=j;f.65.1A=66}}I e},4h:H(l,h){J k=[];h=h||S;G(1j h.3h==\'12\')h=h.2z||h[0]&&h[0].2z||S;D.P(l,H(i,d){G(!d)I;G(d.1q==4L)d+=\'\';G(1j d=="23"){d=d.1o(/(<(\\w+)[^>]*?)\\/>/g,H(b,a,c){I c.1I(/^(aK|4f|7E|aG|4T|7A|aB|3n|az|ay|av)$/i)?b:a+"></"+c+">"});J f=D.3k(d).3y(),1v=h.3h("1v");J e=!f.1h("<au")&&[1,"<2A 7w=\'7w\'>","</2A>"]||!f.1h("<ar")&&[1,"<7v>","</7v>"]||f.1I(/^<(aq|22|am|ak|ai)/)&&[1,"<1T>","</1T>"]||!f.1h("<4F")&&[2,"<1T><22>","</22></1T>"]||(!f.1h("<af")||!f.1h("<ad"))&&[3,"<1T><22><4F>","</4F></22></1T>"]||!f.1h("<7E")&&[2,"<1T><22></22><7q>","</7q></1T>"]||D.14.1f&&[1,"1v<1v>","</1v>"]||[0,"",""];1v.4H=e[1]+d+e[2];1B(e[0]--)1v=1v.5T;G(D.14.1f){J g=!f.1h("<1T")&&f.1h("<22")<0?1v.1x&&1v.1x.3t:e[1]=="<1T>"&&f.1h("<22")<0?1v.3t:[];R(J j=g.K-1;j>=0;--j)G(D.Y(g[j],"22")&&!g[j].3t.K)g[j].1d.37(g[j]);G(/^\\s/.11(d))1v.39(h.5F(d.1I(/^\\s*/)[0]),1v.1x)}d=D.2d(1v.3t)}G(d.K===0&&(!D.Y(d,"3V")&&!D.Y(d,"2A")))I;G(d[0]==12||D.Y(d,"3V")||d.15)k.1p(d);N k=D.2R(k,d)});I k},1K:H(d,f,c){G(!d||d.16==3||d.16==8)I 12;J e=!D.4n(d),40=c!==12,1f=D.14.1f;f=e&&D.3X[f]||f;G(d.2j){J g=/5Q|4d|V/.11(f);G(f=="2W"&&D.14.2k)d.1d.64;G(f 1n d&&e&&!g){G(40){G(f=="O"&&D.Y(d,"4T")&&d.1d)7p"O a3 a1\'t 9V 9U";d[f]=c}G(D.Y(d,"3V")&&d.7i(f))I d.7i(f).76;I d[f]}G(1f&&e&&f=="V")I D.1K(d.V,"9T",c);G(40)d.9Q(f,""+c);J h=1f&&e&&g?d.4G(f,2):d.4G(f);I h===U?12:h}G(1f&&f=="1y"){G(40){d.6B=1;d.1E=(d.1E||"").1o(/7f\\([^)]*\\)/,"")+(3r(c)+\'\'=="9L"?"":"7f(1y="+c*7a+")")}I d.1E&&d.1E.1h("1y=")>=0?(3d(d.1E.1I(/1y=([^)]*)/)[1])/7a)+\'\':""}f=f.1o(/-([a-z])/9H,H(a,b){I b.2r()});G(40)d[f]=c;I d[f]},3k:H(a){I(a||"").1o(/^\\s+|\\s+$/g,"")},2d:H(b){J a=[];G(b!=U){J i=b.K;G(i==U||b.1R||b.4I||b.1k)a[0]=b;N 1B(i)a[--i]=b[i]}I a},2L:H(b,a){R(J i=0,K=a.K;i<K;i++)G(a[i]===b)I i;I-1},2R:H(a,b){J i=0,T,2S=a.K;G(D.14.1f){1B(T=b[i++])G(T.16!=8)a[2S++]=T}N 1B(T=b[i++])a[2S++]=T;I a},4r:H(a){J c=[],2o={};1U{R(J i=0,K=a.K;i<K;i++){J b=D.L(a[i]);G(!2o[b]){2o[b]=M;c.1p(a[i])}}}1V(e){c=a}I c},3C:H(c,a,d){J b=[];R(J i=0,K=c.K;i<K;i++)G(!d!=!a(c[i],i))b.1p(c[i]);I b},2l:H(d,a){J c=[];R(J i=0,K=d.K;i<K;i++){J b=a(d[i],i);G(b!=U)c[c.K]=b}I c.7d.1w([],c)}});J v=9B.9A.3y();D.14={5B:(v.1I(/.+(?:9y|9x|9w|9v)[\\/: ]([\\d.]+)/)||[])[1],2k:/75/.11(v),2G:/2G/.11(v),1f:/1f/.11(v)&&!/2G/.11(v),42:/42/.11(v)&&!/(9s|75)/.11(v)};J y=D.14.1f?"7o":"72";D.1l({71:!D.14.1f||S.70=="6Z",3X:{"R":"9n","9k":"1F","4i":y,72:y,7o:y,9h:"9f",9e:"9d",9b:"99"}});D.P({6W:H(a){I a.1d},97:H(a){I D.4S(a,"1d")},95:H(a){I D.3a(a,2,"2H")},91:H(a){I D.3a(a,2,"4l")},8Z:H(a){I D.4S(a,"2H")},8X:H(a){I D.4S(a,"4l")},8W:H(a){I D.5v(a.1d.1x,a)},8V:H(a){I D.5v(a.1x)},6Q:H(a){I D.Y(a,"8U")?a.8T||a.8S.S:D.2d(a.3t)}},H(c,d){D.17[c]=H(b){J a=D.2l(7,d);G(b&&1j b=="23")a=D.3g(b,a);I 7.2I(D.4r(a))}});D.P({6P:"3v",8Q:"6F",39:"6E",8P:"5q",8O:"7b"},H(c,b){D.17[c]=H(){J a=19;I 7.P(H(){R(J i=0,K=a.K;i<K;i++)D(a[i])[b](7)})}});D.P({8N:H(a){D.1K(7,a,"");G(7.16==1)7.5l(a)},8M:H(a){D.1F.1e(7,a)},8L:H(a){D.1F.21(7,a)},8K:H(a){D.1F[D.1F.3T(7,a)?"21":"1e"](7,a)},21:H(a){G(!a||D.1E(a,[7]).r.K){D("*",7).1e(7).P(H(){D.W.21(7);D.3b(7)});G(7.1d)7.1d.37(7)}},4E:H(){D(">*",7).21();1B(7.1x)7.37(7.1x)}},H(a,b){D.17[a]=H(){I 7.P(b,19)}});D.P(["6N","4b"],H(i,c){J b=c.3y();D.17[b]=H(a){I 7[0]==1b?D.14.2G&&S.1c["5t"+c]||D.14.2k&&1b["5s"+c]||S.70=="6Z"&&S.1C["5t"+c]||S.1c["5t"+c]:7[0]==S?29.2f(29.2f(S.1c["4y"+c],S.1C["4y"+c]),29.2f(S.1c["2i"+c],S.1C["2i"+c])):a==12?(7.K?D.1g(7[0],b):U):7.1g(b,a.1q==56?a:a+"2X")}});H 25(a,b){I a[0]&&3r(D.2a(a[0],b,M),10)||0}J C=D.14.2k&&3r(D.14.5B)<8H?"(?:[\\\\w*3m-]|\\\\\\\\.)":"(?:[\\\\w\\8F-\\8E*3m-]|\\\\\\\\.)",6L=2B 4v("^>\\\\s*("+C+"+)"),6J=2B 4v("^("+C+"+)(#)("+C+"+)"),6I=2B 4v("^([#.]?)("+C+"*)");D.1l({6H:{"":H(a,i,m){I m[2]=="*"||D.Y(a,m[2])},"#":H(a,i,m){I a.4G("2v")==m[2]},":":{8D:H(a,i,m){I i<m[3]-0},8C:H(a,i,m){I i>m[3]-0},3a:H(a,i,m){I m[3]-0==i},79:H(a,i,m){I m[3]-0==i},3o:H(a,i){I i==0},3S:H(a,i,m,r){I i==r.K-1},6D:H(a,i){I i%2==0},6C:H(a,i){I i%2},"3o-4u":H(a){I a.1d.3H("*")[0]==a},"3S-4u":H(a){I D.3a(a.1d.5T,1,"4l")==a},"8z-4u":H(a){I!D.3a(a.1d.5T,2,"4l")},6W:H(a){I a.1x},4E:H(a){I!a.1x},8y:H(a,i,m){I(a.6O||a.8x||D(a).1r()||"").1h(m[3])>=0},4j:H(a){I"1G"!=a.O&&D.1g(a,"18")!="2F"&&D.1g(a,"5g")!="1G"},1G:H(a){I"1G"==a.O||D.1g(a,"18")=="2F"||D.1g(a,"5g")=="1G"},8w:H(a){I!a.3R},3R:H(a){I a.3R},4J:H(a){I a.4J},2W:H(a){I a.2W||D.1K(a,"2W")},1r:H(a){I"1r"==a.O},5O:H(a){I"5O"==a.O},5L:H(a){I"5L"==a.O},5p:H(a){I"5p"==a.O},3Q:H(a){I"3Q"==a.O},5o:H(a){I"5o"==a.O},6A:H(a){I"6A"==a.O},6z:H(a){I"6z"==a.O},2s:H(a){I"2s"==a.O||D.Y(a,"2s")},4T:H(a){I/4T|2A|6y|2s/i.11(a.Y)},3T:H(a,i,m){I D.2q(m[3],a).K},8t:H(a){I/h\\d/i.11(a.Y)},8s:H(a){I D.3C(D.3O,H(b){I a==b.T}).K}}},6x:[/^(\\[) *@?([\\w-]+) *([!*$^~=]*) *(\'?"?)(.*?)\\4 *\\]/,/^(:)([\\w-]+)\\("?\'?(.*?(\\(.*?\\))?[^(]*?)"?\'?\\)/,2B 4v("^([:.#]*)("+C+"+)")],3g:H(a,c,b){J d,1t=[];1B(a&&a!=d){d=a;J f=D.1E(a,c,b);a=f.t.1o(/^\\s*,\\s*/,"");1t=b?c=f.r:D.2R(1t,f.r)}I 1t},2q:H(t,o){G(1j t!="23")I[t];G(o&&o.16!=1&&o.16!=9)I[];o=o||S;J d=[o],2o=[],3S,Y;1B(t&&3S!=t){J r=[];3S=t;t=D.3k(t);J l=Q,3j=6L,m=3j.2D(t);G(m){Y=m[1].2r();R(J i=0;d[i];i++)R(J c=d[i].1x;c;c=c.2H)G(c.16==1&&(Y=="*"||c.Y.2r()==Y))r.1p(c);d=r;t=t.1o(3j,"");G(t.1h(" ")==0)6M;l=M}N{3j=/^([>+~])\\s*(\\w*)/i;G((m=3j.2D(t))!=U){r=[];J k={};Y=m[2].2r();m=m[1];R(J j=0,3i=d.K;j<3i;j++){J n=m=="~"||m=="+"?d[j].2H:d[j].1x;R(;n;n=n.2H)G(n.16==1){J g=D.L(n);G(m=="~"&&k[g])1X;G(!Y||n.Y.2r()==Y){G(m=="~")k[g]=M;r.1p(n)}G(m=="+")1X}}d=r;t=D.3k(t.1o(3j,""));l=M}}G(t&&!l){G(!t.1h(",")){G(o==d[0])d.4s();2o=D.2R(2o,d);r=d=[o];t=" "+t.6v(1,t.K)}N{J h=6J;J m=h.2D(t);G(m){m=[0,m[2],m[3],m[1]]}N{h=6I;m=h.2D(t)}m[2]=m[2].1o(/\\\\/g,"");J f=d[d.K-1];G(m[1]=="#"&&f&&f.61&&!D.4n(f)){J p=f.61(m[2]);G((D.14.1f||D.14.2G)&&p&&1j p.2v=="23"&&p.2v!=m[2])p=D(\'[@2v="\'+m[2]+\'"]\',f)[0];d=r=p&&(!m[3]||D.Y(p,m[3]))?[p]:[]}N{R(J i=0;d[i];i++){J a=m[1]=="#"&&m[3]?m[3]:m[1]!=""||m[0]==""?"*":m[2];G(a=="*"&&d[i].Y.3y()=="49")a="3n";r=D.2R(r,d[i].3H(a))}G(m[1]==".")r=D.5m(r,m[2]);G(m[1]=="#"){J e=[];R(J i=0;r[i];i++)G(r[i].4G("2v")==m[2]){e=[r[i]];1X}r=e}d=r}t=t.1o(h,"")}}G(t){J b=D.1E(t,r);d=r=b.r;t=D.3k(b.t)}}G(t)d=[];G(d&&o==d[0])d.4s();2o=D.2R(2o,d);I 2o},5m:H(r,m,a){m=" "+m+" ";J c=[];R(J i=0;r[i];i++){J b=(" "+r[i].1F+" ").1h(m)>=0;G(!a&&b||a&&!b)c.1p(r[i])}I c},1E:H(t,r,h){J d;1B(t&&t!=d){d=t;J p=D.6x,m;R(J i=0;p[i];i++){m=p[i].2D(t);G(m){t=t.8r(m[0].K);m[2]=m[2].1o(/\\\\/g,"");1X}}G(!m)1X;G(m[1]==":"&&m[2]=="4Y")r=62.11(m[3])?D.1E(m[3],r,M).r:D(r).4Y(m[3]);N G(m[1]==".")r=D.5m(r,m[2],h);N G(m[1]=="["){J g=[],O=m[3];R(J i=0,3i=r.K;i<3i;i++){J a=r[i],z=a[D.3X[m[2]]||m[2]];G(z==U||/5Q|4d|2W/.11(m[2]))z=D.1K(a,m[2])||\'\';G((O==""&&!!z||O=="="&&z==m[5]||O=="!="&&z!=m[5]||O=="^="&&z&&!z.1h(m[5])||O=="$="&&z.6v(z.K-m[5].K)==m[5]||(O=="*="||O=="~=")&&z.1h(m[5])>=0)^h)g.1p(a)}r=g}N G(m[1]==":"&&m[2]=="3a-4u"){J e={},g=[],11=/(-?)(\\d*)n((?:\\+|-)?\\d*)/.2D(m[3]=="6D"&&"2n"||m[3]=="6C"&&"2n+1"||!/\\D/.11(m[3])&&"8q+"+m[3]||m[3]),3o=(11[1]+(11[2]||1))-0,d=11[3]-0;R(J i=0,3i=r.K;i<3i;i++){J j=r[i],1d=j.1d,2v=D.L(1d);G(!e[2v]){J c=1;R(J n=1d.1x;n;n=n.2H)G(n.16==1)n.4q=c++;e[2v]=M}J b=Q;G(3o==0){G(j.4q==d)b=M}N G((j.4q-d)%3o==0&&(j.4q-d)/3o>=0)b=M;G(b^h)g.1p(j)}r=g}N{J f=D.6H[m[1]];G(1j f=="49")f=f[m[2]];G(1j f=="23")f=6u("Q||H(a,i){I "+f+";}");r=D.3C(r,H(a,i){I f(a,i,m,r)},h)}}I{r:r,t:t}},4S:H(b,c){J a=[],1t=b[c];1B(1t&&1t!=S){G(1t.16==1)a.1p(1t);1t=1t[c]}I a},3a:H(a,e,c,b){e=e||1;J d=0;R(;a;a=a[c])G(a.16==1&&++d==e)1X;I a},5v:H(n,a){J r=[];R(;n;n=n.2H){G(n.16==1&&n!=a)r.1p(n)}I r}});D.W={1e:H(f,i,g,e){G(f.16==3||f.16==8)I;G(D.14.1f&&f.4I)f=1b;G(!g.24)g.24=7.24++;G(e!=12){J h=g;g=7.3M(h,H(){I h.1w(7,19)});g.L=e}J j=D.L(f,"3w")||D.L(f,"3w",{}),1H=D.L(f,"1H")||D.L(f,"1H",H(){G(1j D!="12"&&!D.W.5k)I D.W.1H.1w(19.3L.T,19)});1H.T=f;D.P(i.1R(/\\s+/),H(c,b){J a=b.1R(".");b=a[0];g.O=a[1];J d=j[b];G(!d){d=j[b]={};G(!D.W.2t[b]||D.W.2t[b].4p.1k(f)===Q){G(f.3K)f.3K(b,1H,Q);N G(f.6t)f.6t("4o"+b,1H)}}d[g.24]=g;D.W.26[b]=M});f=U},24:1,26:{},21:H(e,h,f){G(e.16==3||e.16==8)I;J i=D.L(e,"3w"),1L,5i;G(i){G(h==12||(1j h=="23"&&h.8p(0)=="."))R(J g 1n i)7.21(e,g+(h||""));N{G(h.O){f=h.2y;h=h.O}D.P(h.1R(/\\s+/),H(b,a){J c=a.1R(".");a=c[0];G(i[a]){G(f)2U i[a][f.24];N R(f 1n i[a])G(!c[1]||i[a][f].O==c[1])2U i[a][f];R(1L 1n i[a])1X;G(!1L){G(!D.W.2t[a]||D.W.2t[a].4A.1k(e)===Q){G(e.6p)e.6p(a,D.L(e,"1H"),Q);N G(e.6n)e.6n("4o"+a,D.L(e,"1H"))}1L=U;2U i[a]}}})}R(1L 1n i)1X;G(!1L){J d=D.L(e,"1H");G(d)d.T=U;D.3b(e,"3w");D.3b(e,"1H")}}},1P:H(h,c,f,g,i){c=D.2d(c);G(h.1h("!")>=0){h=h.3s(0,-1);J a=M}G(!f){G(7.26[h])D("*").1e([1b,S]).1P(h,c)}N{G(f.16==3||f.16==8)I 12;J b,1L,17=D.1D(f[h]||U),W=!c[0]||!c[0].32;G(W){c.6h({O:h,2J:f,32:H(){},3J:H(){},4C:1z()});c[0][E]=M}c[0].O=h;G(a)c[0].6m=M;J d=D.L(f,"1H");G(d)b=d.1w(f,c);G((!17||(D.Y(f,\'a\')&&h=="4V"))&&f["4o"+h]&&f["4o"+h].1w(f,c)===Q)b=Q;G(W)c.4s();G(i&&D.1D(i)){1L=i.1w(f,b==U?c:c.7d(b));G(1L!==12)b=1L}G(17&&g!==Q&&b!==Q&&!(D.Y(f,\'a\')&&h=="4V")){7.5k=M;1U{f[h]()}1V(e){}}7.5k=Q}I b},1H:H(b){J a,1L,38,5f,4m;b=19[0]=D.W.6l(b||1b.W);38=b.O.1R(".");b.O=38[0];38=38[1];5f=!38&&!b.6m;4m=(D.L(7,"3w")||{})[b.O];R(J j 1n 4m){J c=4m[j];G(5f||c.O==38){b.2y=c;b.L=c.L;1L=c.1w(7,19);G(a!==Q)a=1L;G(1L===Q){b.32();b.3J()}}}I a},6l:H(b){G(b[E]==M)I b;J d=b;b={8o:d};J c="8n 8m 8l 8k 2s 8j 47 5d 6j 5E 8i L 8h 8g 4K 2y 5a 59 8e 8b 58 6f 8a 88 4k 87 86 84 6d 2J 4C 6c O 82 81 35".1R(" ");R(J i=c.K;i;i--)b[c[i]]=d[c[i]];b[E]=M;b.32=H(){G(d.32)d.32();d.80=Q};b.3J=H(){G(d.3J)d.3J();d.7Z=M};b.4C=b.4C||1z();G(!b.2J)b.2J=b.6d||S;G(b.2J.16==3)b.2J=b.2J.1d;G(!b.4k&&b.4K)b.4k=b.4K==b.2J?b.6c:b.4K;G(b.58==U&&b.5d!=U){J a=S.1C,1c=S.1c;b.58=b.5d+(a&&a.2e||1c&&1c.2e||0)-(a.6b||0);b.6f=b.6j+(a&&a.2c||1c&&1c.2c||0)-(a.6a||0)}G(!b.35&&((b.47||b.47===0)?b.47:b.5a))b.35=b.47||b.5a;G(!b.59&&b.5E)b.59=b.5E;G(!b.35&&b.2s)b.35=(b.2s&1?1:(b.2s&2?3:(b.2s&4?2:0)));I b},3M:H(a,b){b.24=a.24=a.24||b.24||7.24++;I b},2t:{27:{4p:H(){55();I},4A:H(){I}},3D:{4p:H(){G(D.14.1f)I Q;D(7).2O("53",D.W.2t.3D.2y);I M},4A:H(){G(D.14.1f)I Q;D(7).4e("53",D.W.2t.3D.2y);I M},2y:H(a){G(F(a,7))I M;a.O="3D";I D.W.1H.1w(7,19)}},3N:{4p:H(){G(D.14.1f)I Q;D(7).2O("51",D.W.2t.3N.2y);I M},4A:H(){G(D.14.1f)I Q;D(7).4e("51",D.W.2t.3N.2y);I M},2y:H(a){G(F(a,7))I M;a.O="3N";I D.W.1H.1w(7,19)}}}};D.17.1l({2O:H(c,a,b){I c=="4X"?7.2V(c,a,b):7.P(H(){D.W.1e(7,c,b||a,b&&a)})},2V:H(d,b,c){J e=D.W.3M(c||b,H(a){D(7).4e(a,e);I(c||b).1w(7,19)});I 7.P(H(){D.W.1e(7,d,e,c&&b)})},4e:H(a,b){I 7.P(H(){D.W.21(7,a,b)})},1P:H(c,a,b){I 7.P(H(){D.W.1P(c,a,7,M,b)})},5C:H(c,a,b){I 7[0]&&D.W.1P(c,a,7[0],Q,b)},2m:H(b){J c=19,i=1;1B(i<c.K)D.W.3M(b,c[i++]);I 7.4V(D.W.3M(b,H(a){7.4Z=(7.4Z||0)%i;a.32();I c[7.4Z++].1w(7,19)||Q}))},7X:H(a,b){I 7.2O(\'3D\',a).2O(\'3N\',b)},27:H(a){55();G(D.2Q)a.1k(S,D);N D.3A.1p(H(){I a.1k(7,D)});I 7}});D.1l({2Q:Q,3A:[],27:H(){G(!D.2Q){D.2Q=M;G(D.3A){D.P(D.3A,H(){7.1k(S)});D.3A=U}D(S).5C("27")}}});J x=Q;H 55(){G(x)I;x=M;G(S.3K&&!D.14.2G)S.3K("69",D.27,Q);G(D.14.1f&&1b==1S)(H(){G(D.2Q)I;1U{S.1C.7V("1A")}1V(3e){3B(19.3L,0);I}D.27()})();G(D.14.2G)S.3K("69",H(){G(D.2Q)I;R(J i=0;i<S.4W.K;i++)G(S.4W[i].3R){3B(19.3L,0);I}D.27()},Q);G(D.14.2k){J a;(H(){G(D.2Q)I;G(S.3f!="68"&&S.3f!="1J"){3B(19.3L,0);I}G(a===12)a=D("V, 7A[7U=7S]").K;G(S.4W.K!=a){3B(19.3L,0);I}D.27()})()}D.W.1e(1b,"43",D.27)}D.P(("7R,7Q,43,85,4y,4X,4V,7P,"+"7O,7N,89,53,51,7M,2A,"+"5o,7L,7K,8d,3e").1R(","),H(i,b){D.17[b]=H(a){I a?7.2O(b,a):7.1P(b)}});J F=H(a,c){J b=a.4k;1B(b&&b!=c)1U{b=b.1d}1V(3e){b=c}I b==c};D(1b).2O("4X",H(){D("*").1e(S).4e()});D.17.1l({67:D.17.43,43:H(g,d,c){G(1j g!=\'23\')I 7.67(g);J e=g.1h(" ");G(e>=0){J i=g.3s(e,g.K);g=g.3s(0,e)}c=c||H(){};J f="2P";G(d)G(D.1D(d)){c=d;d=U}N{d=D.3n(d);f="6g"}J h=7;D.3Y({1a:g,O:f,1O:"2K",L:d,1J:H(a,b){G(b=="1W"||b=="7J")h.2K(i?D("<1v/>").3v(a.4U.1o(/<1m(.|\\s)*?\\/1m>/g,"")).2q(i):a.4U);h.P(c,[a.4U,b,a])}});I 7},aL:H(){I D.3n(7.7I())},7I:H(){I 7.2l(H(){I D.Y(7,"3V")?D.2d(7.aH):7}).1E(H(){I 7.34&&!7.3R&&(7.4J||/2A|6y/i.11(7.Y)||/1r|1G|3Q/i.11(7.O))}).2l(H(i,c){J b=D(7).6e();I b==U?U:b.1q==2p?D.2l(b,H(a,i){I{34:c.34,2x:a}}):{34:c.34,2x:b}}).3p()}});D.P("7H,7G,7F,7D,7C,7B".1R(","),H(i,o){D.17[o]=H(f){I 7.2O(o,f)}});J B=1z();D.1l({3p:H(d,b,a,c){G(D.1D(b)){a=b;b=U}I D.3Y({O:"2P",1a:d,L:b,1W:a,1O:c})},aE:H(b,a){I D.3p(b,U,a,"1m")},aD:H(c,b,a){I D.3p(c,b,a,"3z")},aC:H(d,b,a,c){G(D.1D(b)){a=b;b={}}I D.3Y({O:"6g",1a:d,L:b,1W:a,1O:c})},aA:H(a){D.1l(D.60,a)},60:{1a:5Z.5Q,26:M,O:"2P",2T:0,7z:"4R/x-ax-3V-aw",7x:M,31:M,L:U,5Y:U,3Q:U,4Q:{2N:"4R/2N, 1r/2N",2K:"1r/2K",1m:"1r/4t, 4R/4t",3z:"4R/3z, 1r/4t",1r:"1r/as",4w:"*/*"}},4z:{},3Y:H(s){s=D.1l(M,s,D.1l(M,{},D.60,s));J g,2Z=/=\\?(&|$)/g,1u,L,O=s.O.2r();G(s.L&&s.7x&&1j s.L!="23")s.L=D.3n(s.L);G(s.1O=="4P"){G(O=="2P"){G(!s.1a.1I(2Z))s.1a+=(s.1a.1I(/\\?/)?"&":"?")+(s.4P||"7u")+"=?"}N G(!s.L||!s.L.1I(2Z))s.L=(s.L?s.L+"&":"")+(s.4P||"7u")+"=?";s.1O="3z"}G(s.1O=="3z"&&(s.L&&s.L.1I(2Z)||s.1a.1I(2Z))){g="4P"+B++;G(s.L)s.L=(s.L+"").1o(2Z,"="+g+"$1");s.1a=s.1a.1o(2Z,"="+g+"$1");s.1O="1m";1b[g]=H(a){L=a;1W();1J();1b[g]=12;1U{2U 1b[g]}1V(e){}G(i)i.37(h)}}G(s.1O=="1m"&&s.1Y==U)s.1Y=Q;G(s.1Y===Q&&O=="2P"){J j=1z();J k=s.1a.1o(/(\\?|&)3m=.*?(&|$)/,"$ap="+j+"$2");s.1a=k+((k==s.1a)?(s.1a.1I(/\\?/)?"&":"?")+"3m="+j:"")}G(s.L&&O=="2P"){s.1a+=(s.1a.1I(/\\?/)?"&":"?")+s.L;s.L=U}G(s.26&&!D.4O++)D.W.1P("7H");J n=/^(?:\\w+:)?\\/\\/([^\\/?#]+)/;G(s.1O=="1m"&&O=="2P"&&n.11(s.1a)&&n.2D(s.1a)[1]!=5Z.al){J i=S.3H("6w")[0];J h=S.3h("1m");h.4d=s.1a;G(s.7t)h.aj=s.7t;G(!g){J l=Q;h.ah=h.ag=H(){G(!l&&(!7.3f||7.3f=="68"||7.3f=="1J")){l=M;1W();1J();i.37(h)}}}i.3U(h);I 12}J m=Q;J c=1b.7s?2B 7s("ae.ac"):2B 7r();G(s.5Y)c.6R(O,s.1a,s.31,s.5Y,s.3Q);N c.6R(O,s.1a,s.31);1U{G(s.L)c.4B("ab-aa",s.7z);G(s.5S)c.4B("a9-5R-a8",D.4z[s.1a]||"a7, a6 a5 a4 5N:5N:5N a2");c.4B("X-9Z-9Y","7r");c.4B("9W",s.1O&&s.4Q[s.1O]?s.4Q[s.1O]+", */*":s.4Q.4w)}1V(e){}G(s.7m&&s.7m(c,s)===Q){s.26&&D.4O--;c.7l();I Q}G(s.26)D.W.1P("7B",[c,s]);J d=H(a){G(!m&&c&&(c.3f==4||a=="2T")){m=M;G(f){7k(f);f=U}1u=a=="2T"&&"2T"||!D.7j(c)&&"3e"||s.5S&&D.7h(c,s.1a)&&"7J"||"1W";G(1u=="1W"){1U{L=D.6X(c,s.1O,s.9S)}1V(e){1u="5J"}}G(1u=="1W"){J b;1U{b=c.5I("7g-5R")}1V(e){}G(s.5S&&b)D.4z[s.1a]=b;G(!g)1W()}N D.5H(s,c,1u);1J();G(s.31)c=U}};G(s.31){J f=4I(d,13);G(s.2T>0)3B(H(){G(c){c.7l();G(!m)d("2T")}},s.2T)}1U{c.9P(s.L)}1V(e){D.5H(s,c,U,e)}G(!s.31)d();H 1W(){G(s.1W)s.1W(L,1u);G(s.26)D.W.1P("7C",[c,s])}H 1J(){G(s.1J)s.1J(c,1u);G(s.26)D.W.1P("7F",[c,s]);G(s.26&&!--D.4O)D.W.1P("7G")}I c},5H:H(s,a,b,e){G(s.3e)s.3e(a,b,e);G(s.26)D.W.1P("7D",[a,s,e])},4O:0,7j:H(a){1U{I!a.1u&&5Z.9O=="5p:"||(a.1u>=7e&&a.1u<9N)||a.1u==7c||a.1u==9K||D.14.2k&&a.1u==12}1V(e){}I Q},7h:H(a,c){1U{J b=a.5I("7g-5R");I a.1u==7c||b==D.4z[c]||D.14.2k&&a.1u==12}1V(e){}I Q},6X:H(a,c,b){J d=a.5I("9J-O"),2N=c=="2N"||!c&&d&&d.1h("2N")>=0,L=2N?a.9I:a.4U;G(2N&&L.1C.2j=="5J")7p"5J";G(b)L=b(L,c);G(c=="1m")D.5u(L);G(c=="3z")L=6u("("+L+")");I L},3n:H(a){J s=[];G(a.1q==2p||a.5w)D.P(a,H(){s.1p(3u(7.34)+"="+3u(7.2x))});N R(J j 1n a)G(a[j]&&a[j].1q==2p)D.P(a[j],H(){s.1p(3u(j)+"="+3u(7))});N s.1p(3u(j)+"="+3u(D.1D(a[j])?a[j]():a[j]));I s.6s("&").1o(/%20/g,"+")}});D.17.1l({1N:H(c,b){I c?7.2g({1Z:"1N",2h:"1N",1y:"1N"},c,b):7.1E(":1G").P(H(){7.V.18=7.5D||"";G(D.1g(7,"18")=="2F"){J a=D("<"+7.2j+" />").6P("1c");7.V.18=a.1g("18");G(7.V.18=="2F")7.V.18="3I";a.21()}}).3l()},1M:H(b,a){I b?7.2g({1Z:"1M",2h:"1M",1y:"1M"},b,a):7.1E(":4j").P(H(){7.5D=7.5D||D.1g(7,"18");7.V.18="2F"}).3l()},78:D.17.2m,2m:H(a,b){I D.1D(a)&&D.1D(b)?7.78.1w(7,19):a?7.2g({1Z:"2m",2h:"2m",1y:"2m"},a,b):7.P(H(){D(7)[D(7).3F(":1G")?"1N":"1M"]()})},9G:H(b,a){I 7.2g({1Z:"1N"},b,a)},9F:H(b,a){I 7.2g({1Z:"1M"},b,a)},9E:H(b,a){I 7.2g({1Z:"2m"},b,a)},9D:H(b,a){I 7.2g({1y:"1N"},b,a)},9M:H(b,a){I 7.2g({1y:"1M"},b,a)},9C:H(c,a,b){I 7.2g({1y:a},c,b)},2g:H(k,j,i,g){J h=D.77(j,i,g);I 7[h.36===Q?"P":"36"](H(){G(7.16!=1)I Q;J f=D.1l({},h),p,1G=D(7).3F(":1G"),46=7;R(p 1n k){G(k[p]=="1M"&&1G||k[p]=="1N"&&!1G)I f.1J.1k(7);G(p=="1Z"||p=="2h"){f.18=D.1g(7,"18");f.33=7.V.33}}G(f.33!=U)7.V.33="1G";f.45=D.1l({},k);D.P(k,H(c,a){J e=2B D.28(46,f,c);G(/2m|1N|1M/.11(a))e[a=="2m"?1G?"1N":"1M":a](k);N{J b=a.6r().1I(/^([+-]=)?([\\d+-.]+)(.*)$/),2b=e.1t(M)||0;G(b){J d=3d(b[2]),2M=b[3]||"2X";G(2M!="2X"){46.V[c]=(d||1)+2M;2b=((d||1)/e.1t(M))*2b;46.V[c]=2b+2M}G(b[1])d=((b[1]=="-="?-1:1)*d)+2b;e.3G(2b,d,2M)}N e.3G(2b,a,"")}});I M})},36:H(a,b){G(D.1D(a)||(a&&a.1q==2p)){b=a;a="28"}G(!a||(1j a=="23"&&!b))I A(7[0],a);I 7.P(H(){G(b.1q==2p)A(7,a,b);N{A(7,a).1p(b);G(A(7,a).K==1)b.1k(7)}})},9X:H(b,c){J a=D.3O;G(b)7.36([]);7.P(H(){R(J i=a.K-1;i>=0;i--)G(a[i].T==7){G(c)a[i](M);a.7n(i,1)}});G(!c)7.5A();I 7}});J A=H(b,c,a){G(b){c=c||"28";J q=D.L(b,c+"36");G(!q||a)q=D.L(b,c+"36",D.2d(a))}I q};D.17.5A=H(a){a=a||"28";I 7.P(H(){J q=A(7,a);q.4s();G(q.K)q[0].1k(7)})};D.1l({77:H(b,a,c){J d=b&&b.1q==a0?b:{1J:c||!c&&a||D.1D(b)&&b,2u:b,41:c&&a||a&&a.1q!=9t&&a};d.2u=(d.2u&&d.2u.1q==4L?d.2u:D.28.5K[d.2u])||D.28.5K.74;d.5M=d.1J;d.1J=H(){G(d.36!==Q)D(7).5A();G(D.1D(d.5M))d.5M.1k(7)};I d},41:{73:H(p,n,b,a){I b+a*p},5P:H(p,n,b,a){I((-29.9r(p*29.9q)/2)+0.5)*a+b}},3O:[],48:U,28:H(b,c,a){7.15=c;7.T=b;7.1i=a;G(!c.3Z)c.3Z={}}});D.28.44={4D:H(){G(7.15.2Y)7.15.2Y.1k(7.T,7.1z,7);(D.28.2Y[7.1i]||D.28.2Y.4w)(7);G(7.1i=="1Z"||7.1i=="2h")7.T.V.18="3I"},1t:H(a){G(7.T[7.1i]!=U&&7.T.V[7.1i]==U)I 7.T[7.1i];J r=3d(D.1g(7.T,7.1i,a));I r&&r>-9p?r:3d(D.2a(7.T,7.1i))||0},3G:H(c,b,d){7.5V=1z();7.2b=c;7.3l=b;7.2M=d||7.2M||"2X";7.1z=7.2b;7.2S=7.4N=0;7.4D();J e=7;H t(a){I e.2Y(a)}t.T=7.T;D.3O.1p(t);G(D.48==U){D.48=4I(H(){J a=D.3O;R(J i=0;i<a.K;i++)G(!a[i]())a.7n(i--,1);G(!a.K){7k(D.48);D.48=U}},13)}},1N:H(){7.15.3Z[7.1i]=D.1K(7.T.V,7.1i);7.15.1N=M;7.3G(0,7.1t());G(7.1i=="2h"||7.1i=="1Z")7.T.V[7.1i]="9m";D(7.T).1N()},1M:H(){7.15.3Z[7.1i]=D.1K(7.T.V,7.1i);7.15.1M=M;7.3G(7.1t(),0)},2Y:H(a){J t=1z();G(a||t>7.15.2u+7.5V){7.1z=7.3l;7.2S=7.4N=1;7.4D();7.15.45[7.1i]=M;J b=M;R(J i 1n 7.15.45)G(7.15.45[i]!==M)b=Q;G(b){G(7.15.18!=U){7.T.V.33=7.15.33;7.T.V.18=7.15.18;G(D.1g(7.T,"18")=="2F")7.T.V.18="3I"}G(7.15.1M)7.T.V.18="2F";G(7.15.1M||7.15.1N)R(J p 1n 7.15.45)D.1K(7.T.V,p,7.15.3Z[p])}G(b)7.15.1J.1k(7.T);I Q}N{J n=t-7.5V;7.4N=n/7.15.2u;7.2S=D.41[7.15.41||(D.41.5P?"5P":"73")](7.4N,n,0,1,7.15.2u);7.1z=7.2b+((7.3l-7.2b)*7.2S);7.4D()}I M}};D.1l(D.28,{5K:{9l:9j,9i:7e,74:9g},2Y:{2e:H(a){a.T.2e=a.1z},2c:H(a){a.T.2c=a.1z},1y:H(a){D.1K(a.T.V,"1y",a.1z)},4w:H(a){a.T.V[a.1i]=a.1z+a.2M}}});D.17.2i=H(){J b=0,1S=0,T=7[0],3q;G(T)ao(D.14){J d=T.1d,4a=T,1s=T.1s,1Q=T.2z,5U=2k&&3r(5B)<9c&&!/9a/i.11(v),1g=D.2a,3c=1g(T,"30")=="3c";G(T.7y){J c=T.7y();1e(c.1A+29.2f(1Q.1C.2e,1Q.1c.2e),c.1S+29.2f(1Q.1C.2c,1Q.1c.2c));1e(-1Q.1C.6b,-1Q.1C.6a)}N{1e(T.5X,T.5W);1B(1s){1e(1s.5X,1s.5W);G(42&&!/^t(98|d|h)$/i.11(1s.2j)||2k&&!5U)2C(1s);G(!3c&&1g(1s,"30")=="3c")3c=M;4a=/^1c$/i.11(1s.2j)?4a:1s;1s=1s.1s}1B(d&&d.2j&&!/^1c|2K$/i.11(d.2j)){G(!/^96|1T.*$/i.11(1g(d,"18")))1e(-d.2e,-d.2c);G(42&&1g(d,"33")!="4j")2C(d);d=d.1d}G((5U&&(3c||1g(4a,"30")=="5x"))||(42&&1g(4a,"30")!="5x"))1e(-1Q.1c.5X,-1Q.1c.5W);G(3c)1e(29.2f(1Q.1C.2e,1Q.1c.2e),29.2f(1Q.1C.2c,1Q.1c.2c))}3q={1S:1S,1A:b}}H 2C(a){1e(D.2a(a,"6V",M),D.2a(a,"6U",M))}H 1e(l,t){b+=3r(l,10)||0;1S+=3r(t,10)||0}I 3q};D.17.1l({30:H(){J a=0,1S=0,3q;G(7[0]){J b=7.1s(),2i=7.2i(),4c=/^1c|2K$/i.11(b[0].2j)?{1S:0,1A:0}:b.2i();2i.1S-=25(7,\'94\');2i.1A-=25(7,\'aF\');4c.1S+=25(b,\'6U\');4c.1A+=25(b,\'6V\');3q={1S:2i.1S-4c.1S,1A:2i.1A-4c.1A}}I 3q},1s:H(){J a=7[0].1s;1B(a&&(!/^1c|2K$/i.11(a.2j)&&D.1g(a,\'30\')==\'93\'))a=a.1s;I D(a)}});D.P([\'5e\',\'5G\'],H(i,b){J c=\'4y\'+b;D.17[c]=H(a){G(!7[0])I;I a!=12?7.P(H(){7==1b||7==S?1b.92(!i?a:D(1b).2e(),i?a:D(1b).2c()):7[c]=a}):7[0]==1b||7[0]==S?46[i?\'aI\':\'aJ\']||D.71&&S.1C[c]||S.1c[c]:7[0][c]}});D.P(["6N","4b"],H(i,b){J c=i?"5e":"5G",4f=i?"6k":"6i";D.17["5s"+b]=H(){I 7[b.3y()]()+25(7,"57"+c)+25(7,"57"+4f)};D.17["90"+b]=H(a){I 7["5s"+b]()+25(7,"2C"+c+"4b")+25(7,"2C"+4f+"4b")+(a?25(7,"6S"+c)+25(7,"6S"+4f):0)}})})();',62,669,'|||||||this|||||||||||||||||||||||||||||||||||if|function|return|var|length|data|true|else|type|each|false|for|document|elem|null|style|event||nodeName|||test|undefined||browser|options|nodeType|fn|display|arguments|url|window|body|parentNode|add|msie|css|indexOf|prop|typeof|call|extend|script|in|replace|push|constructor|text|offsetParent|cur|status|div|apply|firstChild|opacity|now|left|while|documentElement|isFunction|filter|className|hidden|handle|match|complete|attr|ret|hide|show|dataType|trigger|doc|split|top|table|try|catch|success|break|cache|height||remove|tbody|string|guid|num|global|ready|fx|Math|curCSS|start|scrollTop|makeArray|scrollLeft|max|animate|width|offset|tagName|safari|map|toggle||done|Array|find|toUpperCase|button|special|duration|id|copy|value|handler|ownerDocument|select|new|border|exec|stack|none|opera|nextSibling|pushStack|target|html|inArray|unit|xml|bind|GET|isReady|merge|pos|timeout|delete|one|selected|px|step|jsre|position|async|preventDefault|overflow|name|which|queue|removeChild|namespace|insertBefore|nth|removeData|fixed|parseFloat|error|readyState|multiFilter|createElement|rl|re|trim|end|_|param|first|get|results|parseInt|slice|childNodes|encodeURIComponent|append|events|elems|toLowerCase|json|readyList|setTimeout|grep|mouseenter|color|is|custom|getElementsByTagName|block|stopPropagation|addEventListener|callee|proxy|mouseleave|timers|defaultView|password|disabled|last|has|appendChild|form|domManip|props|ajax|orig|set|easing|mozilla|load|prototype|curAnim|self|charCode|timerId|object|offsetChild|Width|parentOffset|src|unbind|br|currentStyle|clean|float|visible|relatedTarget|previousSibling|handlers|isXMLDoc|on|setup|nodeIndex|unique|shift|javascript|child|RegExp|_default|deep|scroll|lastModified|teardown|setRequestHeader|timeStamp|update|empty|tr|getAttribute|innerHTML|setInterval|checked|fromElement|Number|jQuery|state|active|jsonp|accepts|application|dir|input|responseText|click|styleSheets|unload|not|lastToggle|outline|mouseout|getPropertyValue|mouseover|getComputedStyle|bindReady|String|padding|pageX|metaKey|keyCode|getWH|andSelf|clientX|Left|all|visibility|container|index|init|triggered|removeAttribute|classFilter|prevObject|submit|file|after|windowData|inner|client|globalEval|sibling|jquery|absolute|clone|wrapAll|dequeue|version|triggerHandler|oldblock|ctrlKey|createTextNode|Top|handleError|getResponseHeader|parsererror|speeds|checkbox|old|00|radio|swing|href|Modified|ifModified|lastChild|safari2|startTime|offsetTop|offsetLeft|username|location|ajaxSettings|getElementById|isSimple|values|selectedIndex|runtimeStyle|rsLeft|_load|loaded|DOMContentLoaded|clientTop|clientLeft|toElement|srcElement|val|pageY|POST|unshift|Bottom|clientY|Right|fix|exclusive|detachEvent|cloneNode|removeEventListener|swap|toString|join|attachEvent|eval|substr|head|parse|textarea|reset|image|zoom|odd|even|before|prepend|exclude|expr|quickClass|quickID|uuid|quickChild|continue|Height|textContent|appendTo|contents|open|margin|evalScript|borderTopWidth|borderLeftWidth|parent|httpData|setArray|CSS1Compat|compatMode|boxModel|cssFloat|linear|def|webkit|nodeValue|speed|_toggle|eq|100|replaceWith|304|concat|200|alpha|Last|httpNotModified|getAttributeNode|httpSuccess|clearInterval|abort|beforeSend|splice|styleFloat|throw|colgroup|XMLHttpRequest|ActiveXObject|scriptCharset|callback|fieldset|multiple|processData|getBoundingClientRect|contentType|link|ajaxSend|ajaxSuccess|ajaxError|col|ajaxComplete|ajaxStop|ajaxStart|serializeArray|notmodified|keypress|keydown|change|mouseup|mousedown|dblclick|focus|blur|stylesheet|hasClass|rel|doScroll|black|hover|solid|cancelBubble|returnValue|wheelDelta|view|round|shiftKey|resize|screenY|screenX|relatedNode|mousemove|prevValue|originalTarget|offsetHeight|keyup|newValue|offsetWidth|eventPhase|detail|currentTarget|cancelable|bubbles|attrName|attrChange|altKey|originalEvent|charAt|0n|substring|animated|header|noConflict|line|enabled|innerText|contains|only|weight|font|gt|lt|uFFFF|u0128|size|417|Boolean|Date|toggleClass|removeClass|addClass|removeAttr|replaceAll|insertAfter|prependTo|wrap|contentWindow|contentDocument|iframe|children|siblings|prevAll|wrapInner|nextAll|outer|prev|scrollTo|static|marginTop|next|inline|parents|able|cellSpacing|adobeair|cellspacing|522|maxLength|maxlength|readOnly|400|readonly|fast|600|class|slow|1px|htmlFor|reverse|10000|PI|cos|compatible|Function|setData|ie|ra|it|rv|getData|userAgent|navigator|fadeTo|fadeIn|slideToggle|slideUp|slideDown|ig|responseXML|content|1223|NaN|fadeOut|300|protocol|send|setAttribute|option|dataFilter|cssText|changed|be|Accept|stop|With|Requested|Object|can|GMT|property|1970|Jan|01|Thu|Since|If|Type|Content|XMLHTTP|th|Microsoft|td|onreadystatechange|onload|cap|charset|colg|host|tfoot|specified|with|1_|thead|leg|plain|attributes|opt|embed|urlencoded|www|area|hr|ajaxSetup|meta|post|getJSON|getScript|marginLeft|img|elements|pageYOffset|pageXOffset|abbr|serialize|pixelLeft'.split('|'),0,{}))
```
