## Appendix 3 for "Promoting long-term forest landscape resilience in the Lake Tahoe basin": jquery.tablesorter.min.js

```
(function($){$.extend({tablesorter:new function(){var parsers=[],widgets=[];this.defaults={cssHeader:"header",cssAsc:"headerSortUp",cssDesc:"headerSortDown",sortInitialOrder:"asc",sortMultiSortKey:"shiftKey",sortForce:null,sortAppend:null,textExtraction:"simple",parsers:{},widgets:[],widgetZebra:{css:["even","odd"]},headers:{},widthFixed:false,cancelSelection:true,sortList:[],headerList:[],dateFormat:"us",decimal:'.',debug:false};function benchmark(s,d){log(s+","+(new Date().getTime()-d.getTime())+"ms");}this.benchmark=benchmark;function log(s){if(typeof console!="undefined"&&typeof console.debug!="undefined"){console.log(s);}else{alert(s);}}function buildParserCache(table,$headers){if(table.config.debug){var parsersDebug="";}var rows=table.tBodies[0].rows;if(table.tBodies[0].rows[0]){var list=[],cells=rows[0].cells,l=cells.length;for(var i=0;i<l;i++){var p=false;if($.metadata&&($($headers[i]).metadata()&&$($headers[i]).metadata().sorter)){p=getParserById($($headers[i]).metadata().sorter);}else if((table.config.headers[i]&&table.config.headers[i].sorter)){p=getParserById(table.config.headers[i].sorter);}if(!p){p=detectParserForColumn(table,cells[i]);}if(table.config.debug){parsersDebug+="column:"+i+" parser:"+p.id+"\n";}list.push(p);}}if(table.config.debug){log(parsersDebug);}return list;};function detectParserForColumn(table,node){var l=parsers.length;for(var i=1;i<l;i++){if(parsers[i].is($.trim(getElementText(table.config,node)),table,node)){return parsers[i];}}return parsers[0];}function getParserById(name){var l=parsers.length;for(var i=0;i<l;i++){if(parsers[i].id.toLowerCase()==name.toLowerCase()){return parsers[i];}}return false;}function buildCache(table){if(table.config.debug){var cacheTime=new Date();}var totalRows=(table.tBodies[0]&&table.tBodies[0].rows.length)||0,totalCells=(table.tBodies[0].rows[0]&&table.tBodies[0].rows[0].cells.length)||0,parsers=table.config.parsers,cache={row:[],normalized:[]};for(var i=0;i<totalRows;++i){var c=table.tBodies[0].rows[i],cols=[];cache.row.push($(c));for(var j=0;j<totalCells;++j){cols.push(parsers[j].format(getElementText(table.config,c.cells[j]),table,c.cells[j]));}cols.push(i);cache.normalized.push(cols);cols=null;};if(table.config.debug){benchmark("Building cache for "+totalRows+" rows:",cacheTime);}return cache;};function getElementText(config,node){if(!node)return"";var t="";if(config.textExtraction=="simple"){if(node.childNodes[0]&&node.childNodes[0].hasChildNodes()){t=node.childNodes[0].innerHTML;}else{t=node.innerHTML;}}else{if(typeof(config.textExtraction)=="function"){t=config.textExtraction(node);}else{t=$(node).text();}}return t;}function appendToTable(table,cache){if(table.config.debug){var appendTime=new Date()}var c=cache,r=c.row,n=c.normalized,totalRows=n.length,checkCell=(n[0].length-1),tableBody=$(table.tBodies[0]),rows=[];for(var i=0;i<totalRows;i++){rows.push(r[n[i][checkCell]]);if(!table.config.appender){var o=r[n[i][checkCell]];var l=o.length;for(var j=0;j<l;j++){tableBody[0].appendChild(o[j]);}}}if(table.config.appender){table.config.appender(table,rows);}rows=null;if(table.config.debug){benchmark("Rebuilt table:",appendTime);}applyWidget(table);setTimeout(function(){$(table).trigger("sortEnd");},0);};function buildHeaders(table){if(table.config.debug){var time=new Date();}var meta=($.metadata)?true:false,tableHeadersRows=[];for(var i=0;i<table.tHead.rows.length;i++){tableHeadersRows[i]=0;};$tableHeaders=$("thead th",table);$tableHeaders.each(function(index){this.count=0;this.column=index;this.order=formatSortingOrder(table.config.sortInitialOrder);if(checkHeaderMetadata(this)||checkHeaderOptions(table,index))this.sortDisabled=true;if(!this.sortDisabled){$(this).addClass(table.config.cssHeader);}table.config.headerList[index]=this;});if(table.config.debug){benchmark("Built headers:",time);log($tableHeaders);}return $tableHeaders;};function checkCellColSpan(table,rows,row){var arr=[],r=table.tHead.rows,c=r[row].cells;for(var i=0;i<c.length;i++){var cell=c[i];if(cell.colSpan>1){arr=arr.concat(checkCellColSpan(table,headerArr,row++));}else{if(table.tHead.length==1||(cell.rowSpan>1||!r[row+1])){arr.push(cell);}}}return arr;};function checkHeaderMetadata(cell){if(($.metadata)&&($(cell).metadata().sorter===false)){return true;};return false;}function checkHeaderOptions(table,i){if((table.config.headers[i])&&(table.config.headers[i].sorter===false)){return true;};return false;}function applyWidget(table){var c=table.config.widgets;var l=c.length;for(var i=0;i<l;i++){getWidgetById(c[i]).format(table);}}function getWidgetById(name){var l=widgets.length;for(var i=0;i<l;i++){if(widgets[i].id.toLowerCase()==name.toLowerCase()){return widgets[i];}}};function formatSortingOrder(v){if(typeof(v)!="Number"){i=(v.toLowerCase()=="desc")?1:0;}else{i=(v==(0||1))?v:0;}return i;}function isValueInArray(v,a){var l=a.length;for(var i=0;i<l;i++){if(a[i][0]==v){return true;}}return false;}function setHeadersCss(table,$headers,list,css){$headers.removeClass(css[0]).removeClass(css[1]);var h=[];$headers.each(function(offset){if(!this.sortDisabled){h[this.column]=$(this);}});var l=list.length;for(var i=0;i<l;i++){h[list[i][0]].addClass(css[list[i][1]]);}}function fixColumnWidth(table,$headers){var c=table.config;if(c.widthFixed){var colgroup=$('<colgroup>');$("tr:first td",table.tBodies[0]).each(function(){colgroup.append($('<col>').css('width',$(this).width()));});$(table).prepend(colgroup);};}function updateHeaderSortCount(table,sortList){var c=table.config,l=sortList.length;for(var i=0;i<l;i++){var s=sortList[i],o=c.headerList[s[0]];o.count=s[1];o.count++;}}function multisort(table,sortList,cache){if(table.config.debug){var sortTime=new Date();}var dynamicExp="var sortWrapper = function(a,b) {",l=sortList.length;for(var i=0;i<l;i++){var c=sortList[i][0];var order=sortList[i][1];var s=(getCachedSortType(table.config.parsers,c)=="text")?((order==0)?"sortText":"sortTextDesc"):((order==0)?"sortNumeric":"sortNumericDesc");var e="e"+i;dynamicExp+="var "+e+" = "+s+"(a["+c+"],b["+c+"]); ";dynamicExp+="if("+e+") { return "+e+"; } ";dynamicExp+="else { ";}var orgOrderCol=cache.normalized[0].length-1;dynamicExp+="return a["+orgOrderCol+"]-b["+orgOrderCol+"];";for(var i=0;i<l;i++){dynamicExp+="}; ";}dynamicExp+="return 0; ";dynamicExp+="}; ";eval(dynamicExp);cache.normalized.sort(sortWrapper);if(table.config.debug){benchmark("Sorting on "+sortList.toString()+" and dir "+order+" time:",sortTime);}return cache;};function sortText(a,b){return((a<b)?-1:((a>b)?1:0));};function sortTextDesc(a,b){return((b<a)?-1:((b>a)?1:0));};function sortNumeric(a,b){return a-b;};function sortNumericDesc(a,b){return b-a;};function getCachedSortType(parsers,i){return parsers[i].type;};this.construct=function(settings){return this.each(function(){if(!this.tHead||!this.tBodies)return;var $this,$document,$headers,cache,config,shiftDown=0,sortOrder;this.config={};config=$.extend(this.config,$.tablesorter.defaults,settings);$this=$(this);$headers=buildHeaders(this);this.config.parsers=buildParserCache(this,$headers);cache=buildCache(this);var sortCSS=[config.cssDesc,config.cssAsc];fixColumnWidth(this);$headers.click(function(e){$this.trigger("sortStart");var totalRows=($this[0].tBodies[0]&&$this[0].tBodies[0].rows.length)||0;if(!this.sortDisabled&&totalRows>0){var $cell=$(this);var i=this.column;this.order=this.count++%2;if(!e[config.sortMultiSortKey]){config.sortList=[];if(config.sortForce!=null){var a=config.sortForce;for(var j=0;j<a.length;j++){if(a[j][0]!=i){config.sortList.push(a[j]);}}}config.sortList.push([i,this.order]);}else{if(isValueInArray(i,config.sortList)){for(var j=0;j<config.sortList.length;j++){var s=config.sortList[j],o=config.headerList[s[0]];if(s[0]==i){o.count=s[1];o.count++;s[1]=o.count%2;}}}else{config.sortList.push([i,this.order]);}};setTimeout(function(){setHeadersCss($this[0],$headers,config.sortList,sortCSS);appendToTable($this[0],multisort($this[0],config.sortList,cache));},1);return false;}}).mousedown(function(){if(config.cancelSelection){this.onselectstart=function(){return false};return false;}});$this.bind("update",function(){this.config.parsers=buildParserCache(this,$headers);cache=buildCache(this);}).bind("sorton",function(e,list){$(this).trigger("sortStart");config.sortList=list;var sortList=config.sortList;updateHeaderSortCount(this,sortList);setHeadersCss(this,$headers,sortList,sortCSS);appendToTable(this,multisort(this,sortList,cache));}).bind("appendCache",function(){appendToTable(this,cache);}).bind("applyWidgetId",function(e,id){getWidgetById(id).format(this);}).bind("applyWidgets",function(){applyWidget(this);});if($.metadata&&($(this).metadata()&&$(this).metadata().sortlist)){config.sortList=$(this).metadata().sortlist;}if(config.sortList.length>0){$this.trigger("sorton",[config.sortList]);}applyWidget(this);});};this.addParser=function(parser){var l=parsers.length,a=true;for(var i=0;i<l;i++){if(parsers[i].id.toLowerCase()==parser.id.toLowerCase()){a=false;}}if(a){parsers.push(parser);};};this.addWidget=function(widget){widgets.push(widget);};this.formatFloat=function(s){var i=parseFloat(s);return(isNaN(i))?0:i;};this.formatInt=function(s){var i=parseInt(s);return(isNaN(i))?0:i;};this.isDigit=function(s,config){var DECIMAL='\\'+config.decimal;var exp='/(^[+]?0('+DECIMAL+'0+)?$)|(^([-+]?[1-9][0-9]*)$)|(^([-+]?((0?|[1-9][0-9]*)'+DECIMAL+'(0*[1-9][0-9]*)))$)|(^[-+]?[1-9]+[0-9]*'+DECIMAL+'0+$)/';return RegExp(exp).test($.trim(s));};this.clearTableBody=function(table){if($.browser.msie){function empty(){while(this.firstChild)this.removeChild(this.firstChild);}empty.apply(table.tBodies[0]);}else{table.tBodies[0].innerHTML="";}};}});$.fn.extend({tablesorter:$.tablesorter.construct});var ts=$.tablesorter;ts.addParser({id:"text",is:function(s){return true;},format:function(s){return $.trim(s.toLowerCase());},type:"text"});ts.addParser({id:"digit",is:function(s,table){var c=table.config;return $.tablesorter.isDigit(s,c);},format:function(s){return $.tablesorter.formatFloat(s);},type:"numeric"});ts.addParser({id:"currency",is:function(s){return/^[£$€?.]/.test(s);},format:function(s){return $.tablesorter.formatFloat(s.replace(new RegExp(/[^0-9.]/g),""));},type:"numeric"});ts.addParser({id:"ipAddress",is:function(s){return/^\d{2,3}[\.]\d{2,3}[\.]\d{2,3}[\.]\d{2,3}$/.test(s);},format:function(s){var a=s.split("."),r="",l=a.length;for(var i=0;i<l;i++){var item=a[i];if(item.length==2){r+="0"+item;}else{r+=item;}}return $.tablesorter.formatFloat(r);},type:"numeric"});ts.addParser({id:"url",is:function(s){return/^(https?|ftp|file):\/\/$/.test(s);},format:function(s){return jQuery.trim(s.replace(new RegExp(/(https?|ftp|file):\/\//),''));},type:"text"});ts.addParser({id:"isoDate",is:function(s){return/^\d{4}[\/-]\d{1,2}[\/-]\d{1,2}$/.test(s);},format:function(s){return $.tablesorter.formatFloat((s!="")?new Date(s.replace(new RegExp(/-/g),"/")).getTime():"0");},type:"numeric"});ts.addParser({id:"percent",is:function(s){return/\%$/.test($.trim(s));},format:function(s){return $.tablesorter.formatFloat(s.replace(new RegExp(/%/g),""));},type:"numeric"});ts.addParser({id:"usLongDate",is:function(s){return s.match(new RegExp(/^[A-Za-z]{3,10}\.? [0-9]{1,2}, ([0-9]{4}|'?[0-9]{2}) (([0-2]?[0-9]:[0-5][0-9])|([0-1]?[0-9]:[0-5][0-9]\s(AM|PM)))$/));},format:function(s){return $.tablesorter.formatFloat(new Date(s).getTime());},type:"numeric"});ts.addParser({id:"shortDate",is:function(s){return/\d{1,2}[\/\-]\d{1,2}[\/\-]\d{2,4}/.test(s);},format:function(s,table){var c=table.config;s=s.replace(/\-/g,"/");if(c.dateFormat=="us"){s=s.replace(/(\d{1,2})[\/\-](\d{1,2})[\/\-](\d{4})/,"$3/$1/$2");}else if(c.dateFormat=="uk"){s=s.replace(/(\d{1,2})[\/\-](\d{1,2})[\/\-](\d{4})/,"$3/$2/$1");}else if(c.dateFormat=="dd/mm/yy"||c.dateFormat=="dd-mm-yy"){s=s.replace(/(\d{1,2})[\/\-](\d{1,2})[\/\-](\d{2})/,"$1/$2/$3");}return $.tablesorter.formatFloat(new Date(s).getTime());},type:"numeric"});ts.addParser({id:"time",is:function(s){return/^(([0-2]?[0-9]:[0-5][0-9])|([0-1]?[0-9]:[0-5][0-9]\s(am|pm)))$/.test(s);},format:function(s){return $.tablesorter.formatFloat(new Date("2000/01/01 "+s).getTime());},type:"numeric"});ts.addParser({id:"metadata",is:function(s){return false;},format:function(s,table,cell){var c=table.config,p=(!c.parserMetadataName)?'sortValue':c.parserMetadataName;return $(cell).metadata()[p];},type:"numeric"});ts.addWidget({id:"zebra",format:function(table){if(table.config.debug){var time=new Date();}$("tr:visible",table.tBodies[0]).filter(':even').removeClass(table.config.widgetZebra.css[1]).addClass(table.config.widgetZebra.css[0]).end().filter(':odd').removeClass(table.config.widgetZebra.css[0]).addClass(table.config.widgetZebra.css[1]);if(table.config.debug){$.tablesorter.benchmark("Applying Zebra widget",time);}}});})(jQuery);
```
