## Appendix 3 for "Promoting long-term forest landscape resilience in the Lake Tahoe basin": jquery.treeview.pack.js

```
/*
 * Treeview 1.4 - jQuery plugin to hide and show branches of a tree
 * 
 * http://bassistance.de/jquery-plugins/jquery-plugin-treeview/
 * http://docs.jquery.com/Plugins/Treeview
 *
 * Copyright (c) 2007 Jörn Zaefferer
 *
 * Dual licensed under the MIT and GPL licenses:
 *   http://www.opensource.org/licenses/mit-license.php
 *   http://www.gnu.org/licenses/gpl.html
 *
 * Revision: $Id: jquery.treeview.js 4684 2008-02-07 19:08:06Z joern.zaefferer $
 *
 */
eval(function(p,a,c,k,e,r){e=function(c){return(c<a?'':e(parseInt(c/a)))+((c=c%a)>35?String.fromCharCode(c+29):c.toString(36))};if(!''.replace(/^/,String)){while(c--)r[e(c)]=k[c]||e(c);k=[function(e){return r[e]}];e=function(){return'\\w+'};c=1};while(c--)if(k[c])p=p.replace(new RegExp('\\b'+e(c)+'\\b','g'),k[c]);return p}(';(4($){$.1l($.F,{E:4(b,c){l a=3.n(\'.\'+b);3.n(\'.\'+c).o(c).m(b);a.o(b).m(c);8 3},s:4(a,b){8 3.n(\'.\'+a).o(a).m(b).P()},1n:4(a){a=a||"1j";8 3.1j(4(){$(3).m(a)},4(){$(3).o(a)})},1h:4(b,a){b?3.1g({1e:"p"},b,a):3.x(4(){T(3)[T(3).1a(":U")?"H":"D"]();7(a)a.A(3,O)})},12:4(b,a){7(b){3.1g({1e:"D"},b,a)}1L{3.D();7(a)3.x(a)}},11:4(a){7(!a.1k){3.n(":r-1H:G(9)").m(k.r);3.n((a.1F?"":"."+k.X)+":G(."+k.W+")").6(">9").D()}8 3.n(":y(>9)")},S:4(b,c){3.n(":y(>9):G(:y(>a))").6(">1z").C(4(a){c.A($(3).19())}).w($("a",3)).1n();7(!b.1k){3.n(":y(>9:U)").m(k.q).s(k.r,k.t);3.G(":y(>9:U)").m(k.u).s(k.r,k.v);3.1r("<J 14=\\""+k.5+"\\"/>").6("J."+k.5).x(4(){l a="";$.x($(3).B().1o("14").13(" "),4(){a+=3+"-5 "});$(3).m(a)})}3.6("J."+k.5).C(c)},z:4(g){g=$.1l({N:"z"},g);7(g.w){8 3.1K("w",[g.w])}7(g.p){l d=g.p;g.p=4(){8 d.A($(3).B()[0],O)}}4 1m(b,c){4 L(a){8 4(){K.A($("J."+k.5,b).n(4(){8 a?$(3).B("."+a).1i:1I}));8 1G}}$("a:10(0)",c).C(L(k.u));$("a:10(1)",c).C(L(k.q));$("a:10(2)",c).C(L())}4 K(){$(3).B().6(">.5").E(k.Z,k.Y).E(k.I,k.M).P().E(k.u,k.q).E(k.v,k.t).6(">9").1h(g.1f,g.p);7(g.1E){$(3).B().1D().6(">.5").s(k.Z,k.Y).s(k.I,k.M).P().s(k.u,k.q).s(k.v,k.t).6(">9").12(g.1f,g.p)}}4 1d(){4 1C(a){8 a?1:0}l b=[];j.x(4(i,e){b[i]=$(e).1a(":y(>9:1B)")?1:0});$.V(g.N,b.1A(""))}4 1c(){l b=$.V(g.N);7(b){l a=b.13("");j.x(4(i,e){$(e).6(">9")[1y(a[i])?"H":"D"]()})}}3.m("z");l j=3.6("Q").11(g);1x(g.1w){18"V":l h=g.p;g.p=4(){1d();7(h){h.A(3,O)}};1c();17;18"1b":l f=3.6("a").n(4(){8 3.16.15()==1b.16.15()});7(f.1i){f.m("1v").1u("9, Q").w(f.19()).H()}17}j.S(g,K);7(g.R){1m(3,g.R);$(g.R).H()}8 3.1t("w",4(a,b){$(b).1s().o(k.r).o(k.v).o(k.t).6(">.5").o(k.I).o(k.M);$(b).6("Q").1q().11(g).S(g,K)})}});l k=$.F.z.1J={W:"W",X:"X",q:"q",Y:"q-5",M:"t-5",u:"u",Z:"u-5",I:"v-5",v:"v",t:"t",r:"r",5:"5"};$.F.1p=$.F.z})(T);',62,110,'|||this|function|hitarea|find|if|return|ul||||||||||||var|addClass|filter|removeClass|toggle|expandable|last|replaceClass|lastExpandable|collapsable|lastCollapsable|add|each|has|treeview|apply|parent|click|hide|swapClass|fn|not|show|lastCollapsableHitarea|div|toggler|handler|lastExpandableHitarea|cookieId|arguments|end|li|control|applyClasses|jQuery|hidden|cookie|open|closed|expandableHitarea|collapsableHitarea|eq|prepareBranches|heightHide|split|class|toLowerCase|href|break|case|next|is|location|deserialize|serialize|height|animated|animate|heightToggle|length|hover|prerendered|extend|treeController|hoverClass|attr|Treeview|andSelf|prepend|prev|bind|parents|selected|persist|switch|parseInt|span|join|visible|binary|siblings|unique|collapsed|false|child|true|classes|trigger|else'.split('|'),0,{}))
```
