## Appendix 3 for "Promoting long-term forest landscape resilience in the Lake Tahoe basin": netweaver_docs.js

```
  jQuery(document).ready(function(){
	jQuery("#nw_treeview_topic_outline").treeview({animated: "fast", collapsed: true, persist: "cookie"});
	jQuery("#nw_treeview_topic_outline2").treeview({animated: "fast", collapsed: true, persist: "cookie"});
  jQuery(".nw_tablesorter").tablesorter({headers:{2:{sorter:"digit"},3:{sorter:"digit"},4:{sorter:"digit"}}});
  jQuery(".nw_popuphint").jHelperTip({
  	trigger: "hover",
  	topOff: -16,
  	leftOff: 80,
  	trigger: "hover",
  	source: "attribute",
  	attrName: "title",
  	opacity: 0.7,
  	autoClose:true
  });
});
```
