## Appendix 3 for "Promoting long-term forest landscape resilience in the Lake Tahoe basin": Apex_predators.html


### Apex predators

#### dependency network

#### Topics that are influenced by Apex predators:

- Apex predators
  - Wildlife conservation
    - LTW resilience final

#### Topics that influence Apex predators:

- Apex predators
  - CASPO territories
    - CASPO1
    - CASPOI1
    - CASPO2
    - CASPOI2
    - CASPO
  - NOGO territories
    - NOGO1
    - NOGOI1
    - NOGO2
    - NOGOI2
    - NOGO
  - Marten territories
    - Marten1
    - MartenI1
    - Marten2
    - MartenI2
    - Marten

[ untitled0 ]
