## Appendix 3 for "Promoting long-term forest landscape resilience in the Lake Tahoe basin": Aspen.html


### Aspen

% of total biomass that is aspen

#### Topics that are influenced by Aspen:

- Aspen
  - Aspen forest
    - Composition
      - Upland vegetation health
        - LTW resilience final

[ untitled0 ]
