## Appendix 3 for "Promoting long-term forest landscape resilience in the Lake Tahoe basin": Aspen_forest.html


### Aspen forest

#### dependency network

#### Topics that are influenced by Aspen forest:

- Aspen forest
  - Composition
    - Upland vegetation health
      - LTW resilience final

#### Topics that influence Aspen forest:

- Aspen forest
  - Aspen1
  - AspenI1
  - Aspen2
  - AspenI2
  - Aspen3
  - AspenI3
  - Aspen4
  - AspenI4
  - Aspen

[ untitled0 ]
