## Appendix 3 for "Promoting long-term forest landscape resilience in the Lake Tahoe basin": Big_trees.html


### Big trees

#### dependency network

#### Topics that are influenced by Big trees:

- Big trees
  - Upland vegetation health
    - LTW resilience final

#### Topics that influence Big trees:

- Big trees
  - BigTrees1
  - BigTreesI1
  - BigTrees2
  - BigTreesI2
  - BigTrees3
  - BigTreesI3
  - BigTrees

[ untitled0 ]
