## Appendix 3 for "Promoting long-term forest landscape resilience in the Lake Tahoe basin": BigTrees.html


### BigTrees

### hectares with at least one tree >150 years old

#### BigTrees is compared in the following ways:

| x | y |
| --- | --- |

*(not in use)*  
  
*(not in use)*

#### Topics that are influenced by BigTrees:

- BigTrees
  - Big trees
    - Upland vegetation health
      - LTW resilience final

[ untitled0 ]
