## Appendix 3 for "Promoting long-term forest landscape resilience in the Lake Tahoe basin": Burned_low_sev.html


### Burned low sev

#### dependency network

#### Topics that are influenced by Burned low sev:

- Burned low sev
  - Percent land burned
    - Functional fire
      - LTW resilience final

#### Topics that influence Burned low sev:

- Burned low sev
  - PercentLandscapeLow1
  - PercentLandscapeLowI1
  - PercentLandscapeLow2
  - PercentLandscapeLowI2
  - PercentLandscapeLow3
  - PercentLandscapeLowI3
  - PercentLandscapeLow4
  - PercentLandscapeLowI4
  - PercentLandscapeLow5
  - PercentLandscapeLowI5
  - PercentLandscapeLow6
  - PercentLandscapeLowI6
  - PercentLandscapeLow

[ untitled0 ]
