## Appendix 3 for "Promoting long-term forest landscape resilience in the Lake Tahoe basin": Burned_mod_sev.html


### Burned mod sev

#### dependency network

#### Topics that are influenced by Burned mod sev:

- Burned mod sev
  - Percent land burned
    - Functional fire
      - LTW resilience final

#### Topics that influence Burned mod sev:

- Burned mod sev
  - PercentLandscapeMod1
  - PercentLandscapeModI1
  - PercentLandscapeMod2
  - PercentLandscapeModI2
  - PercentLandscapeMod3
  - PercentLandscapeModI3
  - PercentLandscapeMod4
  - PercentLandscapeModI4
  - PercentLandscapeMod5
  - PercentLandscapeModI5
  - PercentLandscapeMod

[ untitled0 ]
