## Appendix 3 for "Promoting long-term forest landscape resilience in the Lake Tahoe basin": CASPO.html


### CASPO

### territories/max. # territories supported on the landscape

#### Topics that are influenced by CASPO:

- CASPO
  - CASPO territories
    - Apex predators
      - Wildlife conservation
        - LTW resilience final

[ untitled0 ]
