## Appendix 3 for "Promoting long-term forest landscape resilience in the Lake Tahoe basin": CASPO_territories.html


### CASPO territories

#### dependency network

#### Topics that are influenced by CASPO territories:

- CASPO territories
  - Apex predators
    - Wildlife conservation
      - LTW resilience final

#### Topics that influence CASPO territories:

- CASPO territories
  - CASPO1
  - CASPOI1
  - CASPO2
  - CASPOI2
  - CASPO

[ untitled0 ]
