## Appendix 3 for "Promoting long-term forest landscape resilience in the Lake Tahoe basin": Composition.html


### Composition

#### dependency network

#### Topics that are influenced by Composition:

- Composition
  - Upland vegetation health
    - LTW resilience final

#### Topics that influence Composition:

- Composition
  - Yellow pine forest
    - YellowPine1
    - YellowPineI1
    - YellowPine2
    - YellowPineI2
    - YellowPine3
    - YellowPineI3
    - YellowPine4
    - YellowPineI4
    - YellowPine
  - White pine forest
    - WhitePine1
    - WhitePineI1
    - WhitePine2
    - WhitePineI2
    - WhitePine3
    - WhitePineI3
    - WhitePine4
    - WhitePineI4
    - WhitePine
  - Aspen forest
    - Aspen1
    - AspenI1
    - Aspen2
    - AspenI2
    - Aspen3
    - AspenI3
    - Aspen4
    - AspenI4
    - Aspen

[ untitled0 ]
