## Appendix 3 for "Promoting long-term forest landscape resilience in the Lake Tahoe basin": Conifer_cover.html


### Conifer cover

#### dependency network

#### Topics that are influenced by Conifer cover:

- Conifer cover
  - Forest cover
    - Upland vegetation health
      - LTW resilience final

#### Topics that influence Conifer cover:

- Conifer cover
  - Forest1
  - ForestI1
  - Forest2
  - ForestI2
  - Forest3
  - ForestI3
  - Forest4
  - ForestI4
  - Forest5
  - ForestI5
  - Forest

[ untitled0 ]
