## Appendix 3 for "Promoting long-term forest landscape resilience in the Lake Tahoe basin": Cult_water_quant_and_timing.html


### Cult water quant and timing

#### dependency network

#### Topics that are influenced by Cult water quant and timing:

- Cult water quant and timing
  - Cultural resource quality
    - LTW resilience final

#### Topics that influence Cult water quant and timing:

- Cult water quant and timing
  - LMWtrQuantTime1
  - LMWtrQuantTimeI1
  - LMWtrQuantTime2
  - LMWtrQuantTimeI2
  - LMWtrQuantTime

[ untitled0 ]
