## Appendix 3 for "Promoting long-term forest landscape resilience in the Lake Tahoe basin": CultASP4.html


### CultASP4

#### Topics that are influenced by CultASP4:

- CultASP4
  - HCKS aspen habitat
    - Habitat for cultural keystone species
      - Cultural resource quality
        - LTW resilience final

[ untitled0 ]
