## Appendix 3 for "Promoting long-term forest landscape resilience in the Lake Tahoe basin": CultASP.html


### CultASP

% of landscape dominated by aspen

#### Topics that are influenced by CultASP:

- CultASP
  - HCKS aspen habitat
    - Habitat for cultural keystone species
      - Cultural resource quality
        - LTW resilience final

[ untitled0 ]
