## Appendix 3 for "Promoting long-term forest landscape resilience in the Lake Tahoe basin": CultASPI1.html


### CultASPI1

#### Topics that are influenced by CultASPI1:

- CultASPI1
  - HCKS aspen habitat
    - Habitat for cultural keystone species
      - Cultural resource quality
        - LTW resilience final

[ untitled0 ]
