## Appendix 3 for "Promoting long-term forest landscape resilience in the Lake Tahoe basin": CultASPI2.html


### CultASPI2

#### Topics that are influenced by CultASPI2:

- CultASPI2
  - HCKS aspen habitat
    - Habitat for cultural keystone species
      - Cultural resource quality
        - LTW resilience final

[ untitled0 ]
