## Appendix 3 for "Promoting long-term forest landscape resilience in the Lake Tahoe basin": CultASPI4.html


### CultASPI4

#### Topics that are influenced by CultASPI4:

- CultASPI4
  - HCKS aspen habitat
    - Habitat for cultural keystone species
      - Cultural resource quality
        - LTW resilience final

[ untitled0 ]
