## Appendix 3 for "Promoting long-term forest landscape resilience in the Lake Tahoe basin": CultFire.html


### CultFire

% of landscape with low severity fire per decade

#### Topics that are influenced by CultFire:

- CultFire
  - Restoration of low intensity fire
    - Cultural resource quality
      - LTW resilience final

[ untitled0 ]
