## Appendix 3 for "Promoting long-term forest landscape resilience in the Lake Tahoe basin": Decomp_funct_redund.html


### Decomp funct redund

#### dependency network

#### Topics that are influenced by Decomp funct redund:

- Decomp funct redund
  - Ecological function
    - Wildlife conservation
      - LTW resilience final

#### Topics that influence Decomp funct redund:

- Decomp funct redund
  - Decomposer1
  - DecomposerI1
  - Decomposer2
  - DecomposerI2
  - Decomposer

[ untitled0 ]
