## Appendix 3 for "Promoting long-term forest landscape resilience in the Lake Tahoe basin": Decomposer.html


### Decomposer

### decomposers in which 70% of their 2010 habitat is maintained/# decomposers

#### Topics that are influenced by Decomposer:

- Decomposer
  - Decomp funct redund
    - Ecological function
      - Wildlife conservation
        - LTW resilience final

[ untitled0 ]
