## Appendix 3 for "Promoting long-term forest landscape resilience in the Lake Tahoe basin": DecomposerI1.html


### DecomposerI1

#### Topics that are influenced by DecomposerI1:

- DecomposerI1
  - Decomp funct redund
    - Ecological function
      - Wildlife conservation
        - LTW resilience final

[ untitled0 ]
