## Appendix 3 for "Promoting long-term forest landscape resilience in the Lake Tahoe basin": DecomposerI2.html


### DecomposerI2

#### Topics that are influenced by DecomposerI2:

- DecomposerI2
  - Decomp funct redund
    - Ecological function
      - Wildlife conservation
        - LTW resilience final

[ untitled0 ]
