## Appendix 3 for "Promoting long-term forest landscape resilience in the Lake Tahoe basin": Disperser.html


### Disperser

### spore disperser in which 70% of their 2010 habitat is maintained/# spore dispersers

#### Topics that are influenced by Disperser:

- Disperser
  - Seed disp funct redund
    - Ecological function
      - Wildlife conservation
        - LTW resilience final

[ untitled0 ]
