## Appendix 3 for "Promoting long-term forest landscape resilience in the Lake Tahoe basin": DisperserI1.html


### DisperserI1

#### Topics that are influenced by DisperserI1:

- DisperserI1
  - Seed disp funct redund
    - Ecological function
      - Wildlife conservation
        - LTW resilience final

[ untitled0 ]
