## Appendix 3 for "Promoting long-term forest landscape resilience in the Lake Tahoe basin": Early_seral_beta_div.html


### Early seral beta div

#### dependency network

#### Topics that are influenced by Early seral beta div:

- Early seral beta div
  - Species diversity
    - Wildlife conservation
      - LTW resilience final

#### Topics that influence Early seral beta div:

- Early seral beta div
  - EarlyBeta1
  - EarlyBetaI1
  - EarlyBeta2
  - EarlyBetaI2
  - EarlyBeta

[ untitled0 ]
