## Appendix 3 for "Promoting long-term forest landscape resilience in the Lake Tahoe basin": EarlyBeta.html


### EarlyBeta

### species that have habitat in early seral patches/# species

#### Topics that are influenced by EarlyBeta:

- EarlyBeta
  - Early seral beta div
    - Species diversity
      - Wildlife conservation
        - LTW resilience final

[ untitled0 ]
