## Appendix 3 for "Promoting long-term forest landscape resilience in the Lake Tahoe basin": ExtremeEmissionDays.html


### ExtremeEmissionDays

### Days >= 500 tons PM2.5 (Extreme)

#### Topics that are influenced by ExtremeEmissionDays:

- ExtremeEmissionDays
  - Qual air extreme emission days
    - Quality air
      - LTW resilience final

[ untitled0 ]
