## Appendix 3 for "Promoting long-term forest landscape resilience in the Lake Tahoe basin": Fine_sediment_load.html


### Fine sediment load

#### dependency network

#### Topics that are influenced by Fine sediment load:

- Fine sediment load
  - Quality water
    - LTW resilience final

#### Topics that influence Fine sediment load:

- Fine sediment load
  - Sediment1
  - SedimentI1
  - Sediment2
  - SedimentI2
  - Sediment

[ untitled0 ]
