## Appendix 3 for "Promoting long-term forest landscape resilience in the Lake Tahoe basin": Flicker1.html


### Flicker1

#### Topics that are influenced by Flicker1:

- Flicker1
  - HCKS flicker habitat
    - Habitat for cultural keystone species
      - Cultural resource quality
        - LTW resilience final

[ untitled0 ]
