## Appendix 3 for "Promoting long-term forest landscape resilience in the Lake Tahoe basin": Flicker2.html


### Flicker2

#### Topics that are influenced by Flicker2:

- Flicker2
  - HCKS flicker habitat
    - Habitat for cultural keystone species
      - Cultural resource quality
        - LTW resilience final

[ untitled0 ]
