## Appendix 3 for "Promoting long-term forest landscape resilience in the Lake Tahoe basin": Flicker.html


### Flicker

% of high quality habitat

#### Topics that are influenced by Flicker:

- Flicker
  - HCKS flicker habitat
    - Habitat for cultural keystone species
      - Cultural resource quality
        - LTW resilience final

[ untitled0 ]
