## Appendix 3 for "Promoting long-term forest landscape resilience in the Lake Tahoe basin": Forest.html


### Forest

% vegetated hectares that are conifer dominated

#### Forest is compared in the following ways:

| x | y |
| --- | --- |

*(not in use)*

#### Topics that are influenced by Forest:

- Forest
  - Conifer cover
    - Forest cover
      - Upland vegetation health
        - LTW resilience final

[ untitled0 ]
