## Appendix 3 for "Promoting long-term forest landscape resilience in the Lake Tahoe basin": Hardwood.html


### Hardwood

% vegetated hectares that are aspen dominated

#### Topics that are influenced by Hardwood:

- Hardwood
  - Hardwood cover
    - Forest cover
      - Upland vegetation health
        - LTW resilience final

[ untitled0 ]
