## Appendix 3 for "Promoting long-term forest landscape resilience in the Lake Tahoe basin": Hardwood_cover.html


### Hardwood cover

#### dependency network

#### Topics that are influenced by Hardwood cover:

- Hardwood cover
  - Forest cover
    - Upland vegetation health
      - LTW resilience final

#### Topics that influence Hardwood cover:

- Hardwood cover
  - Hardwood1
  - HardwoodI1
  - Hardwood2
  - HardwoodI2
  - Hardwood3
  - HardwoodI3
  - Hardwood4
  - HardwoodI4
  - Hardwood

[ untitled0 ]
