## Appendix 3 for "Promoting long-term forest landscape resilience in the Lake Tahoe basin": HCKS_aspen_habitat.html


### HCKS aspen habitat

#### dependency network

#### Topics that are influenced by HCKS aspen habitat:

- HCKS aspen habitat
  - Habitat for cultural keystone species
    - Cultural resource quality
      - LTW resilience final

#### Topics that influence HCKS aspen habitat:

- HCKS aspen habitat
  - CultASP1
  - CultASPI1
  - CultASP2
  - CultASPI2
  - CultASP3
  - CultASPI3
  - CultASP4
  - CultASPI4
  - CultASP

[ untitled0 ]
