## Appendix 3 for "Promoting long-term forest landscape resilience in the Lake Tahoe basin": HCKS_flicker_habitat.html


### HCKS flicker habitat

#### dependency network

#### Topics that are influenced by HCKS flicker habitat:

- HCKS flicker habitat
  - Habitat for cultural keystone species
    - Cultural resource quality
      - LTW resilience final

#### Topics that influence HCKS flicker habitat:

- HCKS flicker habitat
  - Flicker1
  - FlickerI1
  - Flicker2
  - FlickerI2
  - Flicker3
  - FlickerI3
  - Flicker

[ untitled0 ]
