## Appendix 3 for "Promoting long-term forest landscape resilience in the Lake Tahoe basin": HCKS_mountain_quail_habitat.html


### HCKS mountain quail habitat

#### dependency network

#### Topics that are influenced by HCKS mountain quail habitat:

- HCKS mountain quail habitat
  - Habitat for cultural keystone species
    - Cultural resource quality
      - LTW resilience final

#### Topics that influence HCKS mountain quail habitat:

- HCKS mountain quail habitat
  - MtnQuail1
  - MtnQuailI1
  - MtnQuail2
  - MtnQuailI2
  - MtnQuail3
  - MtnQuailI3
  - MtnQuail

[ untitled0 ]
