## Appendix 3 for "Promoting long-term forest landscape resilience in the Lake Tahoe basin": HCKS_mule_deer_habitat.html


### HCKS mule deer habitat

#### dependency network

#### Topics that are influenced by HCKS mule deer habitat:

- HCKS mule deer habitat
  - Habitat for cultural keystone species
    - Cultural resource quality
      - LTW resilience final

#### Topics that influence HCKS mule deer habitat:

- HCKS mule deer habitat
  - MuleDeer1
  - MuleDeerI1
  - MuleDeer2
  - MuleDeerI2
  - MuleDeer3
  - MuleDeerI3
  - MuleDeer

[ untitled0 ]
