## Appendix 3 for "Promoting long-term forest landscape resilience in the Lake Tahoe basin": Herbivore.html


### Herbivore

### herbivores in which 70% of their 2010 habitat is maintained/# herbivores

#### Topics that are influenced by Herbivore:

- Herbivore
  - Herbivore funct redund
    - Ecological function
      - Wildlife conservation
        - LTW resilience final

[ untitled0 ]
