## Appendix 3 for "Promoting long-term forest landscape resilience in the Lake Tahoe basin": Herbivore_funct_redund.html


### Herbivore funct redund

#### dependency network

#### Topics that are influenced by Herbivore funct redund:

- Herbivore funct redund
  - Ecological function
    - Wildlife conservation
      - LTW resilience final

#### Topics that influence Herbivore funct redund:

- Herbivore funct redund
  - Herbivore1
  - HerbivoreI1
  - Herbivore2
  - HerbivoreI2
  - Herbivore

[ untitled0 ]
