## Appendix 3 for "Promoting long-term forest landscape resilience in the Lake Tahoe basin": HerbivoreI1.html


### HerbivoreI1

#### Topics that are influenced by HerbivoreI1:

- HerbivoreI1
  - Herbivore funct redund
    - Ecological function
      - Wildlife conservation
        - LTW resilience final

[ untitled0 ]
