## Appendix 3 for "Promoting long-term forest landscape resilience in the Lake Tahoe basin": HerbivoreI2.html


### HerbivoreI2

#### Topics that are influenced by HerbivoreI2:

- HerbivoreI2
  - Herbivore funct redund
    - Ecological function
      - Wildlife conservation
        - LTW resilience final

[ untitled0 ]
