## Appendix 3 for "Promoting long-term forest landscape resilience in the Lake Tahoe basin": High_elevation.html


### High elevation

#### dependency network

#### Topics that are influenced by High elevation:

- High elevation
  - Seral stage
    - Upland vegetation health
      - LTW resilience final

#### Topics that influence High elevation:

- High elevation
  - High elevation early
    - HighEarly1
    - HighEarlyI1
    - HighEarly2
    - HighEarlyI2
    - HighEarly3
    - HighEarlyI3
    - HighEarly4
    - HighEarlyI4
    - HighEarly5
    - HighEarlyI5
    - HighEarly6
    - HighEarlyI6
    - HighEarly
  - High elevation mid
    - HighMid1
    - HighMidI1
    - HighMid2
    - HighMidI2
    - HighMid3
    - HighMidI3
    - HighMid4
    - HighMidI4
    - HighMid5
    - HighMidI5
    - HighMid6
    - HighMidI6
    - HighMid
  - High elevation late
    - HighLate1
    - HighLateI1
    - HighLate2
    - HighLateI2
    - HighLate3
    - HighLateI3
    - HighLate4
    - HighLateI4
    - HighLate5
    - HighLateI5
    - HighLate6
    - HighLateI6
    - HighLate

[ untitled0 ]
