## Appendix 3 for "Promoting long-term forest landscape resilience in the Lake Tahoe basin": High_elevation_late.html


### High elevation late

#### dependency network

#### Topics that are influenced by High elevation late:

- High elevation late
  - High elevation
    - Seral stage
      - Upland vegetation health
        - LTW resilience final

#### Topics that influence High elevation late:

- High elevation late
  - HighLate1
  - HighLateI1
  - HighLate2
  - HighLateI2
  - HighLate3
  - HighLateI3
  - HighLate4
  - HighLateI4
  - HighLate5
  - HighLateI5
  - HighLate6
  - HighLateI6
  - HighLate

[ untitled0 ]
