## Appendix 3 for "Promoting long-term forest landscape resilience in the Lake Tahoe basin": High_elevation_mid.html


### High elevation mid

#### dependency network

#### Topics that are influenced by High elevation mid:

- High elevation mid
  - High elevation
    - Seral stage
      - Upland vegetation health
        - LTW resilience final

#### Topics that influence High elevation mid:

- High elevation mid
  - HighMid1
  - HighMidI1
  - HighMid2
  - HighMidI2
  - HighMid3
  - HighMidI3
  - HighMid4
  - HighMidI4
  - HighMid5
  - HighMidI5
  - HighMid6
  - HighMidI6
  - HighMid

[ untitled0 ]
