## Supplementary figures and images for "Promoting long-term forest landscape resilience in the Lake Tahoe basin"

### _ante.png

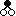

### _args.png

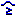

### _calc.png

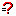

### _carg.png

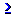

### _choice.png

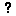

### _choices.png

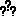

### _constant.png

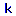

### _db.png

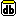

### _dblink.png

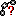

### _dep.png

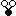

### _doc.png

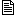

### _docs.png

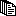

### _farg.png

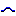

### _goal.png

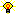

### _group.png

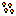

### _hlink.png

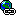

### _id.png

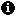

### _map.png

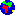

### _outline.png

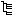

### _places.png

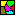

### _properties.png

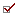

### _stats.png

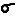

### _var.png

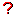

### Apex_predators.net.png

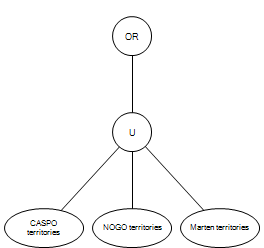

### asc.gif

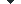

### Aspen_forest.net.png

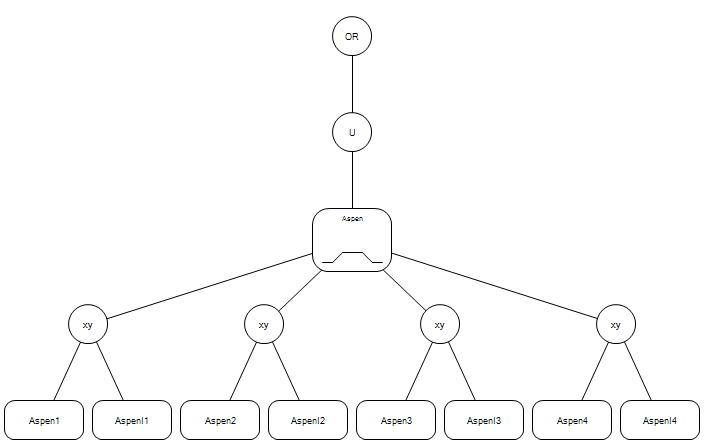

### Aspen_habitat.net.png

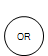

### bg.gif

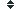

### Big_trees.net.png

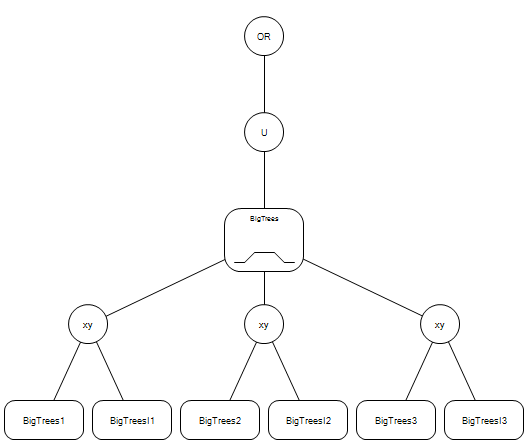

### BigTrees.farg0.png

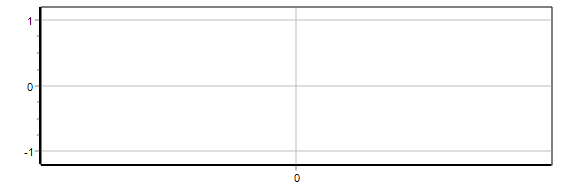

### Burned_high_sev.net.png

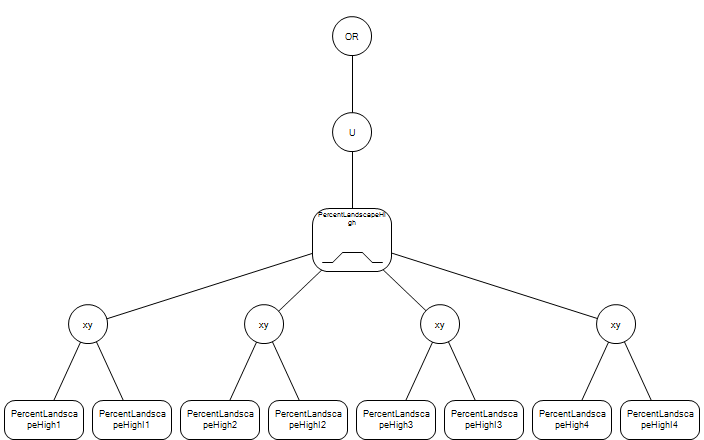

### Burned_low_sev.net.png

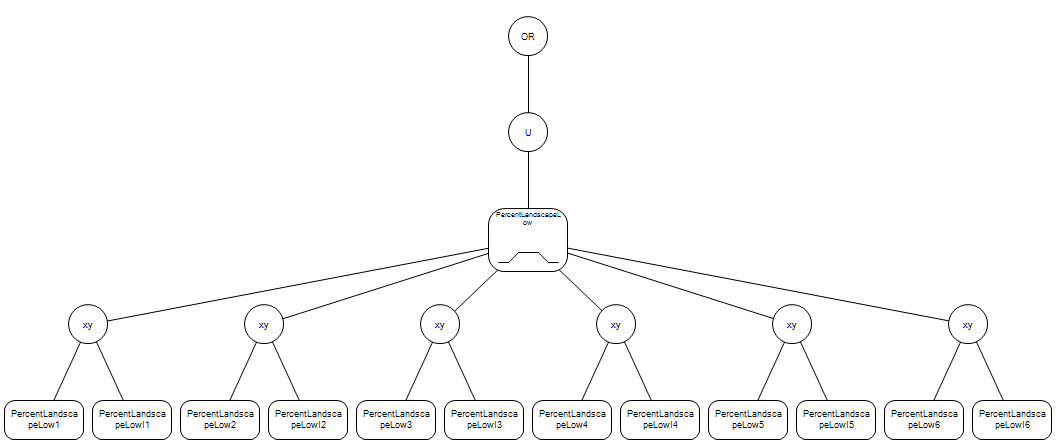

### Burned_mod_sev.net.png

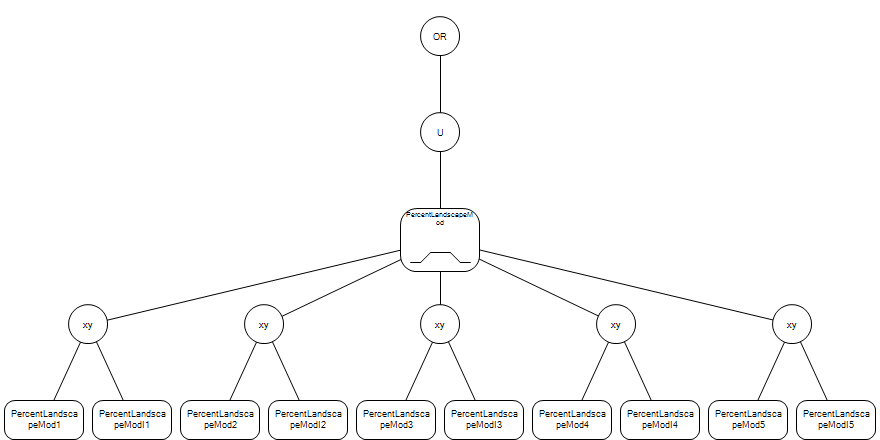

### CASPO_territories.net.png

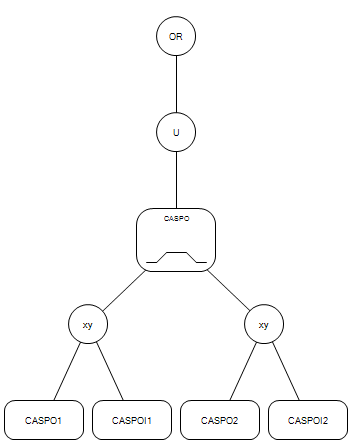

### Composition.net.png

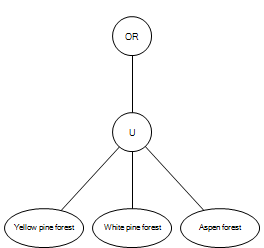

### Conifer_cover.net.png

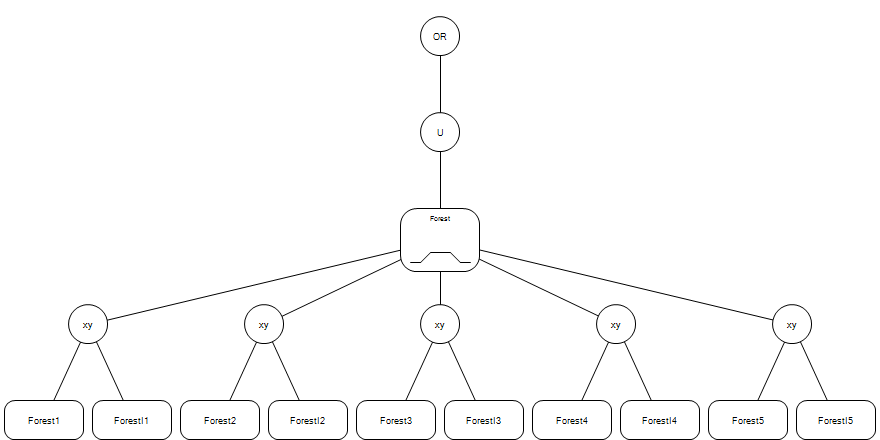

### Cult_water_quant_and_timing.net.png

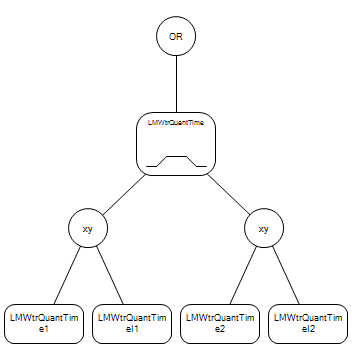

### Cultural_resource_quality.net.png

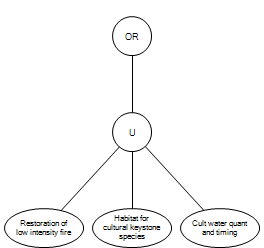

### Decomp_funct_redund.net.png

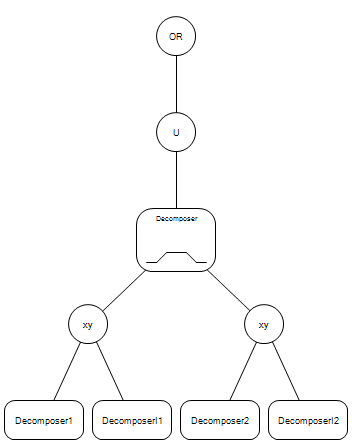

### Defense_zone_fire_severity.net.png

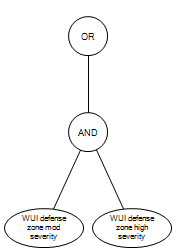

### desc.gif

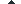

### Early_seral_beta_div.net.png

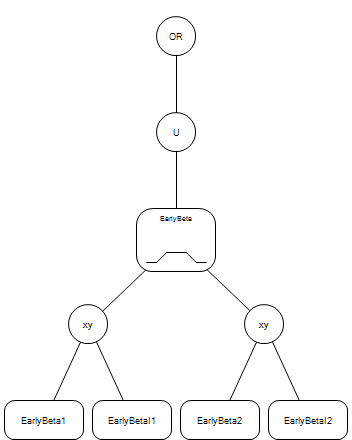

### Ecological_function.net.png

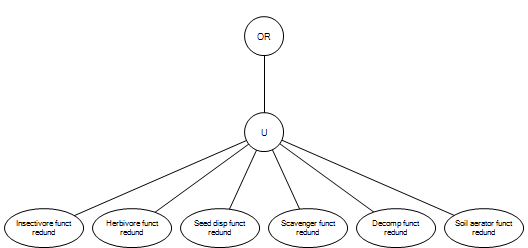

### Fine_sediment_load.net.png

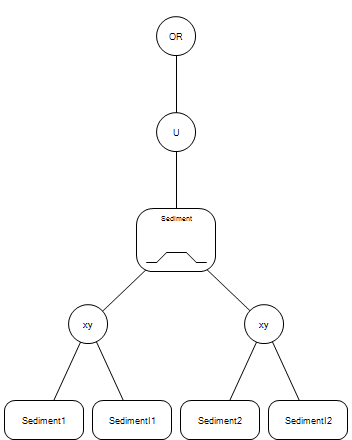

### Forest_cover.net.png

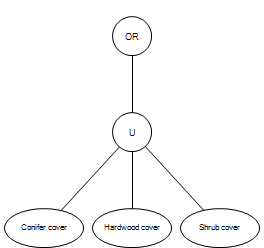

### Functional_fire.net.png

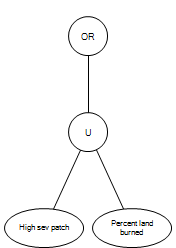

### g_colorbar.jpg

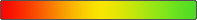

### g_colorbar_red_black_green.jpg

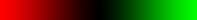

### g_marker.gif

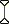

### Habitat_for_cultural_keystone_species.net.png

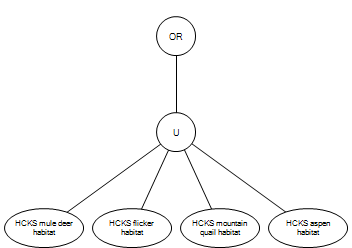

### Hardwood_cover.net.png

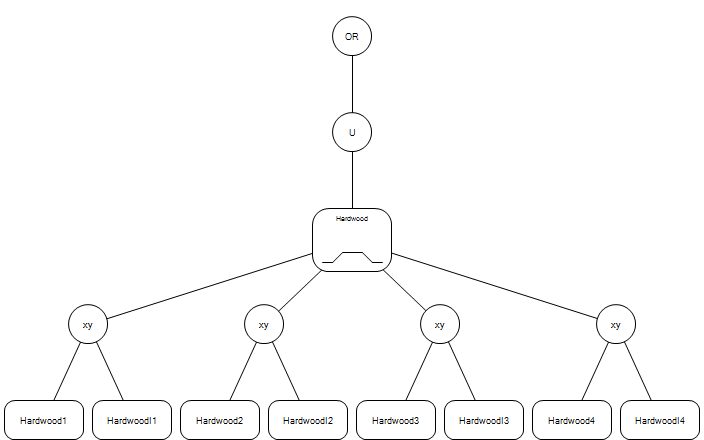

### HCKS_aspen_habitat.net.png

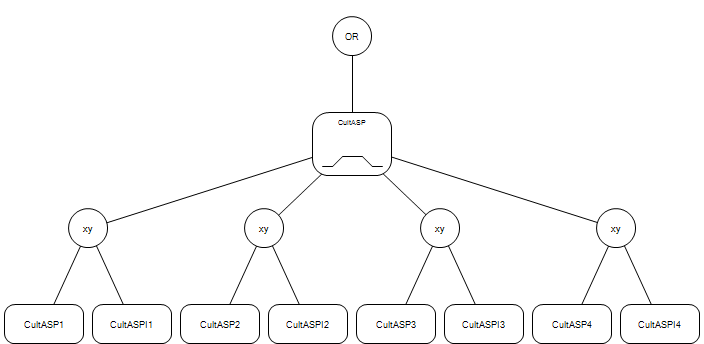

### HCKS_flicker_habitat.net.png

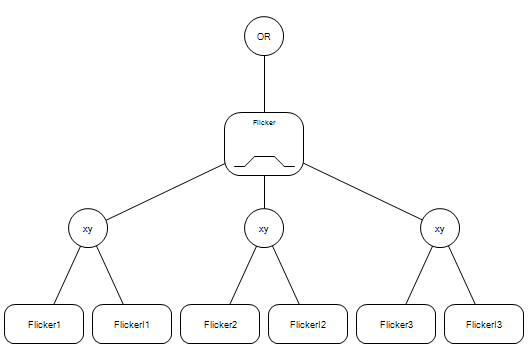

### HCKS_mountain_quail_habitat.net.png

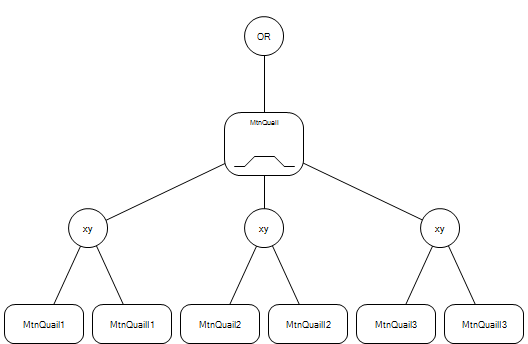

### HCKS_mule_deer_habitat.net.png

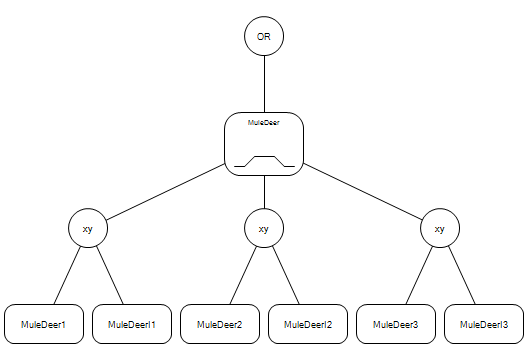

### Herbivore_funct_redund.net.png

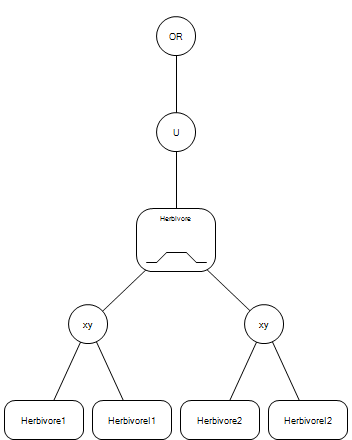

### High_elevation.net.png

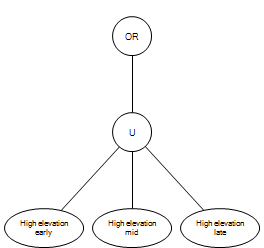

### High_elevation_early.net.png

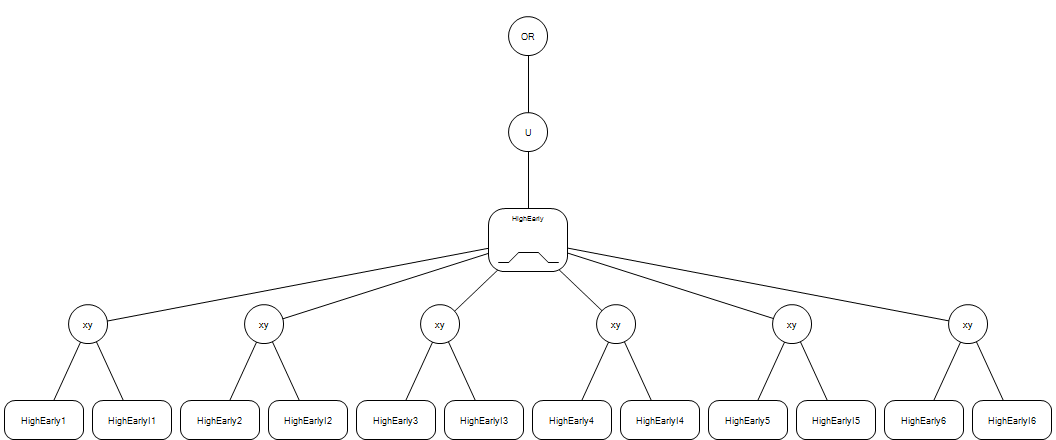

### High_elevation_late.net.png

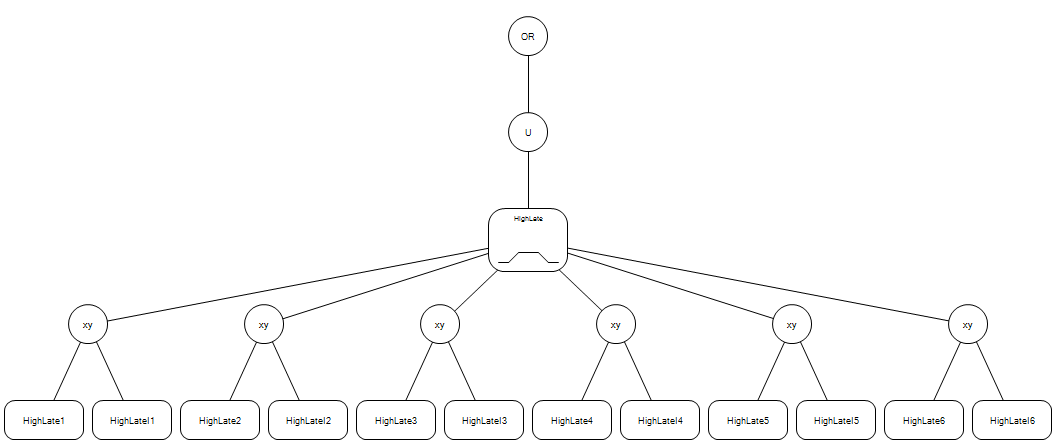

### minus.gif

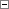

### plus.gif

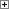

### treeview-line.gif

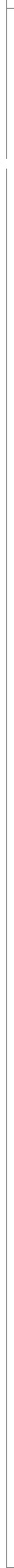

### treeview.gif

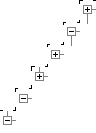
